# A Molecular Glue that Regulates Processing of Non-Coding RNAs in Human Cells

**DOI:** 10.64898/2026.11.14.766697

**Authors:** Haoxin Li, Luca F.R. Gebert, Tiantai Ma, Keegan Flanagan, Jason Tse, Gene W. Yeo, Jens Lykke-Andersen, Bruno Melillo, Ian J. MacRae, Benjamin F. Cravatt

## Abstract

Small nuclear RNAs (snRNAs) are a major class of non-coding RNAs (ncRNAs) and essential components of the spliceosome. Chemical tools for perturbing snRNA structure and function are lacking, and ncRNAs have more generally been considered undruggable. Here we use chemical proteomics and cryo-electron microscopy to identify a covalent molecular glue HL-1B that stabilizes binding of the 3′-exonuclease TOE1 to the spliceosome in a catalytically competent state. We show that HL-1B triggers excess 3′-processing of snRNAs in human cells that is tunable by the expression levels of TOE1 and auxiliary splicing regulators like SNRPB2, which, in turn, create synthetic lethality interactions with HL-1B that correlate with the preferential modulation of intermediate exon inclusion splicing events. Our findings reveal a fundamental role for snRNA tail length in regulating spliceosome function and, more generally, a way to target ncRNAs with non-orthosteric small molecules that remodel the interactomes of RNA processing enzymes in cells.

## Introduction

Non-coding RNAs (ncRNAs) constitute a major portion of the human transcriptome and have emerged as central regulators of gene expression with diverse biogenesis mechanisms, sizes, shapes, and functions^1,2^. Small nuclear RNAs (snRNAs) are a class of ncRNAs that serve as core components of the spliceosome, a large and dynamic molecular machine composed of a set of small nuclear ribonucleoprotein (snRNP) complexes, each containing a seven-member Sm protein ring, additional snRNP-specific protein components, and snRNAs, that together choreograph splice-site definition, catalytic activity, and proofreading across human introns^3–8^.

snRNAs undergo multiple layers of post-transcriptional processing^9^, including 3′-end maturation by the DEDDh-family exonuclease TOE1 (target of EGR1 protein 1)^10–13^. Concentrated in Cajal bodies, TOE1 functions as a quality control factor that preferentially processes Sm ring-assembled snRNAs, thereby promoting their maturation and stability, which are essential for efficient early spliceosome assembly^10,11,14–16^. In addition to regulating snRNAs, TOE1 contributes to 3′-end processing of select small nucleolar RNAs (snoRNAs)^12^, small Cajal body-specific RNAs (scaRNAs)^14^ and the telomerase RNA (TERC)^17^. Biallelic loss-of-function mutations in TOE1 cause pontocerebellar hypoplasia (PCH7), and cells from PCH7 patients accumulate 3′-extended, insufficiently processed (under-trimmed) snRNAs^11^.

Genetic approaches for studying the spliceosome are technically challenged by the essentiality of its core snRNP components for cell growth. Small molecules that can acutely perturb spliceosome function have accordingly served as key research tools^18^. These compounds include natural products and derivatives thereof (e.g., pladienolide B (pladB), spliceostatins)^19,20^ and synthetic covalent ligands (e.g., WX-02-23)^21^ that bind the SF3B subcomplex of the U2 snRNP, as well as molecular glue degraders of the pre-mRNA splicing factor RBM39 (e.g., indisulam)^22–24^. However, small molecules targeting and/or degrading such essential splicing factors can trigger transcriptome-wide splicing alterations and blockade of general cell growth^21,25^. Chemical probes that perturb pathways responsible for the post-transcriptional regulation of the spliceosome may accordingly offer an attractive alternative for studying its structure and function without causing gross cellular toxicity.

Recent efforts by our lab to map ligandable and functional cysteine residues across the human proteome by integrated activity-based protein profiling (ABPP) and genetic base editing identified tryptoline acrylamide small molecules that stereoselectively reacted with a non-orthosteric and paralog-restricted cysteine (C80) in TOE1^26^. These compounds showed initial evidence of affecting TOE1 nuclease activity *in vitro*, but limitations in potency and selectivity precluded a more detailed understanding of their mechanism of action and cellular effects.

Here, we have used *in cellulo* ABPP-guided medicinal chemistry to identify an advanced TOE1 ligand – HL-1B – that exhibits > 30-fold improvements in potency and excellent proteome-wide selectivity. Further *in vitro* characterization of HL-1B revealed that the covalent ligand only reacts with TOE1 in the presence of snRNP complexes, and cryogenic electron microscopy (cryo-EM) localized HL-1B to a non-orthosteric composite pocket containing C80 of TOE1 and residues from the SNRPE and SNRPF subunits of the Sm ring. Notably, this non-orthosteric molecular glue interaction stabilizes the binding of TOE1 to snRNPs in a catalytically competent state, leading to an unprecedented biochemical outcome in cells – the precise over-trimming and removal of the 3′ stem loops of snRNAs. The HL-1B-induced tail remodeling of snRNAs produces discrete changes in splicing enriched for events with intermediate percent spliced in (PSI) values that sharply contrast with the transcriptome-wide perturbations caused by the direct spliceosome inhibitor pladienolide B (pladB)^19^. We further demonstrate that the magnitude of HL-1B effects on snRNA trimming, mRNA splicing, and cell proliferation can be modulated by the expression level of Sm regulatory factors, including TOE1 itself and the U2 snRNA-binding protein SNRPB2, leading to the discovery of synthetic lethality relationships in cancer cells. Finally, we show that HL-1B remains effective on a hypomorphic TOE1 background, where the compound removes 3′-extensions of the U1 snRNA that accumulate in cells expressing PCH7-associated TOE1 mutants. Our findings thus uncover an unexpected chemical mechanism for controlling snRNA tail homeostasis that enables tunable modulation of splicing without disrupting global spliceosome homeostasis.

## Results

### Optimization of a covalent small molecule targeting TOE1

The ligandable and functional cysteine (C80) in TOE1 assigned by integrated ABPP and base editing is located distal to the catalytic D64/E66 dyad based on AlphaFold protein structure prediction^26–28^ (**Figures 1A** and **S1**), suggesting that genetic mutations and small molecules targeting this site may act through a non-orthosteric mechanism rather than direct active site competition with RNA substrates. Consistent with this model, a first-generation covalent tryptoline acrylamide ligand WX-02-33 (IC_50_ = 20 μM; **Figures S2A, B**) modestly enhanced TOE1 binding to multiple snRNPs in the human cancer cell line MCC142^26^. To better understand the functional effects of covalent compounds targeting C80 of TOE1, we optimized the potency and selectivity of these ligands by ABPP-guided medicinal chemistry performed in MCC142 cells expressing recombinant WT-TOE1 (**Figures S2A–F**). These efforts revealed that independent replacement of the C1-appended 1,3-benzodioxole and C3-appended morpholine groups of WX-02-33 with fluorophenyl (WX-03-79) and 2-oxa-8-azaspiro[4.5]decane (WX-02-322) groups, respectively, improved TOE1 engagement in cells (**Figures S2A–F**), and combining both modifications yielded a compound HL-1B that stereoselectively engaged TOE1 with a > 30-fold improvement in potency (IC_50_ = 0.55 μM; **Figures 1B** and **S2A–F**).

**Figure 1.**
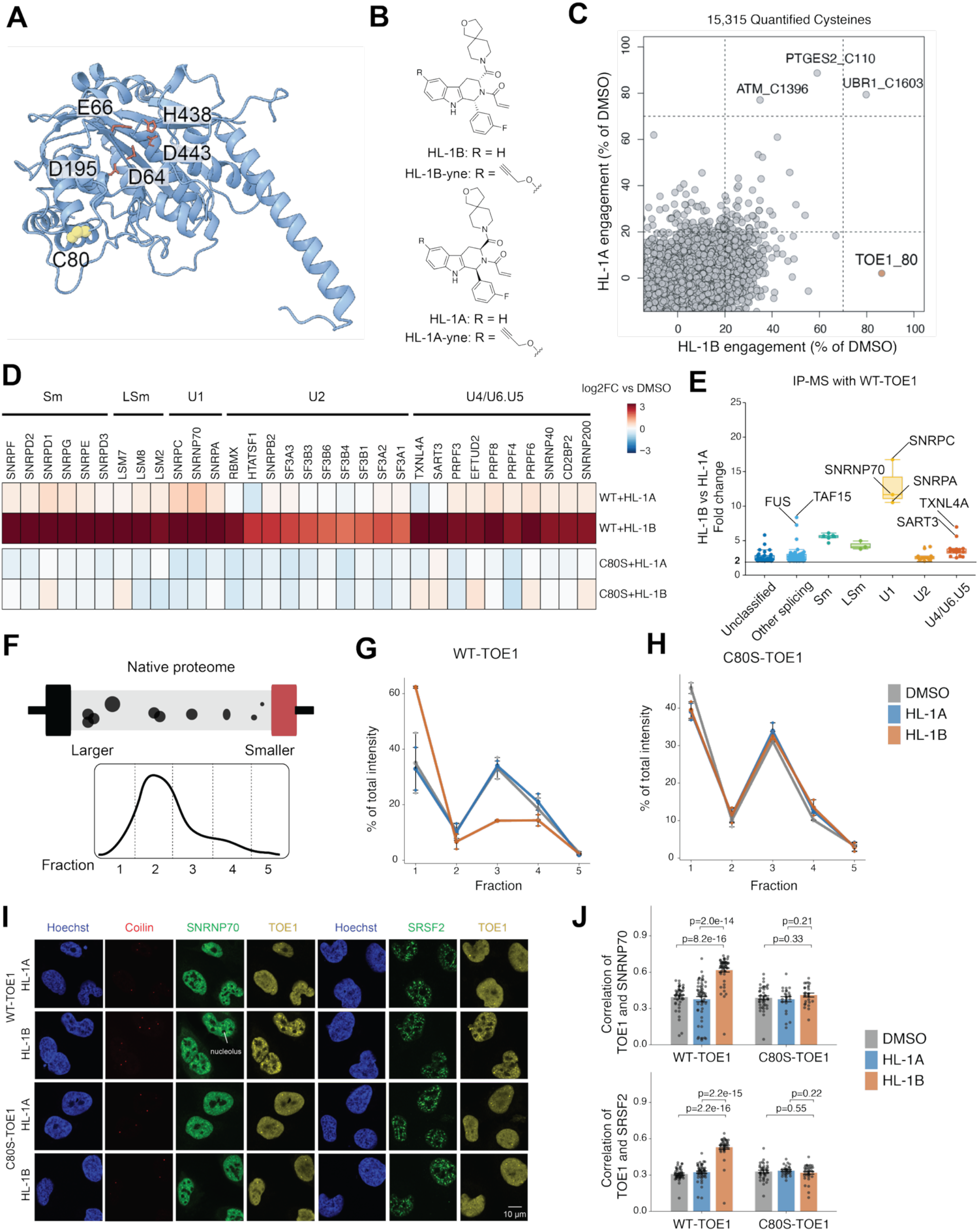
Discovery of a potent and selective covalent small molecule that stabilizes TOE1 binding to the spliceosome. (A) AlphaFold3 prediction of the TOE1 structure showing the tryptoline acrylamide-liganded cysteine (C80) and conserved catalytic residues (D64, E66, D195, H438, and D443). (B) Structures of non-alkyne and alkyne tryptoline acrylamide TOE1 ligands (HL-1B and HL-1B-yne) and inactive enantiomers (HL-1A and HL-1A-yne). (C) Scatter plot showing cysteine-directed ABPP data from parental MCC142 cells treated with HL-1B (x-axis) or HL-1A (y-axis) (5 μM, 4 h). Data are average values from 2 independent experiments. (D) Heatmap showing spliceosomal proteins (organized by spliceosomal sub-complexes) that were co-enriched in anti-FLAG IP-MS experiments from FLAG-tagged WT/C80S-TOE1-expressing MCC142 cells treated with HL-1B, HL-1A (5 μM, 8 h) or DMSO. Data are average values from 4-6 independent experiments. (E) Boxplot showing quantification of proteins co-enriched in anti-FLAG IP-MS experiments as described in (**D**). (F) Schematic showing relationship between size-exclusion chromatography fractions and protein solution size. (**G** and **H**) SEC-MS elution profiles of TOE1 from WT-TOE1-expressing (**G**) or C80S-TOE1-expressing (**H**) MCC142 cells treated with HL-1A, HL-1B (5 μM, 8 h) or DMSO. Clarified lysates from treated cells were separated by FPLC and fractionated into five fractions before MS analysis. Data are average values ± SEM from 2 independent experiments. (**I**) Effects of HL-1B versus HL-1A on the subcellular localization of TOE1. WT-TOE1- and C80S-TOE1-expressing MCC142 cells were treated with HL-1A or HL-1B (2 μM, 24 h) or DMSO before confocal imaging with antibodies for the indicated proteins. Data are representative of 4 independent fields. Scale bar, 10 μm. (**J**) Barplots of nuclear TOE1-SNRNP70 or TOE1-SRSF2 pixel intensity correlations as determined by CellProfiler. Data are average values +/-SEM from 4 independent fields and each dot represents 1 cell. P values were calculated using two-sided Student’s t test.

We next synthesized alkynylated analogues of HL-1B and its inactive enantiomer HL-1A (HL-1B-yne and HL-1A-yne) (**Figure 1B**) and confirmed by gel-ABPP that HL-1B-yne, but not HL-1A-yne, reacted with WT-TOE1, but not a C80S-TOE1 mutant in cells (**Figure S2G**). We then used HL-1B-yne in both gel- and mass spectrometry (protein-directed^29^)-ABPP experiments to verify the stereoselective engagement of recombinant and endogenous TOE1 by HL-1B in MCC142 cells (**Figures S2H, I** and **Table S1**). The protein-directed ABPP experiments further demonstrated that HL-1B (5 μM, 3 h) engaged TOE1 with excellent proteome-wide selectivity (**Figures S2I, J** and **Table S1**). Cysteine-directed ABPP performed as described previously using an iodoacetamide-desthiobiotin probe^29,30^ provided additional evidence for the selectivity of HL-1B, as across > 15,000 quantified cysteines, TOE1_C80 was the principal stereoselective target of this compound in MCC142 cells (∼90% engagement at 5 μM, 4 h; **Figure 1C** and **Table S1**).

Our ABPP-guided medicinal chemistry efforts thus established HL-1B as a potent, site-specific, and stereoselective covalent ligand for TOE1_C80 suitable for more in-depth biological studies in human cells.

### HL-1B stabilizes TOE1 interactions with snRNP complexes in cells

The increased engagement potency displayed by HL-1B in comparison to the first-generation compound WX-02-33 correlated with marked enhancement in the ligand-induced binding of recombinant Flag-tagged WT-TOE1 to spliceosome proteins in cancer cells as determined by immunoprecipitation-mass spectrometry (IP-MS). HL-1B (5 μM, 8 h) strengthened WT-TOE1 interactions with various snRNP proteins (**Figures 1D, E** and **Table S1**), including particularly pronounced effects with components of the U1 snRNP complex (e.g., SNRPC, SNRNP70, and SNRPA), which exhibited ∼20 times higher co-enrichment with TOE1 compared to control experiments performed with DMSO, the inactive enantiomer HL-1A, or a C80S-TOE1 mutant (**Figures 1D, E**). In contrast, WX-02-33 (20 μM, 8 h) only produced ∼2-3-fold increases in U1 snRNP protein interactions with TOE1 (**Figure S3A**). HL-1B also strengthened TOE1 interactions with components of the U4/U6•U5 tri-snRNP and U2-snRNP complexes, as well as the core Sm and LSm complexes, but to lesser magnitudes (∼3 to 5-fold) (**Figures 1D, E**). We verified that HL-1B stereoselectively enhanced snRNP protein interactions with recombinant WT-TOE1, but not C80S-TOE1, in additional cell lines (TC71 and HeLa S3; **Figures S3B, C** and **Table S1**), as well as with endogenous TOE1, as determined by IP-MS experiments performed in cell lines expressing epitope-tagged SNRPC (U1) and SNRNP40 (U5) proteins (**Figure S3D** and **Table S1**).

As a complementary approach to IP-MS, we treated WT-TOE1- and C80S-TOE1-expressing MCC142 cells with DMSO or HL-1B or HL-1A (5 μM, 8 h) and analyzed native proteomes from these cells by size-exclusion chromatography (SEC) and quantitative MS-based proteomics^21^ (**Figure 1F**). These experiments revealed that HL-1B, but not HL-1A, caused a pronounced redistribution of WT-TOE1 from a lower molecular weight SEC fraction (F3) to a higher molecular weight fraction (F1) (**Figure 1G** and **Table S1**) that also contained Sm core and snRNP-specific subunits (**Figures S3E-G**). The SEC profiles of these spliceosome proteins were generally unaltered by HL-1B, although a modest shift from F4 to F1 was also observed for the U1 snRNP protein SNRPC in HL-1B-treated cells (**Figure S3F**). In contrast, HL-1B did not alter the SEC profile of the C80S-TOE1 mutant or snRNP components in C80S-TOE1-expressing MCC142 cells (**Figures 1H, S3H-J** and **Table S1**).

Immunofluorescence microscopy experiments revealed that HL-1B also promoted the stereoselective co-localization of WT-TOE1, but not C80S-TOE1, with components of the U1 snRNP (SNRNP70) in speckle-like nuclear condensates (**Figures 1I, J**). Our combined IP-MS, SEC-MS, and microscopy results thus indicated that HL-1B induces the association of TOE1 with the U1 snRNP complex and related spliceosomal proteins in human cells.

### Structure of a TOE1–HL-1B–U1 snRNP complex

To better understand how HL-1B engages TOE1 and promotes interactions with snRNP complexes, we expressed and purified His-tagged WT- and C80S-TOE1 proteins from BL21(DE3) *E. coli* cells. Surprisingly, we found that purified WT-TOE1 did not react with HL-1B-yne (**Figure 2A**). We next purified the U1 snRNP complex from HeLa S3 cells using a 3′-desthiobiotinylated RNA oligo that captures the solvent-exposed 5′ end of the U1 snRNA^31,32^ and found that addition of this purified U1 snRNP complex restored robust stereoselective reactivity of WT-TOE1, but not C80S-TOE1, with HL-1B-yne (**Figure 2A**). A commercial source of snRNP complexes purified from bovine tissue consisting mostly of U1 snRNP also supported the stereoselective, site-selective, and concentration-dependent liganding of TOE1 by HL-1B(-yne) (**Figures 2B** and **S4**). These data demonstrated that tryptoline acrylamides do not react with TOE1 alone, but instead require the additional presence of the U1 snRNP complex.

**Figure 2.**
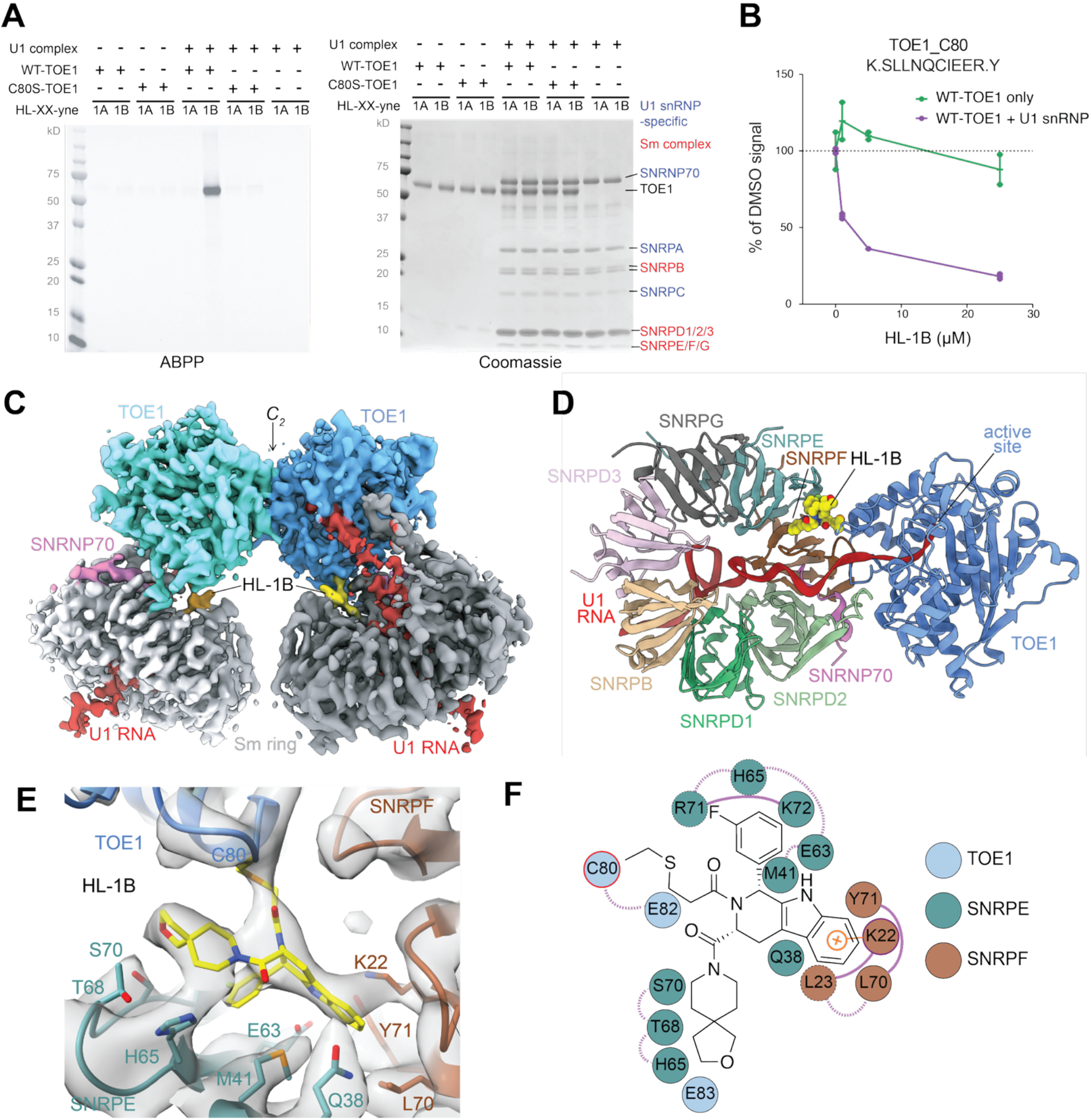
Structural characterization of a TOE1–HL-1B–U1 snRNP complex. (A) Left, gel-ABPP data showing the stereoselective and site-selective engagement of purified His-WT-TOE1 by HL-1B-yne (2 μM, 1 h) in the presence of a purified human U1 snRNP complex using the indicated alkyne stereoprobes. Right, Coomassie blue protein staining. Data are representative of 3 independent experiments. (B) Quantification of HL-1B engagement of TOE1_C80 in the presence or absence of the U1 snRNP complex. WT-TOE1 and bovine U1 snRNP were mixed with HL-1B at indicated concentrations for 1 h at room temperature, after which the samples were denatured, trypsin-digested, and analyzed by MS-based proteomics (see Methods), where HL-1B engagement was quantified by measuring the C80-containing tryptic peptide from TOE1. Data are average values ± SEM normalized to DMSO from 2 independent experiments. (C) Cryo-EM reconstruction of the TOE1–HL-1B–U1 snRNP assembly, comprising two TOE1 molecules, each bound to a separate U1 snRNP. The TOE1 dimer is organized around a twofold symmetry axis (C_2_). TOE1 molecules are shown in blue and cyan, Sm rings in gray, U1 snRNA in red, SNRNP70 in pink, and HL-1B in yellow. (D) Atomic model of one TOE1–HL-1B–U1 snRNP complex, showing the seven Sm ring proteins, U1 snRNA, SNRNP70, and TOE1. HL-1B (yellow spheres) binds a non-orthosteric pocket at the TOE1–Sm ring interface, spatially distinct from the TOE1 catalytic site, into which the U1 snRNA extends from the Sm ring. (E) Close-up view of HL-1B (yellow sticks) covalently bound to C80 of TOE1, fitted into the focused cryo-EM density map. Select residues of TOE1, SNRPE, and SNRPF surrounding the ligand are indicated. (F) Schematic representation of interactions between HL-1B and residues of TOE1, SNRPE, and SNRPF. The covalent linkage to TOE1_C80 and the proposed cation–π interaction with SNRPF_K22 are highlighted.

We next sought to elucidate how the U1 snRNP potentiates the reaction of HL-1B with TOE1 by determining the cryo-EM structure of the TOE1–HL-1B–U1 snRNP complex (**Figures S5–S7** and **Table S2**). For this structural analysis, we co-incubated purified recombinant human TOE1 and bovine U1 snRNP with HL-1B (see Methods). The cryo-EM structure revealed an assembly of two TOE1 molecules forming a dimer stabilized by a hydrophobic interface, with each TOE1 subunit engaging the Sm ring of a separate snRNP (**Figures 2C, D**, **Movie S1** and **S8A-C**). The structure resolved both a segment of the U1-specific protein SNRNP70 and snRNA density consistent with a U1-specific nucleotide (**Figures 2C, D** and **S8D, E**), supporting the identity of the complex as TOE1 bound to the U1 snRNP. A focus-refined map at 3.77 Å resolution contained well-defined density for HL-1B at the TOE1–Sm ring interface, including continuous density indicative of a covalent bond between the reactive carbon of the small molecule and the sulfur atom of C80 of TOE1 (**Figure 2E**). Remarkably, apart from covalent linkage to C80 and contacts with the adjacent E82 and E83 of TOE1, HL-1B interacts primarily with a pocket formed by the SNRPE and SNRPF components of the Sm ring that involves extensive van der Waals contacts and a cation–π interaction between the tryptoline core of HL-1B and K22 of SNRPF (**Figures 2D-F**). Thus, the structure explains the unusual snRNP dependence of HL-1B reactivity with TOE1: while HL-1B covalently modifies C80 of TOE1, the compound mostly binds to the Sm ring of the U1 snRNP complex.

Beyond explaining the basis for HL-1B reactivity, the cryo-EM structure also revealed how the compound stabilizes the TOE1–U1 snRNP complex. HL-1B inserts into a narrow crevice between SNRPE and SNRPF, acting as a molecular piton that anchors TOE1 to the surface of the Sm ring (**Figure 3A**). Comparison of the experimental structure with an otherwise identical model in which HL-1B was computationally removed revealed that the ligand contributes 335 Å^2^ of buried surface area to the TOE1–Sm ring interface, increasing the interface area by 41% beyond TOE1-Sm ring interactions alone (**Figure 3B**). These findings support a model in which HL-1B acts as an obligate molecular glue that binds a composite pocket formed by TOE1 and the Sm ring to capture and stabilize the otherwise transient interaction between these proteins.

**Figure 3.**
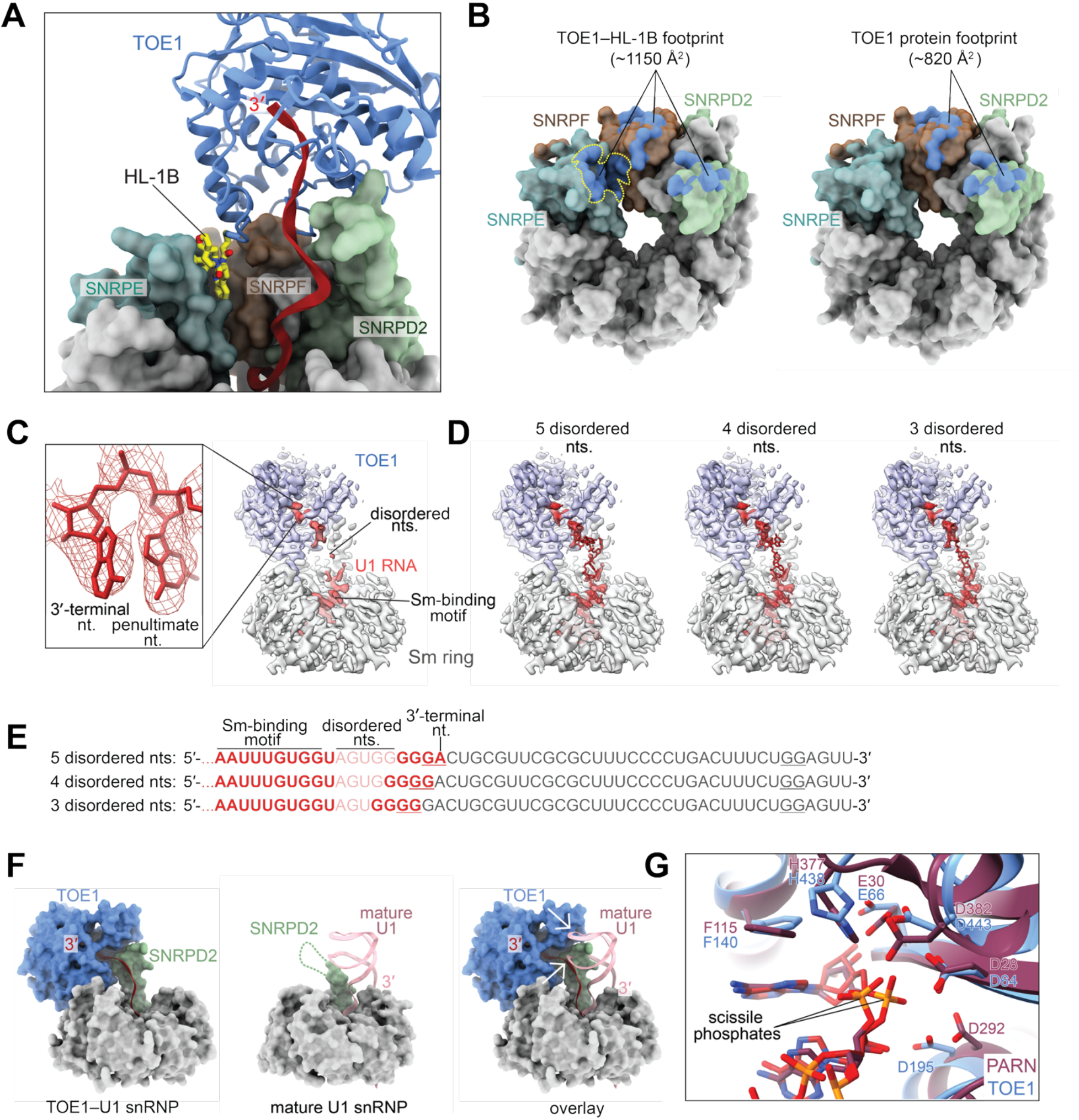
HL-1B stabilizes a catalytically competent state of the TOE1–HL-1B–U1 snRNP complex. (A) Structural model showing HL-1B (yellow sticks) covalently bound to TOE1 (blue cartoon) and inserted into a surface crevice between SNRPE and SNRPF, anchoring TOE1 to the Sm ring. Selected Sm proteins are shown as colored surfaces, and U1 snRNA is shown in red. **(B)** Surface representations of the Sm ring from the TOE1–HL-1B–U1 snRNP structure, with TOE1 removed to reveal the buried surface interface (blue). The left panel shows the surface buried by TOE1–HL-1B, while the right panel shows the surface buried by TOE1 protein alone for comparison. The HL-1B footprint is outlined in yellow. (C) Cryo-EM density of U1 snRNA extending from the Sm ring into the TOE1 active site. Density is well defined for the Sm-binding region and RNA 3′ terminus, but discontinuous in the intervening region, consistent with conformational disorder. The inset shows a close-up view of the 3′-terminal and penultimate nucleotides modeled into the cryo-EM density. TOE1, Sm ring, and U1 snRNA densities are shown in light blue, gray, and red, respectively. (D) Alternative RNA models containing five, four, or three nucleotides spanning the disordered region. At least three nucleotides are required to maintain backbone connectivity between the well-resolved RNA segments, but the precise sequence register could not be established from the density. (E) U1 snRNA sequences corresponding to the models in (D), with modeled nucleotides highlighted in red. The density for the two 3′-terminal nucleotides shown in (C) is consistent with two purines (A or G), allowing multiple possible sequence assignments within the GGGGGA stretch. Underlined dinucleotides indicate the possible assignments, whereas the next downstream purine pairs (underlined in gray) would require a substantially longer disordered segment. **(F)** Structural comparison of the TOE1–U1 snRNP complex (left), mature U1 snRNP (middle; PDB 7B0Y), and superposition of the mature U1 RNA onto the TOE1–U1 snRNP complex (right). TOE1 is blue, SNRPD2 is light green, and U1 snRNA is red (TOE1-bound) or pink (mature). The SNRPD2 loop that binds TOE1 is disordered in the mature U1 snRNP (dashed line). White arrows indicate steric clashes between the mature U1 snRNA stem-loop and both TOE1 and the TOE1-binding SNRPD2 loop. (G) Superposition of the TOE1 (blue) and PARN (burgundy; PDB 2A1R) active sites, highlighting the similar arrangement of catalytic residues and bound RNA substrates. Labeled side chains correspond to conserved active site residues.

The cryo-EM structure displayed an unexpected conformation for the U1 snRNA within the TOE1–HL-1B–U1 snRNP complex (**Figures 3C, D**). Instead of the intact stem-loop 4 (SL4) duplex found in previous structures of the mature U1 snRNP^31–33^, we observed density consistent with approximately 8–10 nucleotides of single-stranded RNA extending from the Sm ring into the TOE1 active site. The length and sequence register of this RNA indicate that its 3′ end is substantially shorter than that of the canonical mature U1 snRNA (**Figures 3C-E** and **S9**). The single-stranded RNA is contacted by a loop representing residues 77–89 of SNRPD2 that also engages TOE1 (**Figures 3A, F**). Notably, this loop is disordered in previous structures of mature U1 snRNP^33^, and its conformation in the TOE1-bound complex is sterically incompatible with binding to an SL4 duplex (**Figures 3A, F**). The 3′ end of the truncated U1 snRNA is positioned in the TOE1 active site near catalytic residues D64, E66, D195, H438, and D443, which adopt an arrangement that closely resembles the active site of the related DEDDh exonuclease PARN bound to an RNA substrate^34^ (**Figure 3G**).

We interpret the unique features of the TOE1–HL-1B–U1 snRNP structure to indicate that HL-1B acts as a non-orthosteric molecular glue to stabilize the TOE1–U1 snRNP complex in a catalytically competent state that may exist beyond its normal lifespan in cells, thereby leading to excessive processing of the 3′ end of the U1 snRNA.

### HL-1B promotes over-trimming of snRNAs in cells

Our biochemical and structural findings raised the possibility that HL-1B might alter snRNA maturation in cells, which we proceeded to test by snRNA-specific 3′-end sequencing^11,14,15^ (**Figure 4A** and **Table S3**). We first confirmed that CRISPRi-mediated knockdown of endogenous TOE1 in NCIH520 cells (a high TOE1-expressing cell line^35^) resulted in decreased (under-) trimming of the 3′-tails of U1, U2, U4, and U5 snRNAs but not the non-substrate U3 snoRNA (**Figures 4B** and **S10A, B**), consistent with previous reports of impaired snRNA tail trimming in the absence of TOE1^11,14^. Conversely, stable expression of WT-TOE1, but not the catalytically inactive D64A/E66A double mutant, in MCC142 cells (a lower TOE1-expressing cell line^35^), resulted in modest increases in trimming (over-trimming) of U1 and U5 3′ ends with minimal effects on other RNAs (**Figures 4C** and **S10C**). These results linked TOE1 expression level and catalytic activity to the extent of processing of the 3′ termini of snRNAs.

**Figure 4.**
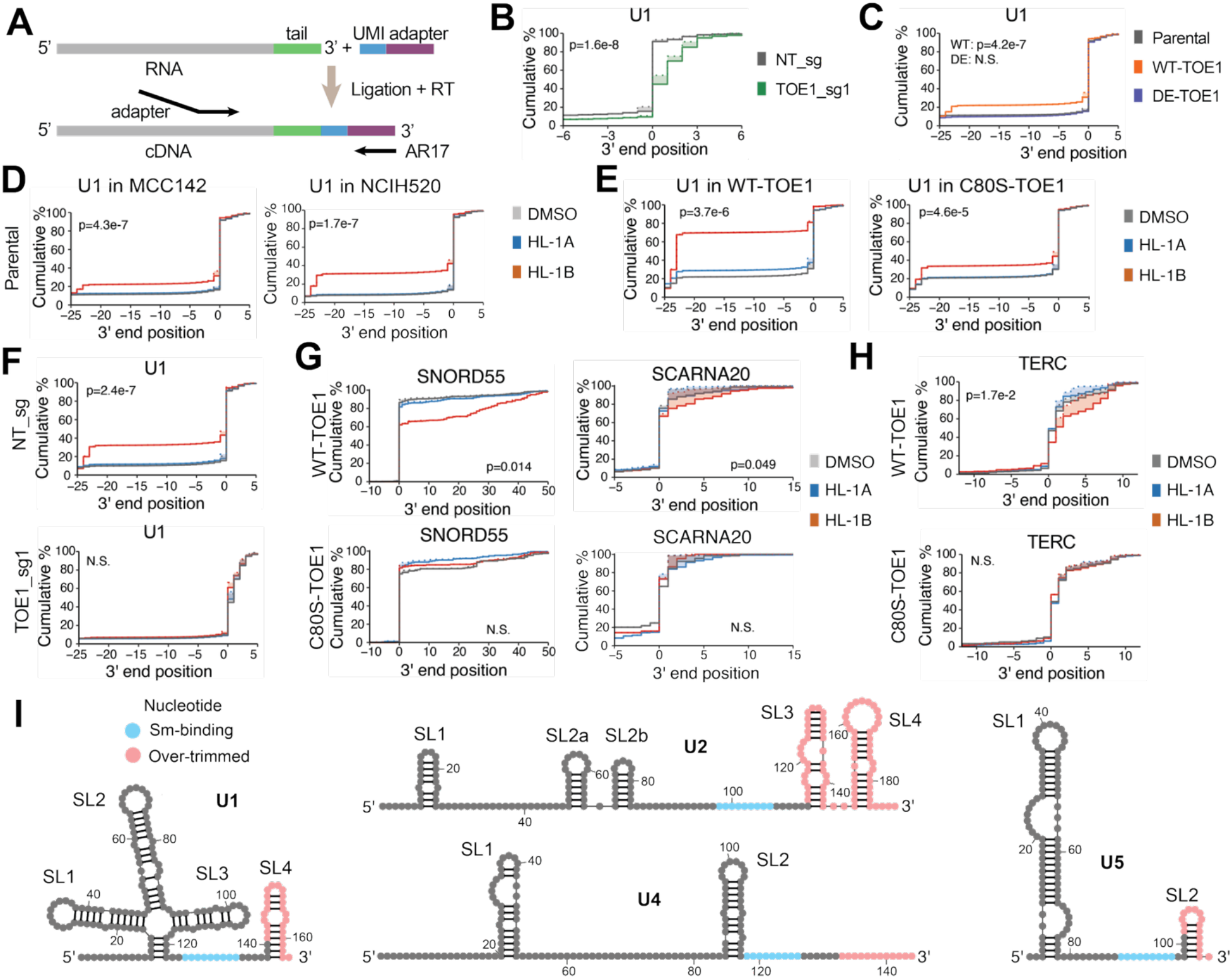
HL-1B promotes TOE1-mediated over-trimming of snRNAs in cells. (A) Workflow for gene-specific 3′ end sequencing, where RNA is extracted from cells, dephosphorylated, and adaptors containing Unique Molecular Identifier (UMI) are ligated to the 3′ end. After reverse transcription (RT), the AR17 primer and gene-specific primers are used to amplify the 3′ region of RNA including post-transcriptionally added tails, followed by indexing PCR and sequencing. (B) Cumulative plots comparing the effects of genetic (CRISPRi) knockdown of TOE1 in NCIH520 cells (TOE1_sg1) versus control NCIH520 cells (treated with a non-targeting sgRNA, NT_sg) on 3′ end position distributions of the U1 snRNA. In this and subsequent cumulative plots, mature 3′ ends of RNAs are represented as position 0 with positive numbers representing under-trimmed 3′ end tails and negative numbers representing over-trimmed 3′ end tails. Solid lines represent actual snRNA 3′ end positions, while dotted lines represent genome-encoded 3′ end positions with shading in between representing post-transcriptionally added nucleotides. (C) Cumulative plots comparing 3′ end position distributions of the U1 snRNA in parental MCC142 cells versus WT-expressing- or DE-TOE1 mutant-expressing MCC142 cells. DE, D64A+E66A, catalytically inactive TOE1 mutant. (D) Cumulative plots comparing 3′ end position distributions of the U1 snRNA in parental MCC142 (left) or NCIH520 (right) cells treated with HL-1A or HL-1B (2 μM, 3 days) or DMSO. (E) Cumulative plots comparing 3′ end position distributions of the U1 snRNA in WT-TOE1- and C80S-TOE1-expressing MCC142 cells treated with HL-1A or HL-1B (2 μM, 3 days) or DMSO. (F) Cumulative plots comparing 3′ end position distributions of the U1 snRNA in TOE1_sg1 and control NT_sg NCIH520 cells treated with HL-1A or HL-1B (2 μM, 3 days) or DMSO. (G) Cumulative plots comparing 3′ end position distributions of SNORD55 (snoRNA) or SCARNA20 (scaRNA) in WT-TOE1- and C80S-TOE1-expressing MCC142 cells treated with HL-1A or HL-1B (2 μM, 3 days) or DMSO. (H) Cumulative plots comparing 3′ end position distributions of TERC in WT-TOE1- and C80S-TOE1-expressing MCC142 cells treated with HL-1A or HL-1B (2 μM, 3 days) or DMSO. (I) Schematic of the positional relationship between over-trimmed snRNA stem loops (SLs) and the adjacent Sm-binding motif. For **B-H**, data are average values from 3 independent experiments of 3′ end sequencing (**B-F**, gene-specific data; **G**, global profiling data; **H**, nascent data). For **B**, the p value indicates comparison between TOE1_sg1 and NT_sg groups. For **C**, the p values indicate comparison between WT-TOE1 or DE-TOE1-expressing vs parental groups. For **D-H**, p values indicate comparisons between HL-1B- vs HL-1A-treated groups. Not significant (N.S.) is defined as p>0.05 or mean difference < 1 nucleotide. All p values were calculated using two-sided Student’s t test. N.S., not significant, * p<0.05, **p<0.01, ***p<0.001.

We next found that HL-1B, but not HL-1A, promoted over-trimming of the U1 snRNA in both MCC142 and NCIH520 cells (**Figure 4D**), and this effect was further potentiated in MCC142 cells by stable expression of WT-TOE1, but not C80S-TOE1 (**Figure 4E**). The over-trimming effects of HL-1B were blocked by CRISPRi-mediated knockdown of endogenous TOE1 (**Figure 4F**). HL-1B also promoted the stereoselective and site-specific over-trimming of U2, U4, and U5 snRNAs (except U3 snoRNA), but these effects were less dramatic and most clearly observed in WT-TOE1-expressing MCC142 cells (**Figure S10D**). We finally found that HL-1B did not promote over-trimming of the U1 snRNA in cells expressing the catalytically inactive D64A/E66A-TOE1 mutant (**Figure S10E**), despite still stereoselectively reacting with C80 in this mutant protein and inducing its binding to snRNP complexes (**Figures S10F, G** and **Table S3**).

We additionally evaluated cells treated with HL-1B (2 μM, 3 days) by global 3′-end sequencing of small RNAs^14^ (isolated by size selection). Interestingly, this global analysis revealed that certain snoRNAs (e.g., SNORD55) and scaRNAs (e.g., SCARNA20, an established TOE1 substrate^14^), exhibited longer, under-trimmed 3′ tails in HL-1B-treated WT-TOE1-, but not C80S-TOE1-expressing cells (**Figure 4G**). The telomerase RNA component TERC—another established TOE1 substrate^17^—also displayed 3′ tail extensions in HL-1B-treated WT-TOE1 cells, as detected by nascent gene-specific 3′ end sequencing to account for the long half-life of the TERC RNA (**Figure 4H**). We interpret these data, along with our previous immunofluorescence microscopy studies (**Figures 1I, J**), to indicate that HL-1B redistributes the subcellular pools of TOE1 activity in cells by prolonging the residence time of this enzyme bound to snRNP complexes, resulting in the over-trimming of snRNAs and under-trimming of other non-spliceosome ncRNA substrates.

The over-trimming effects of HL-1B produced discrete termination points for each snRNA that ranged from ∼12 to 75 nucleotides upstream of the canonical 3′ ends but converged on a site ∼5-7 nucleotides downstream of the Sm-binding motif (**Figure 4I**). Integration of these results with the cryo-EM structure of the TOE1–HL-1B–U1 snRNP complex supports a model where the HL-1B-mediated interactions between TOE1 and the Sm complex create a steric boundary restricting excess 3′-exonuclease processivity at a fixed offset from the Sm-binding site on snRNAs ^31,32^. The structural impact of this discrete overtrimming varied across the snRNAs, but resulted in the removal of key stem loops, including, as predicted from the cryo-EM model, stem loop 4 (SL4) of the U1 snRNA (**Figure 4I**).

### HL-1B produces selective cell growth effects

snRNAs are integral components of the spliceosome, and the genetic or pharmacological disruption of RNA splicing can broadly impair cell growth^21,36^. To our surprise, across a panel of 12 cancer cell lines spanning 10 lineages, we found that HL-1B minimally suppressed the growth of most cells (<20% reduction over ∼6 days) with the exception of the squamous lung cancer cell line NCIH520, which showed a substantial (∼50%), stereoselective decrease in proliferation (**Figure 5A**). Interestingly, NCIH520 cells also showed the highest relative expression of TOE1 across the cancer cell line panel (**Figure 5A***, left*), whereas the respective TOE1 genetic dependency scores for these cell lines from the Cancer Dependency Map^36^ were not correlated with HL-1B sensitivity (**Figure 5A***, right*). The cell growth effects of HL-1B were thus consistent with a gain-of-function mechanism where the anti-proliferative activity of this compound depends on the amount of TOE1 in cells. In further support of this model, we found that CRISPRi knockdown of TOE1 protected NCIH520 cells from the growth effects of HL-1B (**Figures 5B** and **S11A**), and, conversely, ectopic expression of WT-TOE1, but not C80S-TOE1, sensitized MCC142 cells to HL-1B (**Figures 5C** and **S11B**).

**Figure 5.**
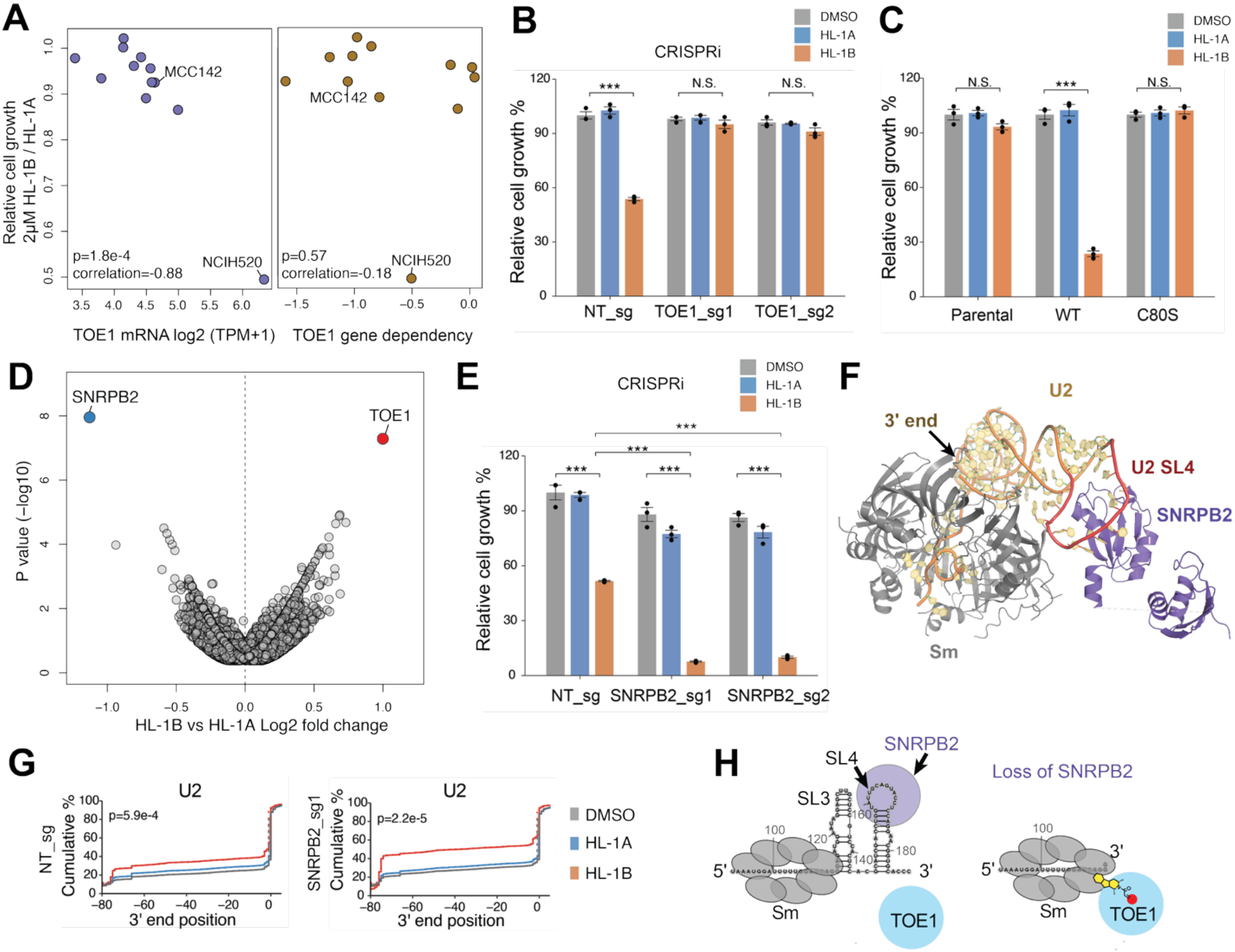
Genetic modifiers of the cancer cell growth effects of HL-1B. (A) Scatter plot showing a panel of 12 human cancer cell lines compared for: 1) the relative cell growth effects of HL-1B vs HL-1A (2 μM, 6 days); 2) the mRNA levels (left) or gene dependency scores (right) for TOE1. Growth effects were measured by CellTiter-Glo and are average values from 3 independent experiments. TOE1 mRNA and gene dependency data were extracted from the DepMap^35,36^. Pearson correlations and associated two-sided p values are shown. (B) Relative growth of NT_sg, TOE1_sg1/sg2 NCIH520-dCas9-KRAB cells treated with HL-1A or HL-1B (2 μM) or DMSO for 6 days as measured by CellTiter-Glo. Data are average values ± SEM from 3 independent experiments. (C) Relative growth of parental, WT-TOE1-expressing, and C80S-TOE1-expressing MCC142 cells treated with HL-1A or HL-1B (2 μM) or DMSO for 6 days as measured by CellTiter-Glo. Data are average values ± SEM from 3 independent experiments. (D) Volcano plots showing effects of gene knockdown on relative growth of HL-1B vs HL-1A-treated NCIH520 cells from the genome-wide CRISPRi screen. Sequencing data were collected at day 16 (10 days after treatment with HL-1A or HL-1B). (E) Relative growth of NT_sg, SNRPB2_sg1/sg2 NCIH520-dCas9-KRAB cells treated with HL-1A or HL-1B (2 μM) or DMSO for 6 days as measured by CellTiter-Glo. Data are average values ± SEM for 3 independent experiments. (F) Cryo-EM structure of human Sm complex containing a U2 snRNA fragment and associated SNRPB2 (PDB: 6Y5Q^38^). The black arrow marks the 3′ end of the U2 snRNA and Stem Loop 4 (SL4) is highlighted in red. (G) Cumulative plots comparing 3′ end position distributions of the U2 snRNA in NT_sg and SNRPB2_sg1 NCIH520 cells treated with HL-1A, HL-1B (2 μM, 3 days) or DMSO. Data are average values from 3 independent experiments and the p values indicate comparisons between HL-1B- vs HL-1A-treated groups. (H) Model for how SNRPB2 modulates the impact of HL-1B on TOE1-mediated trimming of U2 snRNA. SNRPB2 binds to the SL4 of U2 snRNA, which in turn suppresses the over-trimming of the U2 snRNA in HL-1B-treated cells. All p values were calculated using two-sided Student’s t test. N.S., not significant, * p<0.05, **p<0.01, ***p<0.001.

We next sought to uncover additional genetic modifiers of HL-1B activity in cells by performing a genome-wide CRISPRi screen in NCIH520 cells. Cells were spin-infected with the Dolcetto library^37^ (57,050 sgRNAs targeting 18,901 genes), selected with puromycin for 4 days, and treated with HL-1B or HL-1A (2 μM) for 10-20 additional days, after which sgRNAs were sequenced (**Figure S11C**). This experiment verified TOE1 as a major sensitizer for the antiproliferative activity of HL-1B (i.e., knockdown of TOE1 promoted resistance to HL-1B), whereas a single gene – SNRPB2 – scored highly as a resistance factor (i.e., knockdown of SNRPB2 sensitized cells to HL-1B) (**Figures 5D**, **S11D** and **Table S4**). We confirmed a synthetic lethal relationship between HL-1B and SNRPB2 using targeted CRISPRi, which, in control (DMSO- or HL-1A-treated) cells, produced only a modest (∼10-20%) growth defect, but, in HL-1B-treated cells, produced a near-complete blockade of cell proliferation (**Figures 5E** and **S11E**). SNRPB2, together with SNRPA1, forms the U2 snRNP heterodimer that specifically recognizes SL4 of the U2 snRNA^38^. SNRPB2 engages SL4 via its RRM, while SNRPA1 docks onto SNRPB2 to tune specificity and affinity (**Figure 5F**). Considering that SL4 was a major secondary structural element of the U1 snRNA removed by HL-1B-mediated stabilization of the TOE1–Sm complex, we hypothesized that SNRPB2 might prevent a similar processing of the U2 snRNA by physically hindering TOE1 interactions (**Figure 5F**). Supporting this idea, CRISPRi knockdown of SNRPB2 increased baseline and HL-1B-dependent over-trimming of the U2 snRNA (**Figures 5G** and **S11F**), resulting in the removal of both SL4 and SL3 and the halting of excess trimming ∼6 nucleotides downstream of the Sm complex binding site (**Figures 5G, 5H**). Other snRNAs were negligibly affected by knockdown of SNRPB2 (**Figures S11F, S11G**).

Taken together, our data identify opposing regulators of the phenotypic effects of molecular gluing of the TOE1–Sm complex, wherein high TOE1 and low SNRPB2 expression confer enhanced vulnerability to HL-1B through chemical-genetic synthetic lethality interactions.

### HL-1B modulates alternative splicing

Based on biochemical and structural studies of the spliceosome, removal of the U1 SL4 combined with stabilized binding of TOE1 to the Sm complex would be expected to disrupt the U1-SL4:SF3A1 bridge to the U2 snRNA^39–41^, thereby impeding the E→A transition and compromising 5′ splice site selection. The loss of SL4 might, however, be at least partly compensated for by the neighboring SL3, which remained intact in HL-1B-overtrimmed U1 snRNAs and has been shown to form independent interactions with SF3A1 and U2 through the bridging UAP56 and U2AF2 proteins^42^.

We first evaluated whether TOE1 was chemically glued onto spliceosomes that contain mRNA substrate. eCLIP-seq^43^ analysis revealed that HL-1B enhanced WT-TOE1 interactions with UTRs/introns (>5-fold), mRNA exons (>15-fold), and 3′/5′ splice sites (10-50-fold), while the number of enriched windows in the small RNA category (snRNAs/snoRNAs among others) changed minimally (**Figure S12A**). These HL-1B-induced shifts in WT-TOE1:RNA interactions were stereoselective and absent in C80S-TOE1-expressing cells (**Figures S12A, B**). When benchmarked against ENCODE eCLIP datasets^44^, HL-1B changed the primary RNA class of TOE1 interactors from snRNA/snoRNA toward splice-site-proximal mRNA regions (**Figure 6A**), consistent with the molecular gluing of TOE1 to the Sm complex placing this exonuclease in close proximity to pre-mRNAs bound to the spliceosome.

**Figure 6.**
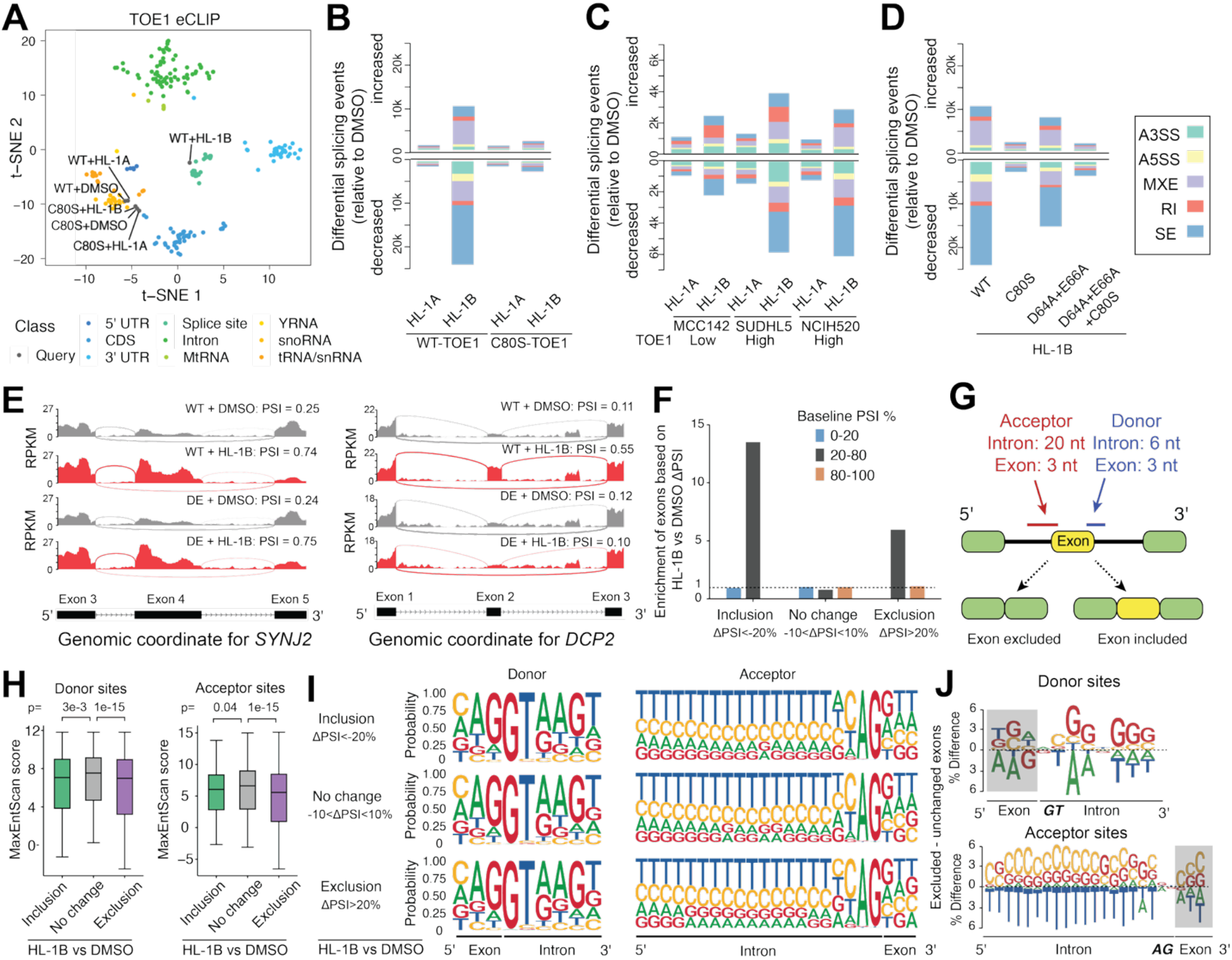
Alternative splicing effects of HL-1B and their modulation by TOE1 expression. (A) tSNE (t-distributed stochastic neighbor embedding) of TOE1 eCLIP data in the context of all available eCLIP datasets that were generated by the ENCODE consortium^44^. TOE1 eCLIP data were from WT-TOE1- and C80S-TOE1-expressing MCC142 cells treated with HL-1A or HL-1B (2 μM, 24 h) or DMSO. Data are average values from 2 independent experiments. (B) Summary of alternative splicing events induced by HL-1A or HL-1B (2 μM, 3 days) versus DMSO in WT-TOE1-and C80S-TOE1-expressing MCC142 cells, as quantified by RNAseq and rMATS analysis. Data for significantly increased or decreased splicing events are shown in upper and lower portion of graph, respectively. Significant events defined as having differential (DMSO vs probe) percent spliced in (ΔPSI) values of > 0.1 or < -0.1 and FDR < 0.05. Data are significant events from 3 independent experiments. A3SS, alternative 3′ splice site; A5SS, alternative 5′ splice site; MXE, mutually exclusive exons; RI, retained intron; SE, skipped exons. Inclusion forms are defined as: SE, target exon included; MXE, 5′ exon; A3SS/A5SS, longer exon; RI, intron retained. (C) Summary of alternative splicing events induced by HL-1A or HL-1B (2 μM, 24 h) versus DMSO in parental MCC142, SUDHL5, and NCIH520 cells that exhibit different expression levels of endogenous TOE1. (D) Summary of alternative splicing events induced by HL-1B (2 μM, 3 days) versus DMSO in MCC142 cells expressing the indicated TOE1 protein variants. (E) Sashimi plots of SYNJ2 and DCP2 splicing in WT-TOE1- and DE-TOE1-expressing MCC142 cells treated with HL-1B (2 μM, 3 days) or DMSO. (F) Barplot comparing the enrichment of different baseline exon usage groups for included (ΔPSI<-0.2), unchanged (-0.1<ΔPSI<0.1) or excluded exons (ΔPSI>0.2). Enrichment is defined as the proportion of exons in each baseline exon usage group within a given ΔPSI category divided by the proportion of that group seen across all exons. ΔPSI is calculated as DMSO - HL-1B. (G) Schematic showing the definition of flanking donor and acceptor sites for an exemplary exon going through alternative splicing. For donor sites, 3 nts of exon and 6 nts of downstream intron sequences were extracted. For acceptor sites, 3 nts of exon and 20 nts of upstream intron sequences were extracted. (H) Boxplots comparing the MaxEntScan scores of donor and acceptor sites associated with 3 classes of exons based on ΔPSI (DMSO - HL-1B). The box whiskers cover 95% of the data. P values were calculated using two-sided Student’s t test. (I) Nucleotide composition analysis for donor and acceptor sites of the 3 classes of exons based on ΔPSI (DMSO - HL-1B). (J) Nucleotide composition differences comparing the HL-1B-excluded versus unchanged exons for donor or acceptor site sequences.

We next evaluated the global splicing effects of HL-1B by RNA-seq in comparison to the general spliceosome inhibitor pladB in WT-TOE1-expressing MCC142 cells. We found that HL-1B induced far fewer alternative splicing changes than pladB (**Figure S12C**). Interestingly, the HL-1B-mediated splicing changes were mostly distinct from the effects of pladB (**Figure S12D**), pointing to a qualitatively distinct splicing program modulated by the TOE1–HL-1B–spliceosome complex.

The splicing effects induced by HL-1B in WT-TOE1-expressing MCC142 cells were both stereoselective and site-selective (**Figure 6B**). As one example of these changes, we show the HL-1B-dependent decrease in inclusion of exon 16 of the REXO5 gene (**Figure S12E**). We further found that cancer cell lines with higher basal TOE1 expression (SUDHL5, NCIH520^35^) displayed a greater number of HL-1B-induced splicing changes compared to a cell line with lower TOE1 expression (MCC142) (**Figure 6C**), and we confirmed these effects were attenuated by CRISPRi knockdown of TOE1 in NCIH520 cells (**Figure S12F**).

We next sought to determine the proportion of HL-1B-dependent splicing changes that were attributable to the physical gluing of the TOE1–Sm complex versus the resulting over-trimming of snRNAs. We surmised that these distinct biochemical effects of HL-1B could be parsed, at least in part, by comparing the splicing profiles of cells expressing WT-TOE1, the catalytically inactive D64A/E66A (DE) TOE1 mutant, and the respective C80S mutant forms of these proteins. We confirmed that the various WT and TOE1 mutant proteins were expressed at similar levels in MCC142 cells (**Figure S12G**) and found that the HL-1B effects on splicing and gene expression in WT-TOE1-expressing cells were partly attenuated in DE-TOE1-expressing cells, but not to the same extent as C80S-TOE1-expressing cells (**Figures 6D**, **S12H** and **Table S5**). Additionally, the residual splicing effects of HL-1B in the DE-TOE1-expressing cells were mostly ablated in DE/C80S-TOE1-expressing cells (**Figure 6D** and **Table S5**), suggesting that these changes were caused by the molecular gluing of the DE-TOE1–Sm complex and independent of TOE1 catalytic activity. From these results, we could partition HL-1B-sensitive splicing events into two classes (**Figure S12I**): (1) ∼30% that were shared by the WT-TOE1 and DE-TOE1-expressing cells and therefore interpreted to be caused by steric/architectural effects of TOE1 gluing (e.g., *SYNJ2*, where HL-1B switched the use of exon 5 to exon 4 in both cell models; **Figure 6E**); and (2) ∼70% that were specific to WT-TOE1-expressing cells and therefore interpreted to require exonuclease activity-dependent snRNA over-trimming (e.g., *DCP2*, where HL-1B increased exon 2 inclusion only in WT-TOE1-expressing cells; **Figure 6E**).

We finally found that exons with intermediate, rather than high or low baseline inclusion (as determined by percent spliced in (PSI) measurements), were more likely to be impacted by HL-1B treatment (**Figures 6F**, **S12J** and **Table S5**). To further characterize the HL-1B-regulated exons, we analyzed the splice-site strength of their flanking donor and acceptor sequences using maximum entropy modeling^45^ (**Figure 6G**). Compared to unchanged exons, exons showing altered inclusion or exclusion in HL-1B-treated cells displayed, on average, lower MaxEntScan scores, indicative of weaker splice sites (**Figure 6H**). The nucleotide composition at donor and acceptor sites associated with HL-1B-regulated exons revealed a modest enrichment of G/C nucleotides and a decrease in T (equivalent to U) nucleotides across acceptor sites in excluded exons (**Figures 6I, J**). These findings thus indicate that the HL-1B-mediated overtrimming of snRNAs modulates splicing in a unique manner that preferentially affects exons with partial rather than extreme percentages of inclusion or exclusion.

### HL-1B retains activity with hypomorphic disease variants of TOE1

Reduced TOE1 exonuclease activity leads to snRNAs that retain 3′ end extensions, and these immature species accumulate as stalled biogenesis intermediates that are subject to nuclear exosome–mediated decay rather than progressing to mature snRNPs for splicing^14,15^. Hypomorphic TOE1 mutations identified in patients with PCH7 exemplify this state and cause an accumulation of under-trimmed snRNA tails in cells that correlates with a mix of impaired TOE1 stability and partial loss of exonuclease activity *in vitro*^11,46^ (**Figure 7A**). Our findings have revealed that HL-1B exerts an opposing effect on snRNA processing: by gluing TOE1 onto snRNP complexes in a catalytically competent state, the compound prolongs the residence time of TOE1 with its snRNA substrates to promote their over-trimming (**Figure 7A**). We therefore wondered whether HL-1B might counteract the snRNA under-trimming phenotype observed with hypomorphic TOE1 variants, not by restoring their defective catalytic efficiency, but instead by extending their duration of substrate engagement in cells (**Figure 7A**).

**Figure 7.**
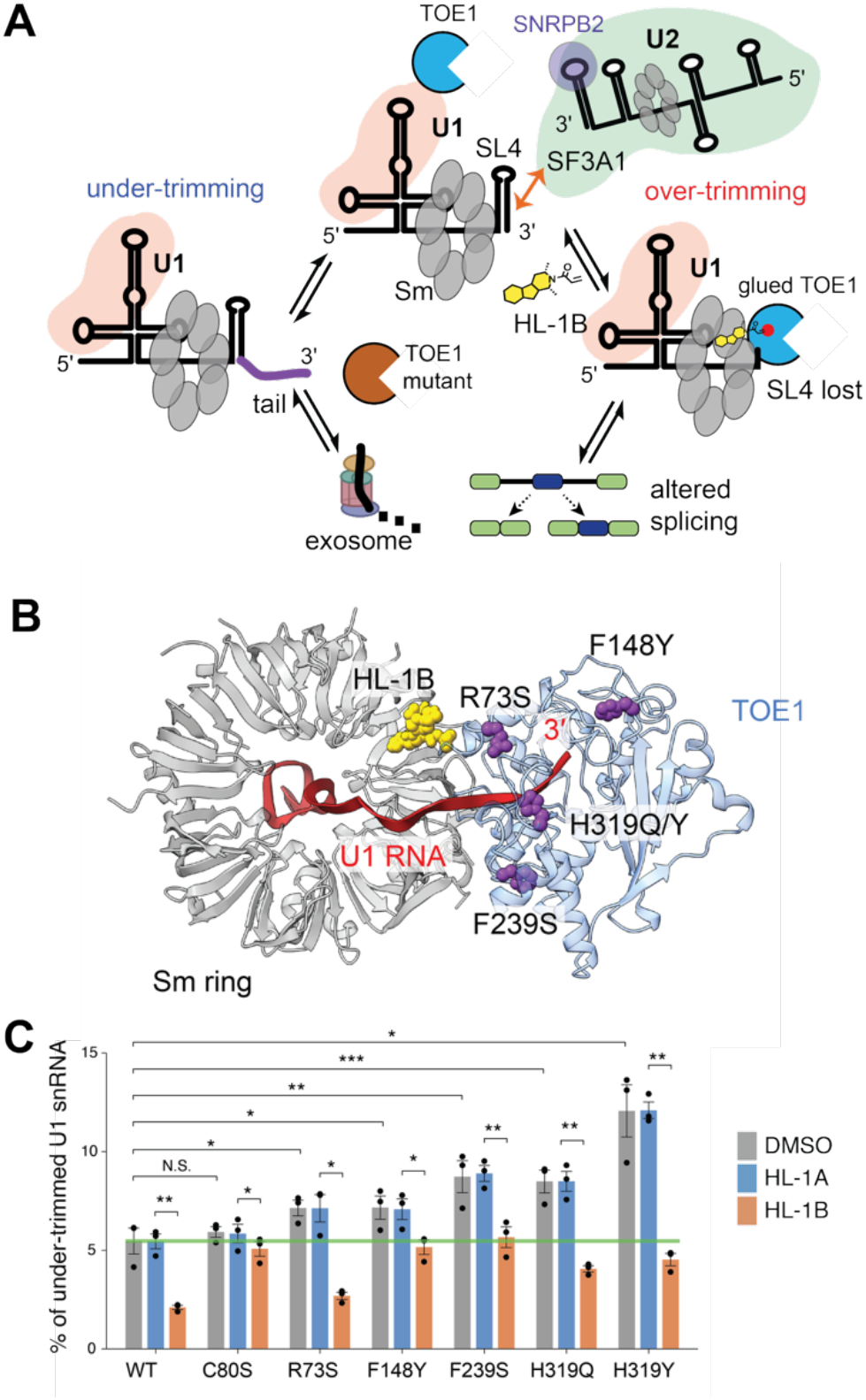
Counteracting effects of HL-1B and pathogenic TOE1 mutants on snRNA tail processing equilibrium. (A) A model for opposing effects of HL-1B and PCH7-associated mutants of TOE1 on the processing of snRNA 3′-tail lengths. (B) Location of PCH7-associated mutations in TOE1 in the TOE1–HL-1B-U1 snRNP structure. (C) Barplots comparing the 3′ tail lengths of the U1 snRNA in MCC142 cells expressing the indicated TOE1 variants and treated with HL-1A or HL-1B (2 μM, 3 days) or DMSO. Endogenous TOE1 levels were reduced in these cell models by using dCas9-KRAB paired with TOE1_sg1. The % under-trimming values were calculated as the fraction of U1 snRNA with 3′ end extended tail lengths ≥1 nucleotide from the canonical 3′ end position. Data are average values ± SEM from 3 independent experiments. All p values were calculated using two-sided paired Student’s t test. N.S., not significant, * p<0.05, **p<0.01, ***p<0.001.

We first localized PCH7-associated TOE1 mutations^11,46^ in the TOE1–HL-1B–U1 snRNP structure. Variants reported to cause partial loss of nuclease activity^11,46^ map to the DEDD nuclease domain (e.g., R73S, F148Y, F239S) and the C3H zinc finger (e.g., H319Q, H319Y) (**Figures S8A** and **S13A–D**). None of the mutated residues directly contact the U1 snRNA in the cryo-EM structure, nor do any lie within the composite HL-1B-binding pocket formed by TOE1_C80 and the SNRPE/SNRPF surface (**Figure 7B**). We interpreted these findings to indicate that the PCH7-associated mutants may remain capable of interacting with both the Sm ring and HL-1B. In support of this prediction, gel-ABPP experiments demonstrated that each of the aforementioned TOE1 mutants reacted in a stereoselective manner with HL-1B in cells (**Figure S13E**).

We next expressed each TOE1 variant in cells that were also knocked down for endogenous TOE1 by CRISPRi and measured U1 snRNA 3′ tail lengths by gene-specific 3′-end sequencing. At baseline, the mutants variably increased the fraction of under-trimmed U1 snRNA in comparison to control cells expressing WT-TOE1 or C80S-TOE1, with the H319Y mutant showing the largest magnitude effect (**Figure 7C**). Treatment with HL-1B significantly lowered the under-trimmed U1 snRNA fraction in cells expressing each hypomorphic mutant to control (or slightly below control) levels observed in DMSO-treated WT-TOE1-expressing cells (**Figure 7C**). The HL-1B-mediated amelioration of U1 snRNA under-trimming in TOE1 mutant-expressing cells was accompanied by increases in over-trimmed species that were mostly modest in magnitude with the exception of the R73S mutant, which showed a large increase in over-trimmed snRNA similar to WT-TOE1 (**Figure S13F**).

Our results thus indicate that HL-1B can compensate for the diminished catalytic activity of PCH7-associated TOE1 mutants by prolonging their interactions with snRNA substrates in the cell, supporting a model in which snRNA tail length is regulated by a kinetic balance between tail addition and exonuclease pathways, the latter of which can be augmented by chemically stabilizing TOE1-snRNP interactions (**Figure 7A**).

## Discussion

Among the diverse ways that small molecules regulate protein function in cells, the promotion or stabilization of protein-protein interactions (induced proximity) has emerged as a particularly attractive approach to expand the ligandability of the proteome^47,48^. Molecular glues, including the immunosuppressive natural products cyclosporin, FK506, and rapamycin, as well as the immunomodulatory drug (IMiD) class of protein degraders, often bind independently to a presenter protein to create a composite surface that then recruits and perturbs the function of a previously undruggable neo-partner protein^48–55^. HL-1B, on the other hand, stabilizes a dynamic endogenous protein-protein interaction and may therefore alternatively be described as a molecular clamp, an emerging category of compounds that can act through diverse mechanisms involving the binding to allosteric sites on a single protein or to composite sites shared by both partners^48,49,56–60^. Our work shows that this form of induced proximity can provide a means to chemically regulate the structure and function of a specific class of RNA biomolecules (snRNAs) through strengthening a catalytically competent complex between an exonuclease (TOE1) and its ribonucleoprotein (snRNP) substrates. This feat was accomplished with a monofunctional covalent small molecule (HL-1B), thus offering a compelling and complementary approach to emerging heterobifunctional compounds (RIBOTACs) that promote the nuclease-mediated degradation of RNAs in cells^61,62^.

TOE1 was originally found to bind components of the spliceosome by IP-MS^11,26^ and has more recently been shown to mainly associate with preassembled Sm cores as a key determinant of snRNA 3′-end maturation and for discrimination of functional versus defective snRNAs^11,14,15,26^. The structural basis for these TOE1-spliceosome interactions, however, has remained unclear. Our cryo-EM structure provides a compelling model for how HL-1B stabilizes the transient and catalytically competent interactions between TOE1 and the Sm complex within the U1 snRNP. By binding to a non-orthosteric composite pocket distal from the TOE1 active site, HL-1B substantially increases the surface area of interaction between TOE1 and the Sm complex while leaving the active site of TOE1 unperturbed. As a consequence, HL-1B is inferred to increase the residence time of TOE1 in proximity to its snRNA substrates in the cell, ultimately leading to excessive processing (over-trimming) of their 3′ ends, which in the case of the U1 snRNA removes the SL4 stem loop. We speculate that the base pairing and stacking energy of SL4 and other U snRNA 3′ stem loops impose a thermodynamic barrier to continued 3′→5′ processing by TOE1 under basal conditions where the TOE1-Sm ring interactions are transient. By gluing TOE1 onto snRNP complexes, HL-1B overrides this barrier to produce a striking and, to our knowledge, unprecedented pharmacological outcome in cells: the discrete *over-trimming* of the 3′ stem loops of snRNAs and formation of new termination sites that are set by the location of the Sm complex on these snRNAs.

Previous studies have established that SL4 of the U1 snRNA is important for splicing; the interaction between SL4 in the U1 snRNA and the UBL domain of SF3A1 in the U2 snRNP is one of the molecular bridges that help to pair the 5′ and 3′ splice sites during early spliceosome assembly^39–41^. However, impairments in SL4-SF3A1 interactions can, in part, be compensated for by additional U1-U2 contacts mediated by SL3 of U1 and its bridging protein partner UAP56^42^. We found that, in comparison to direct spliceosome inhibitors, such as the natural product pladB, that induce widespread splicing collapse, impairments in nascent transcription, and general cytotoxicity^19,63^, the splicing alterations induced by HL-1B were preferentially enriched among exons exhibiting intermediate baseline inclusion levels, while largely sparing exons with constitutively high or low inclusion. We postulate that splice sites with strong signals remain saturated under these conditions, whereas sites that depend on efficient U1–U2 communication for commitment are disproportionately affected, consistent with the selective impact of HL-1B on exons with intermediate inclusion. In this model, loss of SL4 converts spliceosomes into a new state with partly corrupted U1-U2 crosstalk, leading to pharmacological effects of HL-1B that are restricted to “regulatable” splice sites intrinsically poised for dynamic control. Considering further the recent discovery of additional proteins that interact with SF3A1 in a stress-responsive manner^64^, it appears that the SL4-SF3A1 subcomplex may represent a specialized unit for dynamic regulation of spliceosome function.

Our large-scale CRISPRi screen identified additional factors that can tune the snRNA trimming profile of cells exposed to HL-1B. These include: i) the expression level of TOE1 itself, which influences the fraction of each snRNA that undergoes over-trimming; and ii) SNRPB2, which binds to SL4 of the U2 snRNA and shields this stem loop from TOE1 processing. Notably, these positive (TOE1) and negative (SNRPB2) regulators also impacted the antiproliferative effects of HL-1B, illuminating synthetic lethality interactions that may serve as future biomarkers for predicting cancer types that are sensitive to the molecular gluing of TOE1–Sm complexes. Similar opportunities for synthetic lethality interactions have been noted with other small-molecule modulators of the spliceosome, such as the RBM39 degrader indisulam^22–24^, where the expression level of the E3 ligase DCAF15 correlates with cancer cell growth inhibition^65^.

From a broader translational perspective, our work provides new insights into how PCH7-associated TOE1 mutations impair snRNA processing, as well as how HL-1B may counteract these effects. H319 coordinates a zinc ion within a zinc finger that also contains W308, which stacks against the terminal nucleotides of U1 snRNA and likely helps position the substrate for hydrolysis. Mutations at H319 may therefore perturb substrate positioning by altering the zinc-finger structure. F239S may impair TOE1 association with the Sm ring, as F239 stabilizes a loop that directly contacts SNRPD2. Interestingly, HL-1B reduces U1 under-trimming caused by these mutations without also promoting substantial over-trimming. This restorative profile may indicate that the HL-1B-induced stabilization of TOE1–Sm interactions is sufficient to partly compensate for mutation-related defects in substrate positioning or binding to the U1 snRNP such that proper U1 tail processing is restored, but excessive trimming is minimized.

Although our preliminary results suggest that HL-1B can alleviate U1 snRNA under-trimming caused by hypomorphic TOE1 variants, future studies are needed to determine the extent to which TOE1-dependent snRNA processing defects contribute directly to PCH7 pathogenesis and whether HL-1B can rescue the disease phenotypes associated with this neurological disorder. We also note that some of the PCH7 mutants in our panel, such as R73S, did not cause major under-trimming of the U1 snRNA despite R73S being positioned to interact with 3′-terminal nucleotides in the active site. The R73S-TOE1 mutant, however, also showed the highest expression in our cell models, which may have led to an underestimate of its tail processing effects. It will additionally be important to understand the functional consequences of HL-1B-dependent under-trimming of other ncRNAs (snoRNA, scaRNA, TERC), which we hypothesize is the result of ligand-induced sequestration of TOE1 in spliceosome complexes. Indeed, a similar mechanism may explain why covalently liganded TOE1 isolated from cells shows impaired activity with exogenous RNA substrates^26^. Finally, we call attention to recurrent hotspot cancer mutations in the spliceosome and, more specifically, at the 5′ end of the U1 snRNA^66,67^. It would be interesting to investigate how molecular gluing of the TOE1-Sm complex impacts splicing in the context of such cancer-relevant spliceosome mutations.

In summary, the pursuit of small molecules that can regulate RNA structure and function in cells remains a major goal of chemical biology and drug discovery. Identifying compounds that directly bind RNA represent one exciting avenue of research^68–70^, but is challenged by the somewhat uniform physicochemical properties and limited structural diversity of RNAs^71^. Our findings provide a compelling conceptual illustration for how small molecules can control ncRNA structure and function by stabilizing protein-protein interfaces that physically redistribute an ncRNA processing enzyme in the cell. We believe this form of induced proximity should be applicable to additional ncRNA-protein complexes to provide chemical probes that complement more traditional genetic loss-of-function approaches, propelling both fundamental and translational research.

## Supporting information

Synthetic Chemistry Information

## Materials availability

All unique reagents generated in this study will be made available upon reasonable request without restrictions.

## Data availability

Raw sequencing datasets have been deposited in Sequence Read Archive (SRA) database under BioProject PRJNA1358761. The proteomics data have been deposited in the ProteomeXchange Consortium via the PRIDE repository with the dataset identifiers PXD070432, PXD070435, and PXD070450. The cryo-EM density maps generated in this study have been deposited in the Electron Microscopy Data Bank (EMDB) under accession codes EMD-79946 (consensus reconstruction), EMD-79947 (TOE1–TOE1 focused reconstruction), EMD-79948 (U1–TOE1 focused reconstruction), EMD-79949 (Sm-ring focused reconstruction), and EMD-79950 (composite reconstruction). The corresponding atomic coordinates have been deposited in the Protein Data Bank (PDB) under accession code 40BP. Additional data supporting the findings of this study are available from the corresponding authors upon reasonable request.

## Code availability

The custom code used in this paper is deposited in GitHub (https://github.com/jasonli1314/TOE1_clamp).

## Acknowledgments

We thank Nino Makasarashvili, Rees Garmann, Colin Fowler, and Karen Adelman for their kind feedback. We thank John Shimashita, Steven Head, and Sean Wang at the Scripps Sequencing Core facility for assistance with sequencing. We thank Kathryn Spencer and Scott Henderson at the Scripps Microscopy core for assistance with confocal imaging. We thank WuXi AppTec for the synthesis of chemical probes used in this study and the Scripps Automated Synthesis Facility for HRMS measurements. We thank Steven Blue and Assael Madrigal in the Yeo lab for guidance in performing eCLIP. We acknowledge this work was supported by the Damon Runyon Cancer Research Foundation (DRG: 2406-20 awarded to H.L.), NCI (R35 CA231991 awarded to B.F.C. and K99/R00 CA290143 awarded to H.L.), NIGMS (R35 GM127090 awarded to I. J. M. and R35 GM118069 awarded to J.L.-A.), NHGRI (R01 HG004659 awarded to G.W.Y).

## Author contributions

Conceptualization: H.L., B.F.C.; Data curation: H.L., L.F.R.G., T.M., J.T., B.M.; Formal analysis: H.L., L.F.R.G., B.M., T.M.; Investigation: H.L., L.F.R.G., T.M., K.F., J.T.; Software: H.L., L.F.R.G., T.M., B.M.; Supervision: G.W.Y., B.M., J.L.-A., I.J.M., B.F.C.; Visualization: H.L., L.F.R.G., K.F., B.M., I.J.M.; Writing – original draft: H.L., L.F.R.G., B.M., I.J.M., B.F.C.; Writing – review & editing: all authors.

## Competing interests

B.F.C. is a founder and scientific advisor to Vividion Therapeutics. G.W.Y. is an SAB member of Jumpcode Genomics and a co-founder, member of the Board of Directors, on the SAB, equity holder, and paid consultant for Eclipse BioInnovations. G.W.Y. is a visiting professor at the National University of Singapore. G.W.Y.’s interests have been reviewed and approved by the University of California, San Diego in accordance with its conflict-of-interest policies. B.F.C. and B.M., and H.L. are inventors on a provisional patent application filed by The Scripps Research Institute covering covalent ligands targeting cysteine 80 of TOE1.The other authors declare no competing interests.

## Methods

### Cell lines

Cell lines used in this study were MCC142 (ECACC), TC71 (Texas Tech University), HeLa S3 (ATCC), PANC1005 (ATCC), FaDu (ATCC), MERO-14 (Sigma), JEG-3 (ATCC), ES-2 (ATCC), RKN (JCRB), CALU-6 (ATCC), T98G (ATCC), HT-1080 (ATCC), A375 (ATCC), and NCIH520 (ATCC), SUDHL5 (ATCC). All lines were authenticated by the providers using short tandem repeat (STR) profiling and were maintained under standard culture conditions (37 °C, 5% CO_2_) in RPMI-1640 medium (Gibco) supplemented with 10% FBS (Omega Scientific), 2 mM GlutaMAX (Gibco), 100 U/mL penicillin, and 100 μg/ml streptomycin (Gibco). Cultures were routinely tested and confirmed negative for microbial contamination, including mycoplasma.

### Creation of stable cell lines

For lentivirus production, plasmids comprising the gene-of-interest construct, packaging vector (pCMV-dR8.91), and envelope vector (VSV-G) were co-transfected into 293T cells at a 9:6:1 ratio in OPTI-MEM. Polyethylenimine (PEI; Polysciences) was added at 3 μl of 1 mg/ml solution per μg total DNA, and DNA–PEI complexes were incubated 20 min at room temperature before being added dropwise to cells at ∼50% confluence. After 8 h, the medium was replaced with fresh DMEM (Corning) supplemented with 30% FBS, penicillin–streptomycin, and 2 mM GlutaMAX. Viral supernatants were collected at 48 h post-transfection and clarified by filtration through 0.45 μm syringe filters (Millipore). For stable line generation, target cells were seeded in 12-well plates and infected with viral supernatant containing 8 μg/ml polybrene (Millipore). Spin-infection was performed at 900 × g for 1 h at 30 °C. The following day, cells were split into 10 cm plates. After another day, antibiotic selection appropriate to the vector was initiated to enrich for successfully transduced cells.

### Synthetic chemistry

See Supplementary Synthetic Chemistry file for information on synthetic chemistry procedures.

### Gel ABPP

Approximately 2 × 10^6 MCC142 cells stably expressing FLAG-tagged TOE1 were treated with competitor probes followed by alkyne probes, or with alkyne probes alone, in RPMI media. After treatment, cells were collected, washed with ice-cold PBS, and lysed in 300 μl of buffer (25 mM Tris-HCl, pH 7.5, 150 mM NaCl, 1% Triton X-100) followed by sonication (20 pulses, 10% power output). Lysates were clarified by centrifugation at 16,000 × g for 5 min, and soluble protein fractions were normalized. Ten percent of the normalized lysate was reserved as input, and the remainder was incubated with anti-FLAG M2 magnetic beads at 4 °C for 2 h. Beads were washed twice with wash buffer (25 mM Tris-HCl, pH 7.5, 150 mM NaCl, 0.1% Triton X-100). For click chemistry labeling, a reaction mix containing 1 μl CuSO_4_ (50 mM), 3 μl TBTA ligand (1.7 mM in DMSO:tBuOH, 1:4 v/v), 1 μl freshly prepared TCEP (50 mM), and 0.5 μl rhodamine-azide (1.25 mM) was mixed separately in order and then added to each sample in 50 μl PBS. After 1 h, samples were mixed with loading dye, denatured at 90 °C for 5 min, and analyzed by SDS–PAGE. Samples were visualized using the ChemiDoc MP system.

### MS ABPP

Approximately 1 × 10^7 parental MCC142 cells were treated with the indicated compounds for the indicated time in 15 cm plates. Cells were scraped, washed with ice-cold PBS, frozen at -80°C before further processing. The step-by-step protein-directed ABPP, cysteine-directed ABPP protocols and MS instrumentation parameters have been described in detail^21,29,30^.

### Immunoprecipitation mass spectrometry (IP-MS)

Cells stably expressing N-FLAG–tagged WT TOE1, the C80S mutant, or parental control were treated in situ with the indicated compounds for the indicated time. Cells were harvested, washed with ice-cold PBS, and frozen. Pellets were lysed in lysis buffer (50 mM EPPS, pH 7.5, 150 mM NaCl, 1% Triton X-100, 10% glycerol) supplemented with cOmplete Protease Inhibitor Cocktail (Roche). Lysates were sonicated with a probe sonicator (32 pulses, 10% energy output), incubated on ice for 10 min, and clarified by centrifugation at 16,000 × g for 5 min. Protein concentrations were determined using a BCA assay (Pierce), and 2 mg of protein from each sample was incubated with 40 μl of pre-washed anti-FLAG M2 agarose slurry (Sigma) at 4 °C for 3 h with rotation. Beads were collected by centrifugation and washed three times with IP wash buffer (50 mM EPPS, pH 7.5, 150 mM NaCl, 0.1% Triton X-100) and twice with 50 mM EPPS, pH 7.5 (2,000 × g, 1 min each). Bound proteins were eluted in 50 μl of 8 M urea in EPPS at 65 °C for 10 min and then filtered by spin-columns to remove agarose beads. The proteins were reduced with 200 mM DTT at 65 °C for 15 min, alkylated with 400 mM iodoacetamide at 37 °C for 30 min, and diluted to 2 M urea with 50 mM EPPS, pH 7.5. Proteins were digested overnight at 37 °C with MS-grade trypsin (Promega), TMT-labeled (Thermo Scientific), desalted on Sep-Pak C18 cartridges (Waters), and analyzed using an Orbitrap Fusion™ mass spectrometer (Thermo Fisher).

### Size exclusion chromatography-mass spectrometry (SEC–MS)

The SEC–MS platform was performed as previously described with minor modifications^21^. MCC142 cells were treated *in situ* with the indicated compounds, washed with ice-cold PBS, and frozen at −80 °C prior to processing. Frozen cell pellets were lysed by probe sonication (32 pulses, 10% power) in 650 μl ice-cold PBS supplemented with cOmplete protease inhibitors (Roche), clarified by centrifugation at 16,000 × g for 5 min, and further ultracentrifuged at 100,000 × g for 30 min. A total of 500 μl clarified lysate was loaded onto a Superdex 200 Increase 10/300 GL column connected to an ÄKTA Pure FPLC system (Cytiva) and fractionated in PBS at 0.5 ml/min into five 2 ml fractions. Eluates were collected into 15 ml tubes containing 12 ml of cold acetone and incubated overnight at −20 °C to precipitate proteins, which were pelleted by centrifugation at 4,500 × g for 20 min. Pellets were air-dried and resuspended in 125 μl 8 M urea in 100 mM EPPS (pH 7.5) supplemented with 10 mM DTT, at 65 °C for 15 min, followed by probe sonication. Samples were alkylated with 6.25 μl of 500 mM iodoacetamide at 37 °C for 30 min, diluted with 370 μl of 100 mM EPPS, and digested overnight at 37 °C with 2 μg MS-grade trypsin per fraction. For TMT labeling, 70 μl of digest was mixed with 30 μl acetonitrile and 6 μl TMT reagent (20 μg/μl in acetonitrile), incubated at room temperature for 1 h, quenched with 6 μl of 5% hydroxylamine for 15 min, and acidified with 6 μl formic acid. Labeled fractions were pooled, dried by speed vacuum, desalted on Sep-Pak cartridges (Waters), and fractionated at high pH as previously described before MS analysis^21^.

### Purification of human U1 snRNP

HeLa S3 cells (4 × 10^9) were harvested after PBS washes and base buffer (25 mM HEPES, pH 7.5, 150 mM NaCl, 0.1 mM DTT) was maintained on ice. Cell pellets were resuspended and lysed by probe sonication in base buffer supplemented with 1% Triton X-100 and cOmplete Protease Inhibitor Cocktail (Roche). Lysates were clarified by centrifugation at 3,000 × g for 5 min followed by 30,000 × g for 30 min at 4 °C. Clarified lysates were mixed with 60 μl of 100 μM desthiobiotin-labeled U1 oligonucleotides (mAmGmGmUmAmAmGmUmAmUmAmAmGmGmUmAmAmGmU-mAmUmAmAmGmGmUmAmAmGmUmAmUmA/3deSBioTEG/, 2’ O-methyl bases with HPLC Purification) for 1 h at 4 °C, then mixed with high-capacity streptavidin beads (30 ml slurry) for 3 h at 4 °C with gentle stirring. Beads were captured on a gravity column and washed extensively with base buffer. Bound U1 snRNP was eluted by overnight incubation in base buffer containing 10 mM biotin (pH 7.5) with gentle stirring at 4 °C. The next day, beads were removed by gravity flow and the eluate was concentrated using protein concentrators (30 kDa MWCO; Pierce) and was aliquoted and stored at −80 °C until use.

### Purification of recombinant His-TOE1

The coding sequence for N-His-tagged TOE1 was cloned into pET24 (Twist) between EcoRI and NotI restriction sites. Recombinant protein was expressed in E. coli BL21 Star™ (DE3) by induction with 0.5 mM IPTG for 18 h at 16 °C after cultures reached OD_600_ about 0.6. Bacterial cells were pelleted by centrifugation and frozen at −80 °C prior to purification. Frozen pellets were resuspended in base buffer (50 mM HEPES, pH 7.5, 150 mM NaCl, 0.1 mM DTT, 0.1% Triton X-100) and lysed by probe sonication (Emerson Branson). Lysates were clarified by centrifugation (30,000 × g, 30 min), and His-TOE1 was captured on TALON® metal affinity resin (Takara Bio) by gravity flow. The resin was washed extensively with base buffer and eluted with base buffer containing 150 mM imidazole. Eluates were dialyzed (SnakeSkin, Thermo Scientific) against base buffer, concentrated (Pierce concentrators, 10 kDa MWCO), flash-frozen in liquid nitrogen, and stored at −80 °C until use.

### AlphaFold structure prediction and confidence assessment

The structure of human TOE1 was predicted using AlphaFold 3^28^ (AF3) through the AlphaFold Server, using the full-length amino acid sequence of human TOE1 (UniProt: Q96GM8) as input. Five structural models were generated and visualized in UCSF ChimeraX. To assess model confidence and structural consistency, residues with predicted local distance difference test (pLDDT) scores below 50 were removed, followed by the removal of remaining contiguous peptide fragments shorter than eight residues. Per-residue pLDDT scores were extracted from all five predictions and plotted individually and as their mean using R. The curated models showed high structural similarity, and model 1 was selected as the starting model for subsequent cryo-EM model building. The predicted aligned error (PAE) matrix of model 1 was visualized as a heatmap.

### Cryo-EM sample preparation

A commercial preparation of purified bovine snRNPs (Surmodics IVD) was validated by proteomic analysis and shown to consist predominantly of U1 snRNP proteins, with minor fractions of other U snRNP proteins (**Figure S4A**). Purified recombinant TOE1 (50 μg) was mixed with purified bovine U1 snRNP (50 μg) in a final volume of 480 μl PBS containing 2 mM MgCl_2_ and 0.1 mM DTT and incubated for 5 min at room temperature. The TOE1–U1 mixture was then supplemented with 20 μl HL-1B to a final concentration of 25 μM and incubated for 1 h at room temperature. Benzonase (20 units in 1 μl) was added, and the reaction was incubated for 10 min at 37 °C before being quenched with 5 mM EDTA (final) and cooled on ice. The mixture was buffer-exchanged into 25 mM HEPES (pH 7.5), 150 mM NaCl, 0.1 mM DTT and then concentrated to 20 μl using a Vivaspin® 500 centrifugal concentrator (Sartorius, MWCO 100 kDa) prior to freezing. The *in vitro*-assembled sample was diluted to a nominal U1 snRNP concentration of 1.2 mg/mL, calculated from the amount of U1 snRNP used in the assembly reaction, in 25 mM HEPES (pH 7.5), 150 mM NaCl, and 0.1 mM DTT. UltrAuFoil R1.2/1.3 300-mesh gold grids (Quantifoil) were glow-discharged for 30 s at 15 mA using a PELCO easiGlow system. A 3 µL aliquot of sample was applied to each grid and blotted for 4 s at a blot force setting of 6 using Whatman No. 1 filter paper in a Vitrobot Mark IV (Thermo Fisher Scientific), maintained at 4 °C and >95% relative humidity. Grids were subsequently plunge-frozen in liquid ethane.

### Cryo-EM data collection

Cryo-EM data were collected using a Talos Arctica transmission electron microscope (Thermo Fisher Scientific) operated at an accelerating voltage of 200 kV and equipped with a Falcon 4i direct electron detector. Data acquisition was performed using EPU software (version 3.9.1.8206), with nominal defocus settings of −1.5 and −2.5 µm. Movies were acquired over 3.37 s with a total electron exposure of 40 e⁻/Å², using a 50 µm C2 aperture and a nominal magnification of 190,000×, corresponding to a calibrated pixel size of 0.74 Å. Initial data collection revealed pronounced preferred particle orientation, which persisted despite testing alternative sample preparation conditions, including the addition of detergents and the use of graphene oxide and continuous carbon supports. To improve angular sampling, additional datasets were collected at nominal specimen tilt angles of 30° and 40°.

### Cryo-EM data processing and model building

The cryo-EM data processing workflow is summarized in **Figures S5** and **S6**. A total of 21,248 movies were collected across multiple acquisition sessions and processed using cryoSPARC Live (Structura Biotechnology). Following motion correction and contrast transfer function (CTF) estimation, exposures with estimated tilt angles greater than 20° and CTF fit resolutions between 2 and 8 Å were selected for downstream processing, yielding 7,832 exposures. Particles were identified through iterative rounds of blob picking, template-based picking, and Topaz picking, initially using subsets of exposures. Particles were extracted using a box size of 360 pixels and Fourier-cropped to 180 pixels, corresponding to a pixel size of 1.48 Å. The final particle selection comprised 121,195 particles from four acquisition sessions.

A subset of 40,000 particles was used to generate four *ab initio* reconstructions, yielding a pseudo-tetrameric complex comprising two TOE1 proteins and two U1 Sm-ring assemblies, a pseudo-dimeric complex, and two poorly resolved classes. These four reconstructions were subsequently used as references for heterogeneous refinement to separate the particle populations. The resulting pseudo-tetrameric class contained 53,467 particles and exhibited well-defined structural features, including density in the putative ligand-binding region. Homogeneous refinement without imposed symmetry (C1) yielded a reconstruction at 3.8 Å resolution, as determined by the gold-standard Fourier shell correlation (FSC) criterion of 0.143. Particles were subsequently recentered and re-extracted using an unbinned box size of 512 pixels at 0.74 Å/pixel, followed by a final C1 homogeneous refinement to generate the consensus reconstruction (EMD-79946). The consensus reconstruction was used for initial rigid-body fitting of the U1 snRNP component from PDB 7B0Y and the AF3-predicted TOE1 structure.

Focused refinements were subsequently performed to improve the density of individual regions of the complex. For the TOE1–TOE1 interface, local refinement was performed without imposed symmetry (C1), using a mask encompassing both TOE1 proteins. Density outside the region of interest was removed by particle subtraction, followed by a final C1 local refinement, yielding a reconstruction at 3.86 Å resolution (FSC = 0.143; EMD-79947). For the U1–TOE1 subcomplex, particles were symmetry-expanded using C2 symmetry to account for the approximately twofold-related subcomplexes within the tetramer. Following local refinement, density outside the region of interest was removed by particle subtraction, and a final C1 local refinement yielded a reconstruction at 3.61 Å resolution (FSC = 0.143; EMD-79948). The Sm-ring region was processed similarly, using a more restricted focus mask, yielding a reconstruction at 3.77 Å resolution (FSC = 0.143; EMD-79949).

The focused reconstructions were aligned to the C1 TOE1–TOE1 reconstruction and combined in UCSF ChimeraX using a voxel-wise maximum operation to generate a composite map (EMD-79950). As the composite map was generated computationally from multiple focused reconstructions, an independent gold-standard FSC resolution was not determined. The constituent focused reconstructions had global resolutions ranging from 3.61 to 3.86 Å, corresponding to a mean resolution of approximately 3.75 Å. The focused reconstructions were aligned and combined in UCSF ChimeraX (version 1.12) using a voxel-wise maximum operation and resampled onto the consensus grid to generate a composite map (EMD-79950).

The U1 snRNP structure from PDB 7B0Y and the AF3-predicted TOE1 model were rigid-body fitted into the composite density and subjected to real-space refinement using Phenix (version 1.21.2-5419). Additional density at TOE1 residue C80 was consistent with covalent modification by HL-1B, allowing the modified residue to be modeled. The chemical structure of the modified residue was generated in ChemDraw and exported as a SMILES representation. Geometry restraints were generated using Phenix eLBOW and incorporated into subsequent real-space refinement. The atomic model was iteratively improved through manual model building in Coot (version 0.9.8.96) and ISOLDE, interspersed with real-space refinement in Phenix. Model geometry and stereochemical quality were assessed using MolProbity. The final atomic coordinates were deposited in the Protein Data Bank (PDB ID: 40BP), and the corresponding cryo-EM reconstructions were deposited in the Electron Microscopy Data Bank under accession codes EMD-79946–EMD-79950.

### *In vitro* reconstitution of TOE1-U1 snRNP complex for gel ABPP

Purified recombinant TOE1 (5 μg) was mixed with purified human or bovine U1 snRNP (10 μg) in a final volume of 48 μl PBS and incubated at room temperature for 5 min. The TOE1–U1 mixture was then supplemented with alkyne probes (2 μM, 1 h) only or competitor probes (1 h) followed by alkyne probes (2 μM, 1 h) and incubated at room temperature. For CuAAC reaction, a click mix comprising 1 μl CuSO_4_ (50 mM), 3 μl TBTA ligand (1.7 mM in DMSO:tBuOH, 1:4 v/v), 1 μl fresh TCEP (50 mM), and 0.5 μl rhodamine–azide (1.25 mM) was prepared in this order, then added to each sample. After 1 h, samples were mixed with loading dye, denatured at 90 °C for 5 min, resolved by SDS–PAGE. Rhodamine fluorescence was imaged first, after which the gel was stained with InstantBlue Coomassie (Abcam) to visualize total protein using a ChemiDoc MP system.

### *In vitro* reconstitution of TOE1-U1 snRNP complex for MS analysis

Purified recombinant TOE1 (2 μg) was mixed with purified bovine U1 snRNP (4 μg) in a final volume of 18 μl PBS and incubated at room temperature for 5 min. The TOE1–U1 mixture was then supplemented with HL-1B at the indicated doses and incubated at room temperature for 1 h. Proteins were mixed with 40 μl of 8 M urea in EPPS at 65 °C for 10 min, reduced with 200 mM DTT at 65 °C for 15 min, alkylated with 400 mM iodoacetamide at 37 °C for 30 min, and diluted to 2 M urea with 50 mM EPPS, pH 7.5. Proteins were digested overnight at 37 °C with MS-grade trypsin (Promega), TMT-labeled (Thermo Scientific), desalted on Sep-Pak C18 cartridges (Waters), and analyzed using an Orbitrap Fusion™ mass spectrometer (Thermo Fisher).

### RNA 3′ end sequencing

Total RNA was extracted from cells using TRIzol, followed by column-based purification. 2 μg total RNA was treated with FastAP (1 μl), T4 polynucleotide kinase (1 μl), Turbo DNase (1 μl), and RNase inhibitor (1 μl) in a total of 40 μl reaction buffer at 37 °C for 30 min followed by RNA clean-up (Zymo). Cleaned RNA was mixed with 2 μl of 40 μM AG10 adapter and heated at 65 °C for 2 min before adding T4 RNA ligase (1 μl), 10 mM ATP (2 μl), RNase inhibitor (1 μl) in a total of 20 μl reaction buffer at 16 °C overnight. Ligated RNA was column-purified, mixed with 10 mM dNTP (1 μl) and 20 μM AR17 (1 μl), heated at 65 °C for 2 min before adding SuperScript III reverse transcriptase (1 μl) and RNase inhibitor (1 μl) in a total of 20 μl reaction buffer. After incubating at 55°C for 1 h and 70°C for 15 min, the resulting cDNA was diluted 6-fold and then amplified in a two-step PCR strategy using the Phusion High-Fidelity PCR Kit. The first PCR used AR17 (1 μl of 10 μM) and gene-specific primers with inline barcodes (1 μl of 10 μM) to amplify 5 μl cDNA in a total of 20 μl. The PCR1 reactions were carried out as follows: 98°C for 2 min, 6 cycles of (20 s at 98 °C, 25 s at 50 °C, 25 s at 72 °C), 12 cycles of (20 s at 98 °C, 25 s at 55 °C, 25 s at 72 °C), followed by final extension at 72 °C for 2 min. The PCR1 products were pooled based on the inline barcodes and purified using Mag-Bind® TotalPure NGS beads following manufacturer’s protocol. PCR2 uses Illumina indexing primers (1 μl of 20 μM per 20 μl reaction). The PCR2 reactions were carried out as follows: 98°C for 2 min, 6 cycles of (20 s at 98°C, 25 s at 56 °C, 25 s at 72 °C), 2 cycles of (20 s at 98 °C, 25 s at 68 °C, 25 s at 72 °C), followed by final extension at 72 °C for 2 min. Final PCR products were pooled and purified using Mag-Bind® and sequenced on an Illumina platform. For global 3′ end sequencing, the protocol was based on the previous study without change^14^. The custom pipeline used in the analysis is modified from the Tailer package^76^ and is available in Github (https://github.com/jasonli1314/TOE1_clamp).

### Immunofluorescence microscopy and quantification

MCC142 cells were grown on poly-D-lysine-coated coverslips in 12-well plates and were treated with the indicated compound for the indicated time. Cells were washed with PBS and fixed in freshly prepared 4% paraformaldehyde diluted in PBS for 15 min at room temperature, followed by quenching with 0.3 M glycine and additional PBS washes. Samples were permeabilized with 0.1% Triton X-100 diluted in PBS for 30 min and blocked in 5% normal goat serum with 0.1% Triton X-100 for 1 h at room temperature. Primary antibodies were diluted in PBS containing 1% goat serum and were incubated overnight at 4 °C. The primary antibody dilutions include FLAG, 1:100; coilin, 1:800; SRSF2, 1:200; SNRNP70, 1:500 (vol/vol). After PBS washes, samples were incubated for 1 h with Alexa Fluor 488-/568-/647-conjugated secondary antibodies diluted 1:500 (vol/vol) with 1% goat serum in PBS, followed by additional washes. Nuclei were stained with Hoechst (0.1 μg/ml), and samples were washed, covered in mounting media, and cured in the dark for 2 days before imaging. Images were collected sequentially on a Zeiss LSM 780 confocal microscope with a Plan-Apochromat 63x NA 1.4 objective. Images were then processed with Cell Profiler 4.2.8 using the IdentifyPrimaryObjects function followed by the MeasureColocalization function for each cell detected in the images. The resulting correlations were exported and plotted in R (v4.5.1).

### Cell growth assay

Cells were seeded at 4,000 cells per well in 96-well, flat-bottom, white-wall plates (Greiner Bio-One) in 190 μl medium. 10 μl medium containing DMSO or compound dilutions (20× stocks to achieve the desired final concentrations) was added to each well. After 3 days, the medium containing probes was replaced with fresh medium and the same treatments, and incubation continued for an additional 3 days. Cell viability was measured using CellTiter-Glo (Promega) on an EnVision multimode plate reader (PerkinElmer) according to the manufacturer’s instructions.

### Immunoblot analysis

Cells with genetic alteration (gene knockdown or overexpression) were collected by centrifugation (400g, 5 min). After PBS washes, protein was extracted using RIPA lysis buffer with sonication (8 pulses, 10% output energy) and centrifugation. The supernatants were normalized using a BCA assay kit (Thermo), mixed with 4× LDS sample buffer (Invitrogen), then resolved by SDS-PAGE, transferred to PVDF membrane using a Mini Trans-Blot® Cell (Bio-Rad). The membrane was blocked with 5% milk in TBST buffer (20 mM Tris-HCl pH 7.5, 150 mM NaCl, 0.1% Tween 20) for 1 h at room temperature. The blot was then incubated with primary antibodies overnight at 4 °C followed with TBST washes and secondary antibody incubation for 1 h at room temperature. After three TBST washes, the blot was incubated with chemiluminescent HRP substrates (Thermo) and images were acquired on a Bio-Rad ChemiDoc imaging system.

### Genome-wide CRISPRi screens

The human CRISPRi sgRNA library Dolcetto in the XPR_050 backbone was used for the pooled screens. On day 0, 1.5 × 10^8 NCIH520-pZR071 cells stably expressing dCas9–KRAB were seeded in 12-well plates at 2 × 10^6 cells/well and infected with viral supernatant containing 8 μg/ml polybrene. Spin-infection was performed at 900 × g for 2 h. One day post-infection, cells were transferred to 15-cm dishes. On day 2, puromycin selection was initiated to enrich for sgRNA-transduced cells; the infection rate was ∼45% to have an optimal multiplicity of infection and about 1,000 cells per sgRNA. On day 6, cells were split into treatment arms and exposed to 2 μM probes or DMSO control. Cells were maintained for an additional 10-20 days, with media changes every 3-4 days including fresh 2 μM probe. Genomic DNA was isolated from cell pellets collected on day 16 and day 26 using NucleoSpin Blood L kits (Macherey-Nagel) following the manufacturer’s instructions and quantified with the PicoGreen dsDNA Assay Kit (Thermo Fisher).

### Library amplification and sequencing

For each PCR, 5 μg gDNA was combined with 50 μl Q5 Hot Start High-Fidelity 2× Master Mix (NEB), 5 μl each of 10 μM forward and reverse primers bearing Illumina adapters and indexes, and nuclease-free water to 100 μl total. Thermocycling: 98 °C 3 min; 28 cycles of 98 °C 20 s, 53 °C 25 s, 72 °C 25 s; final extension 72 °C 2 min. PCR products were pooled and purified with Mag-Bind® TotalPure NGS beads (Omega Bio-tek). Sequencing was performed on the Element Biosciences AVITI™ platform (single-end) with a 20% PhiX spike-in. sgRNA counts were obtained with PoolQ (Broad Institute) and normalized to counts per million (CPM). Gene-level enrichment/depletion significance was assessed using a hypergeometric test and log2 fold changes were computed by averaging fold changes across sgRNAs targeting the same gene.

### Enhanced cross-linking and immunoprecipitation (eCLIP)

The step-by-step eCLIP protocol has been described in detail and was followed in this study without change^43^. The resulting sequencing files were analyzed and plotted using Skipper^74,81^.

### RNA sequencing (RNAseq) and splicing analysis

Cells were plated in 6-well plates and treated with the indicated compounds for the indicated durations. At collection, medium was decanted and TRIzol™ (Thermo Fisher) was added to extract RNA, followed by column purification (Zymo). Total RNA was depleted of rRNA using the NEBNext® rRNA Depletion Kit v2 and libraries were prepared with the NEBNext® Ultra™ II Directional RNA Library Prep Kit according to the manufacturer’s instructions. Sequencing was performed on the Element Biosciences AVITI™ platform (paired-end 2 × 84 bp), targeting ∼80 million reads per sample. For expression analysis, transcript-level quantification was performed with Salmon (v1.10.3) using GENCODE v38 annotation; gene-level counts were derived with tximeta (v1.8.4). Differential expression was assessed with DESeq2 (v1.30.1); log2 fold changes were adjusted using apeglm v1.12.0, and P values were adjusted by the Benjamini–Hochberg procedure. For alternative splicing analysis, reads were aligned to hg38 with STAR, and splicing events were identified using rMATS v3.2.5. Finally, rmats2sashimiplot was used to generate Sashimi plots.

### Quantification and statistical analysis

Statistical analyses were performed using R and Python. Data visualization was carried out in R, Python, and GraphPad Prism 9. Comparisons between two groups were made using two-sided Student’s t-tests to calculate p-values, and two-sided statistical tests were applied throughout unless otherwise specified.

---

**Figure S1.**
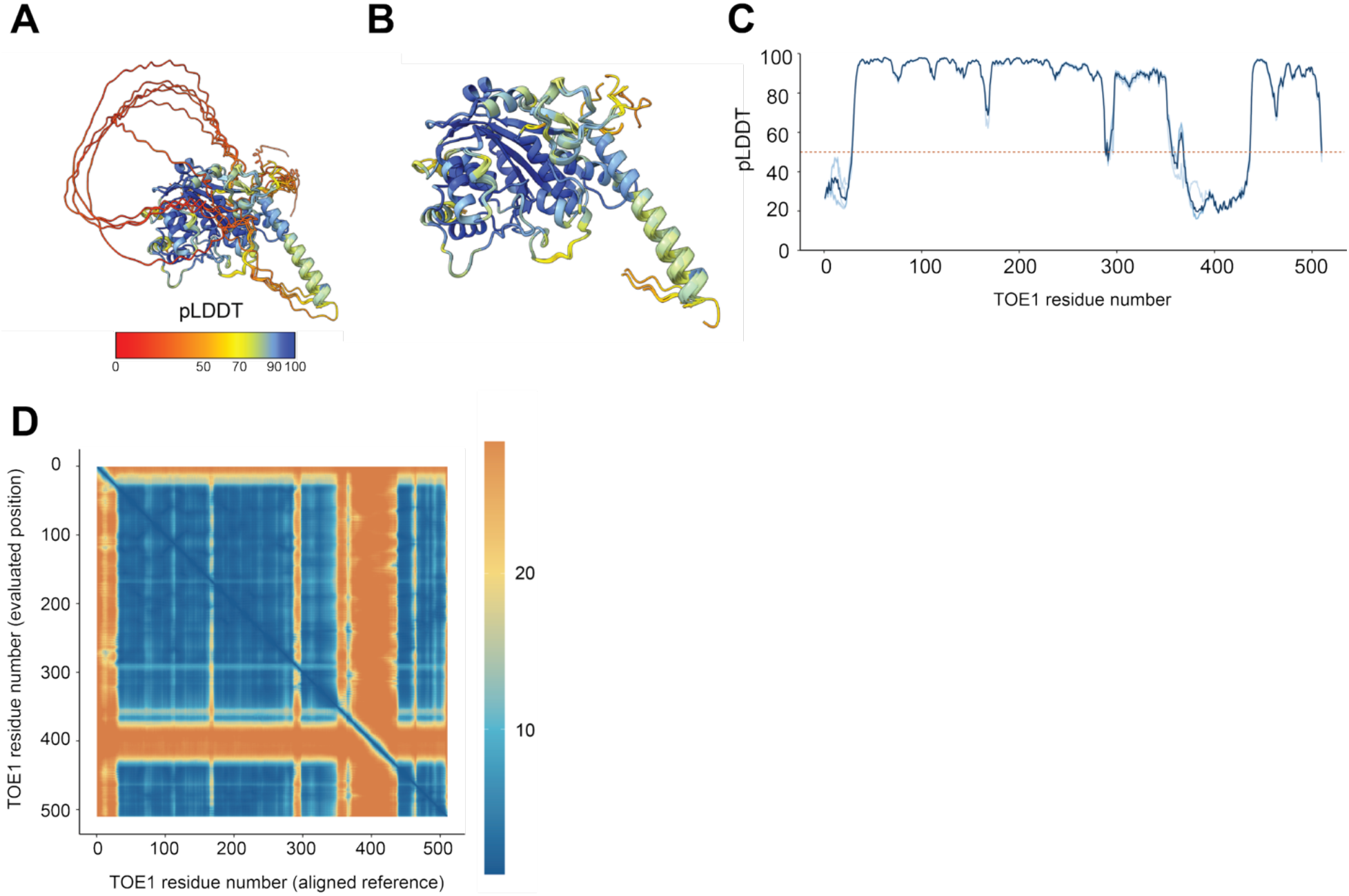
AlphaFold3 prediction of the TOE1 structure and assessment of model confidence. **(A)** Ensemble of five structural predictions generated by AlphaFold3^28^ with the amino acid sequence of human TOE1 (Uniprot: Q96GM8). Models are colored according to their predicted local distance difference test (pLDDT) scores. **(B)** Same ensemble as in (A), following removal of residues with pLDDT scores below 50 and remaining contiguous peptide fragments shorter than eight residues. **(C)** Per-residue pLDDT profiles of the five AF3 predictions (thin lines), with the mean pLDDT across all five models shown as a thick line. **(D)** Predicted aligned error (PAE) heatmap of the AF3 model selected for subsequent structural analysis (model 1 of 5).

**Figure S2.**
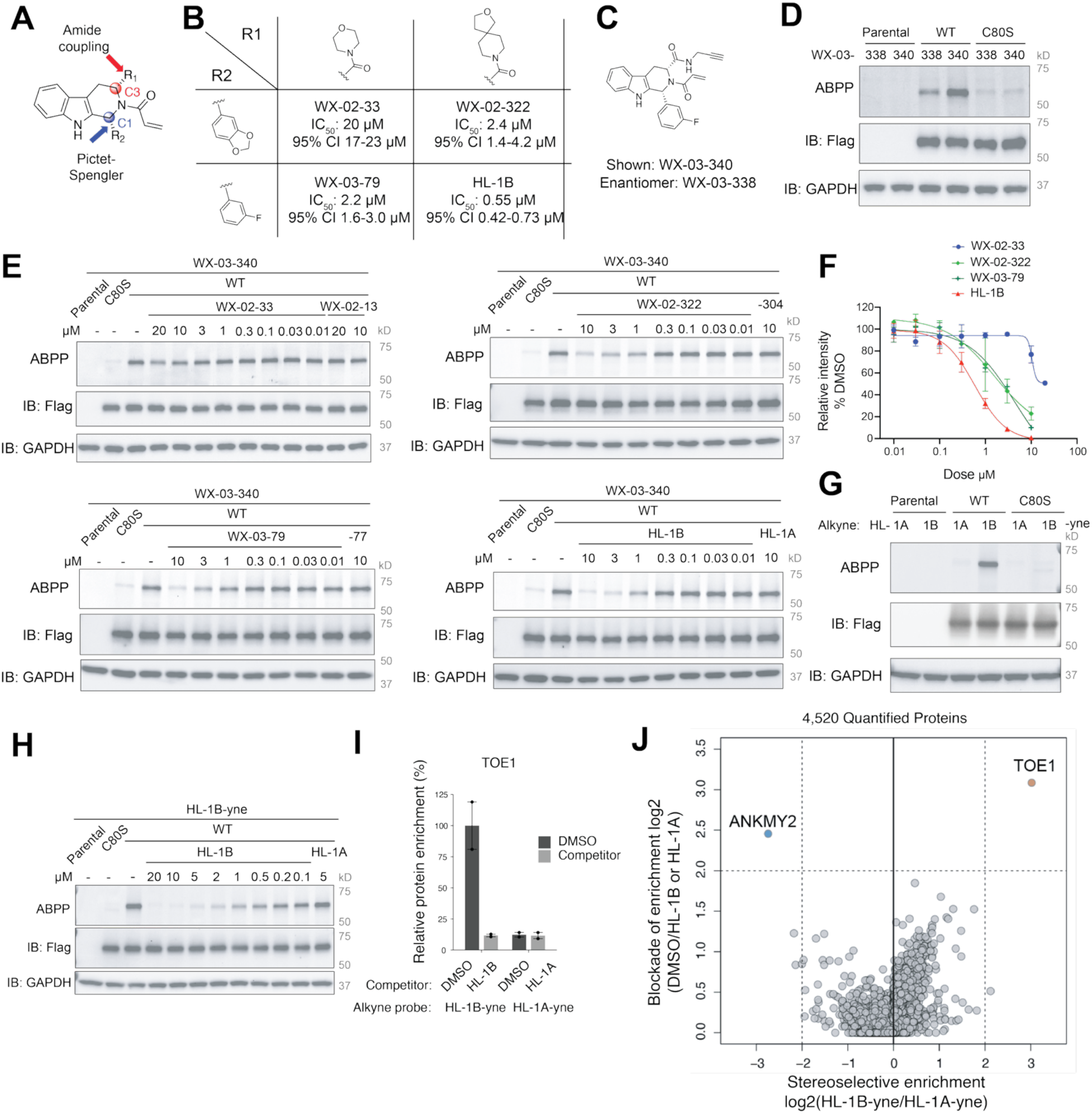
Optimization of tryptoline acrylamide stereoprobes targeting TOE1_C80. (A) General structure of tryptoline acrylamide stereoprobes showing C1 and C3 diversification points where groups appended to these sites are termed R2 and R1, respectively. (B) Parent tryptoline acrylamide structures and half-maximal values (IC_50_ values) for blockade of alkyne stereoprobe (WX-03-340) engagement of TOE1 as determined by gel-ABPP. WT-TOE1-expressing MCC142 cells were treated with the indicated tryptoline acrylamides for 3 h, followed by WX-03-340 (1 μM, 1 h) and gel-ABPP analysis. CI, confidence interval. Data are average values from 2 independent experiments. (C) Chemical structure of alkyne stereoprobe WX-03-340. (D) Gel-ABPP data showing stereoselective reactivity of WT-TOE1, but not C80S-TOE1 with alkyne stereoprobes WX-03-340 versus WX-03-338. Parental, WT-TOE1-, or C80S-TOE1-expressing MCC142 cells were treated with WX-03-340 or WX-03-338 (1 μM, 1 h) and samples were analyzed by gel-ABPP. Data are from a single experiment representative of 2 independent experiments. (E) Gel-ABPP data showing concentration-dependent blockade of WX-03-340 reactivity with WT-TOE1 by WX-02-33, WX-03-79, WX-02-322, or HL-1B in comparison to their respective enantiomers WX-02-13, WX-03-77, WX-02-304, or HL-1A. Parental, WT-TOE1-, or C80S-TOE1-expressing MCC142 cells were treated with parent compounds (3 h), followed by WX-03-340 (1 μM, 1 h) and gel-ABPP analysis. Data are from single experiments representative of 2 independent experiments. (F) Quantification of gel-ABPP data measuring engagement of TOE1_C80 by the indicated parent compounds. Data are average values ± SEM from 2 independent experiments. (G) Gel-ABPP data showing stereoselective reactivity of WT-TOE1, but not C80S-TOE1, with alkyne stereoprobes HL-1B-yne versus HL-1A-yne. Parental, WT-TOE1-, or C80S-TOE1-expressing MCC142 cells were treated with HL-1A-yne or HL-1B-yne (1 μM, 1 h) followed by gel-ABPP analysis. Data are from a single experiment representative of 2 independent experiments. (H) Gel-ABPP data showing concentration-dependent blockade of HL-1B-yne reactivity with WT-TOE1 by HL-1B. Parental, WT-TOE1-, or C80S-TOE1-expressing MCC142 cells were treated with HL-1A or HL-1B (3 h), followed by HL-1B-yne (1 μM, 1 h) and analysis by gel-ABPP. Data are from a single experiment representative of 2 independent experiments. (I) Protein-directed ABPP data showing stereoselective enrichment of endogenous TOE1 by HL-1B-yne and stereoselective blockade of this enrichment by HL-1B. Data are average values ± SEM from 2 independent experiments. (J) Scatter plot showing protein-directed ABPP data performed as follows: parental MCC142 cells were treated with DMSO or HL-1A or HL-1B (5 μM, 3 h), followed by stereomatched alkynes HL-1A-yne and HL-1B-yne (5 μM, 1 h), respectively. X axis: log2 protein enrichment ratios for HL-1B-yne versus HL-1A-yne from cells pre-treated with DMSO. Y axis: log2 ratios for blockade of protein enrichment by HL-1B (red) or HL-1A (blue). Data are average values from 2 independent experiments.

**Figure S3.**
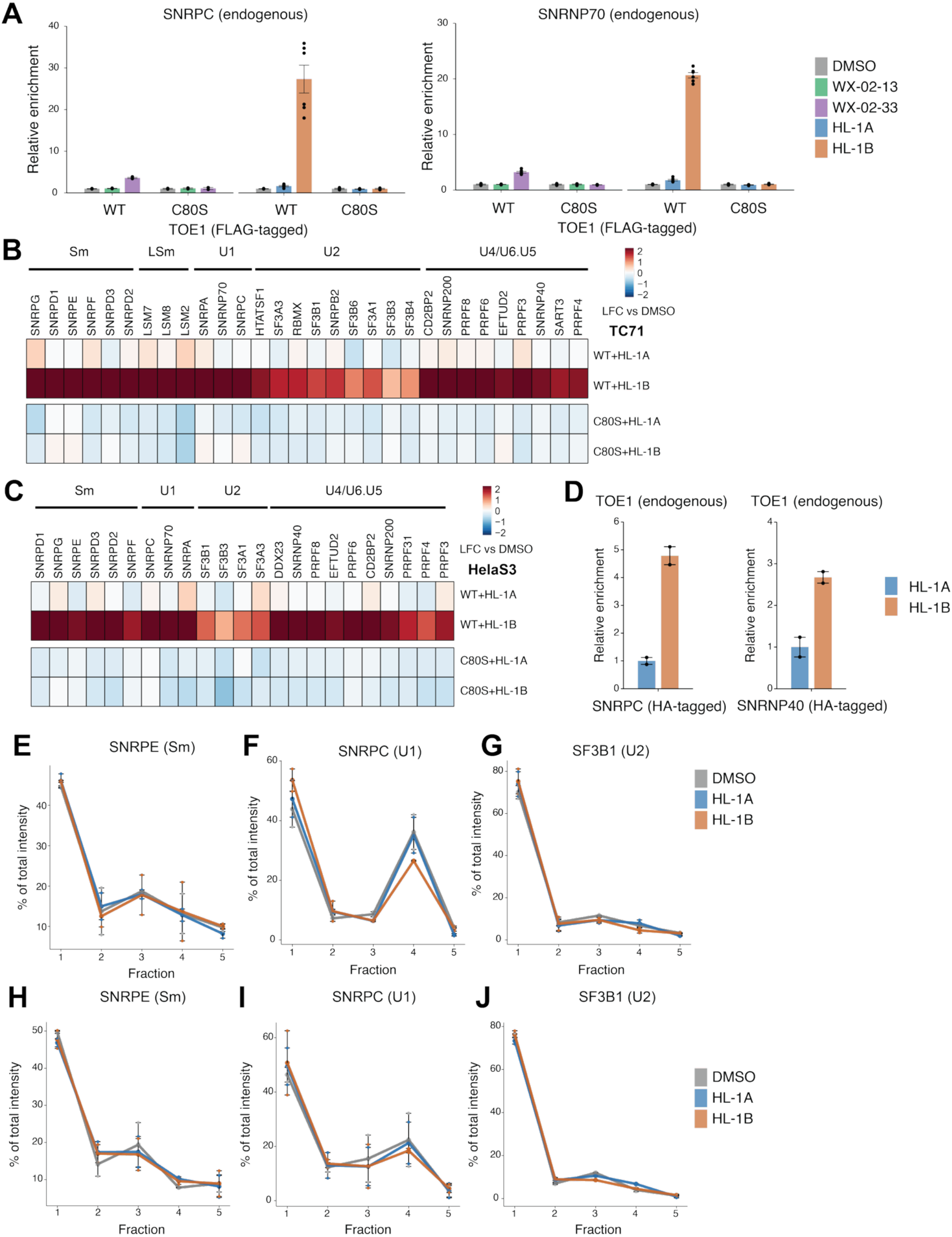
HL-1B promotes TOE1 binding to spliceosomal proteins in cells. (A) Barplot comparing the relative enrichment of endogenous SNRPC (left) and SNRNP70 (right) in anti-FLAG IP-MS experiments using FLAG-tagged WT-TOE1- or C80S-TOE1-expressing MCC142 cells treated with the original TOE1 probe WX-02-33 and its inactive enantiomer control WX-02-13 (20 μM each, 8 h) versus HL-1B and HL-1A (5 μM, 8 h). Data are average values ± SEM from 4-6 independent experiments. (B) Heatmap showing spliceosomal proteins (organized by spliceosomal sub-complexes) that were co-enriched in anti-FLAG IP-MS experiments from FLAG-tagged WT-TOE1- or C80S-TOE1-expressing TC71 cells treated with HL-1B or HL-1A (5 μM, 8 h) or DMSO. Data are average values from 4-6 independent experiments. (C) Heatmap showing spliceosomal proteins (organized by spliceosomal sub-complexes) that were co-enriched in anti-FLAG IP-MS experiments from FLAG-tagged WT-TOE1- or C80S-TOE1-expressing HeLa S3 cells treated with HL-1B or HL-1A (5 μM, 8 h) or DMSO. Data are average values from 4-6 independent experiments. (D) Barplot showing relative enrichment of endogenous TOE1 in anti-HA-SNRPC (left) or anti-HA-SNRNP40 (right) IP-MS experiments using MCC142 cells treated with HL-1A or HL-1B (5 μM, 8 h). Data are average values ± SEM from 2 independent experiments. (**E-G**) SEC-MS elution profiles of SNRPE (**E**), SNRPC (**F**), and SF3B1 (**G**) from WT-TOE1-expressing MCC142 cells. Cells were treated with HL-1A, HL-1B (5 μM, 8 h) or DMSO and clarified lysates were separated by FPLC and fractionated into five fractions before MS analysis. Data are average values ± SEM from 2 independent experiments. (**H-J**) SEC-MS elution profiles of SNRPE (**H**), SNRPC (**I**), and SF3B1 (**J**) from C80S-TOE1-expressing MCC142 cells. Cells were treated with HL-1A, HL-1B (5 μM, 8 h) or DMSO and clarified lysates were separated by FPLC and fractionated into five fractions before MS analysis. Data are average values ± SEM from 2 independent experiments.

**Figure S4.**
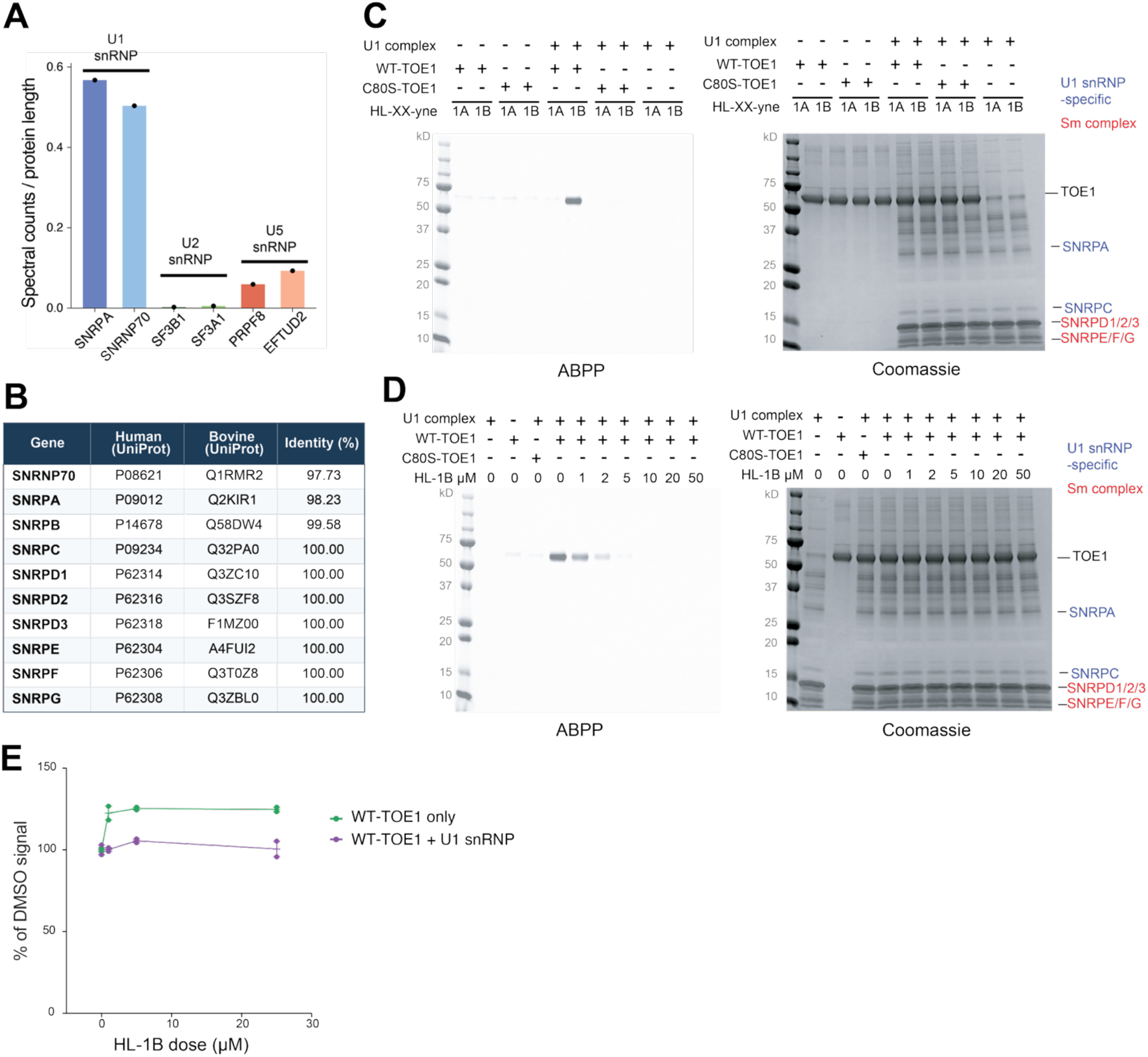
HL-1B reactivity with TOE1 requires the presence of U1 snRNP proteins. (A) Proteomic characterization of a commercial purified bovine snRNP preparation obtained from Surmodics IVD. Spectral counts for each protein were normalized to the protein length to provide a semi-quantitative estimate of relative protein abundances, which are shown for unique members of the U1, U2, and U5 snRNP complexes. (B) Summary of human and bovine U1 snRNP protein sequence alignment showing very high overall sequence identity. (C) Left, gel-ABPP data showing the stereoselective and site-selective engagement of purified His-WT-TOE1 by HL-1B-yne (2 μM, 1 h) in the presence of a purified bovine U1 snRNP complex using the indicated alkyne stereoprobes. Right, Coomassie blue protein staining. Data are from a single experiment representative of 2 independent experiments. (D) Left, gel-ABPP data showing the stereoselective and concentration-dependent blockade of HL-1B-yne (2 µM, 1 h) reactivity with purified His-WT-TOE1 by HL-1B (1 h) in the presence of a purified bovine U1 snRNP complex. Right, Coomassie blue protein staining. Data are from a single experiment representative of 2 independent experiments. (E) MS-based proteomic monitoring of TOE1 tryptic peptides other than the peptide containing TOE1_C80. Data are from the experiment described in Figure 2B and are average values ± SEM normalized to DMSO from 2 independent experiments.

**Figure S5.**
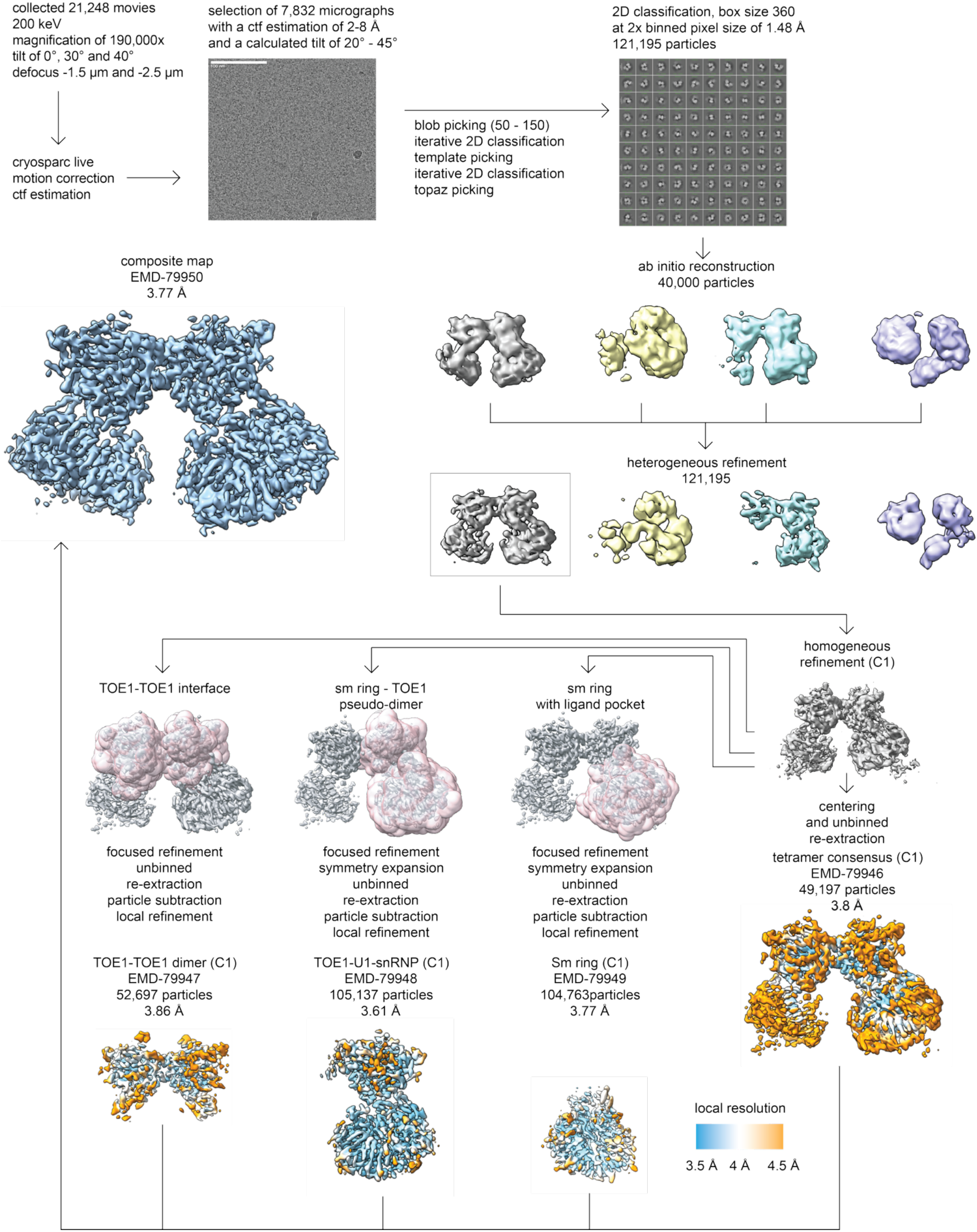
Cryo-EM data processing workflow for the TOE1–HL-1B–U1 snRNP complex. 21,248 movies were collected at 200 kV with stage tilts of 0°, 30°, and 40°. Following motion correction and CTF estimation in cryoSPARC, 7,832 micrographs were selected for particle picking using blob picking, template picking, and Topaz, followed by iterative 2D classification. A total of 121,195 particles were subjected to *ab initio* reconstruction and heterogeneous refinement. Subsequent homogeneous refinement without imposed symmetry (C1) yielded a consensus reconstruction of the TOE1–U1 snRNP assembly. Focused refinements incorporating particle re-extraction, subtraction, and symmetry expansion where indicated produced reconstructions of the complete assembly (3.8 Å; 49,197 particles), TOE1 dimer interface (3.86 Å; 52,697 particles), TOE1–U1 snRNP subcomplex (3.61 Å; 105,137 particles), and Sm ring containing the HL-1B binding pocket (3.77 Å; 104,763 particles). A composite map was generated at 3.7 Å resolution. Local resolution distributions are shown for the final focused reconstructions.

**Figure S6.**
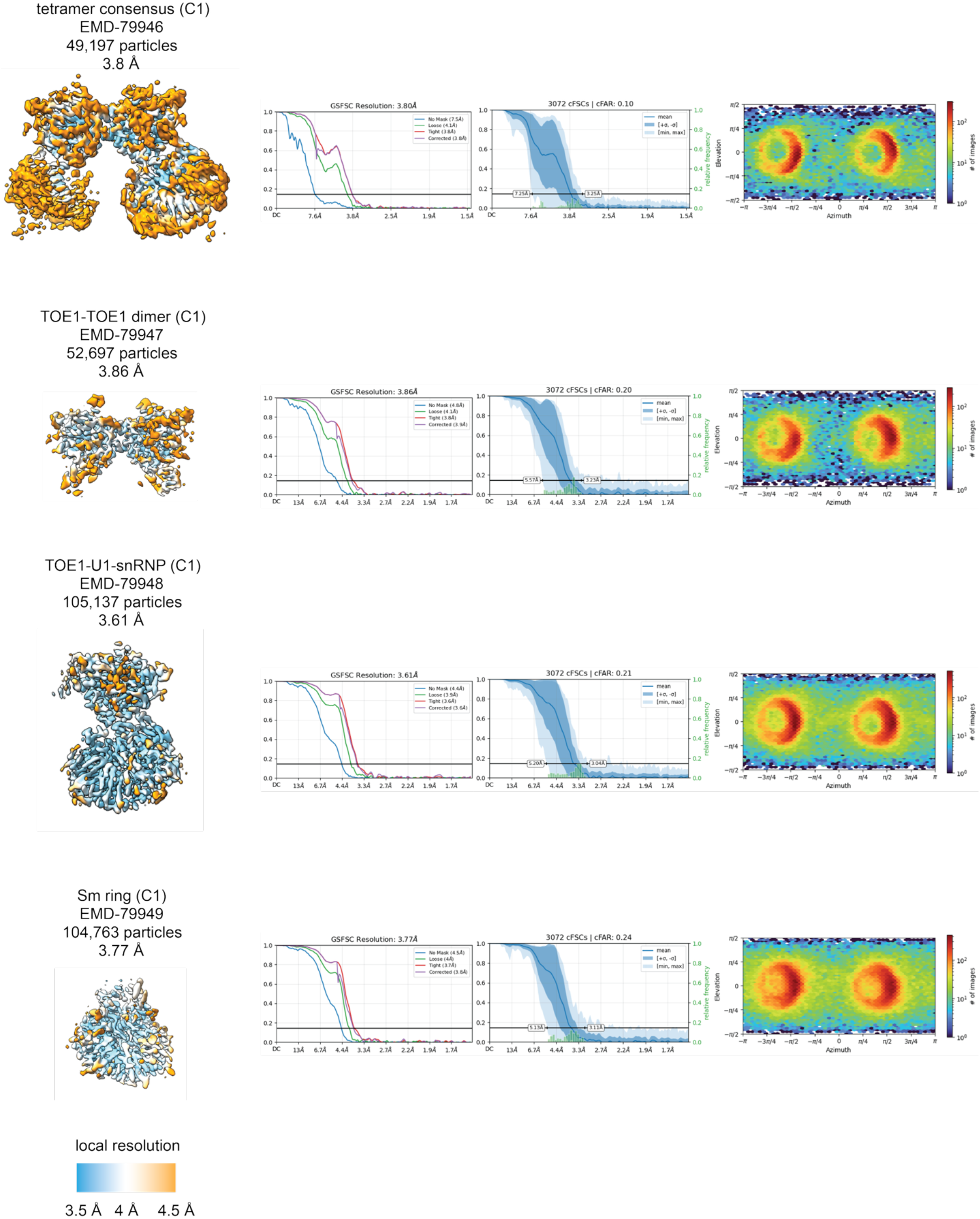
Cryo-EM reconstruction quality and validation of the TOE1–HL-1B–U1 snRNP complex. Cryo-EM maps, Fourier shell correlation (FSC) curves, and angular particle distributions for the four reconstructions described in **Figure S5**: complete assembly (3.80 Å), TOE1–TOE1 dimer interface (3.86 Å), TOE1–U1 snRNP subcomplex (3.61 Å), and Sm ring containing the HL-1B binding pocket (3.77 Å), shown from top to bottom. Maps are colored by local resolution. FSC plots show resolution estimates using the 0.143 criterion, and angular distribution plots illustrate particle orientations contributing to each reconstruction.

**Figure S7.**
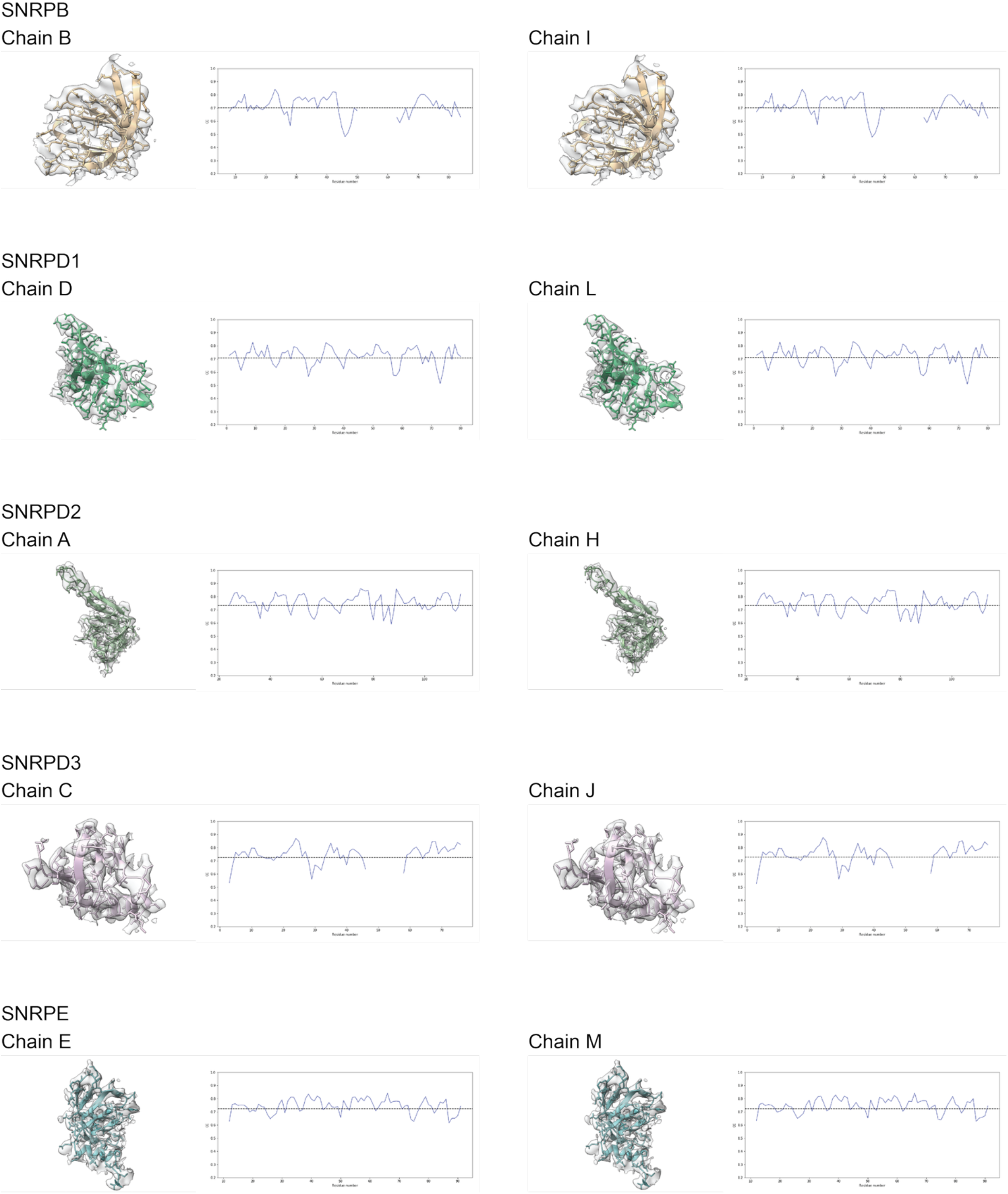
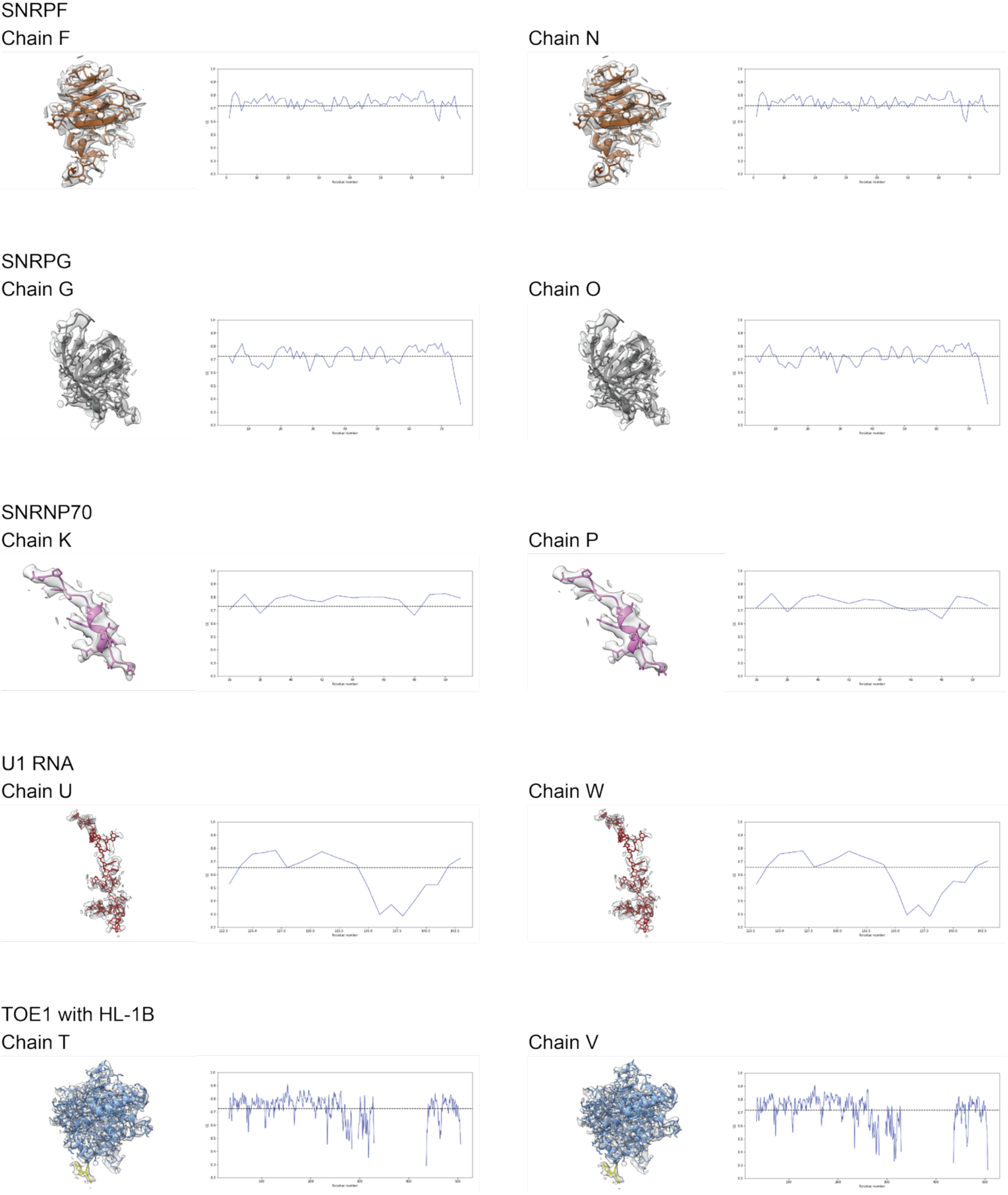
Cryo-EM density and model-to-map correlation for individual components of the TOE1–HL-1B–U1 snRNP complex. Representative views of the cryo-EM density (gray) with fitted atomic models (colored ribbons or sticks) for TOE1 and its covalently bound HL-1B adduct, U1 snRNA, SNRNP70, and the seven Sm ring proteins (SNRPB, SNRPD1, SNRPD2, SNRPD3, SNRPE, SNRPF, and SNRPG). Each component is shown for both copies of the complex, with chain identifiers indicated. Corresponding plots show local model-to-map correlation coefficients (CC) as a function of residue or nucleotide position, providing a measure of agreement between the atomic models and experimental density.

**Figure S8.**
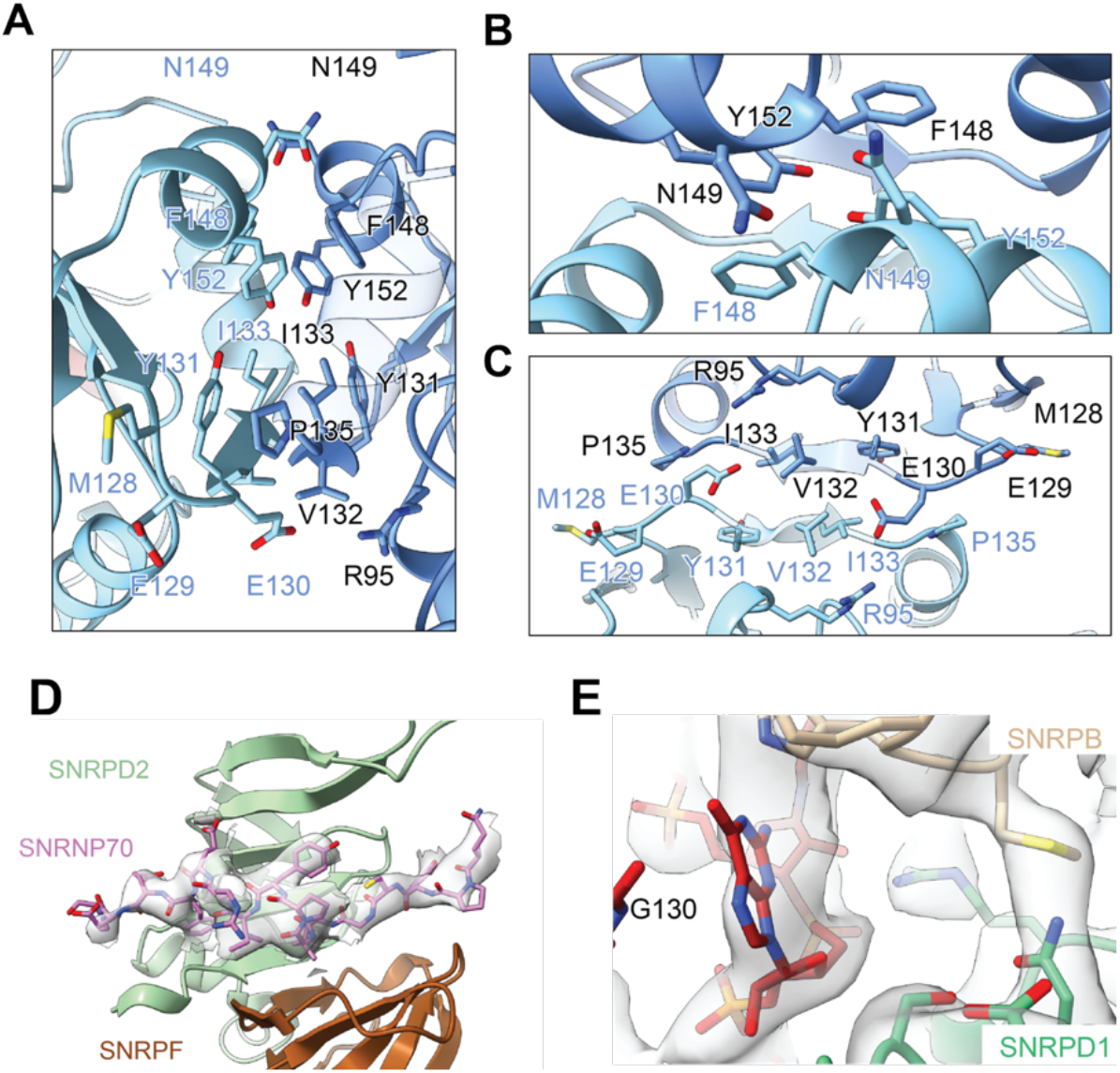
Structural details of the TOE1 dimer interface and identification of U1 snRNP. **(A–C)** Close-up views of the TOE1 dimer interface from the side (A), top (B), and bottom (C). The two TOE1 protomers are shown in light and dark blue, with interacting residues displayed as sticks and labeled. The interface is formed primarily by hydrophobic packing involving M128, Y131, V132, I133, P135, F148, and Y152, with additional contributions from R95, E129, E130, and N149. The twofold symmetry axis is indicated (C_2_). **(D)** Cryo-EM density corresponding to a segment of SNRNP70, a U1-specific protein, associated with the Sm ring. SNRNP70 is shown in pink, with neighboring Sm proteins colored as indicated. **(E)** Cryo-EM density surrounding G130 of U1 snRNA within the Sm-binding motif, showing its *syn* conformation. The presence of guanosine at this position distinguishes U1 from other candidate U snRNAs containing uridine at the corresponding position. Together with the identification of SNRNP70 in (D), this supports assignment of the complex as TOE1–U1 snRNP.

**Figure S9.**
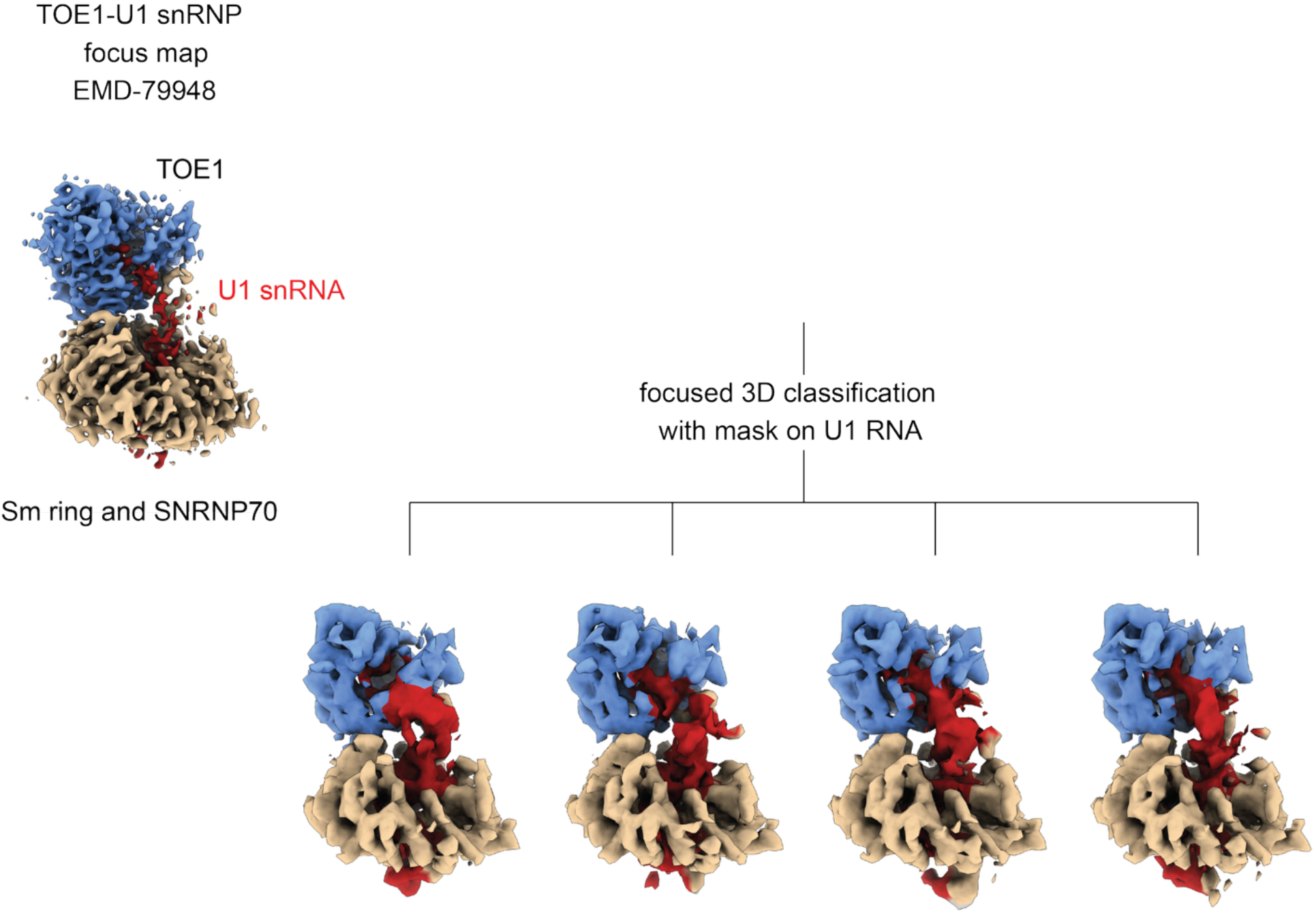
Focused classification improves the continuity of U1 snRNA density between the Sm ring and the TOE1 active site. Focused 3D classification of the TOE1–U1 snRNP reconstruction (EMD-79948), using a mask encompassing U1 snRNA, yielded four classes with improved continuity of RNA density between the Sm-binding region and the TOE1 active site. Although the precise RNA backbone path and sequence register remain uncertain, the continuous density supports an extended RNA configuration rather than formation of a stable stem-loop structure. TOE1, U1 snRNA, and the Sm ring are shown in blue, red, and tan, respectively. The continuous density supports an extended RNA configuration bridging the Sm ring and TOE1 active site, rather than formation of a stable stem-loop structure within the intervening region.

**Figure S10.**
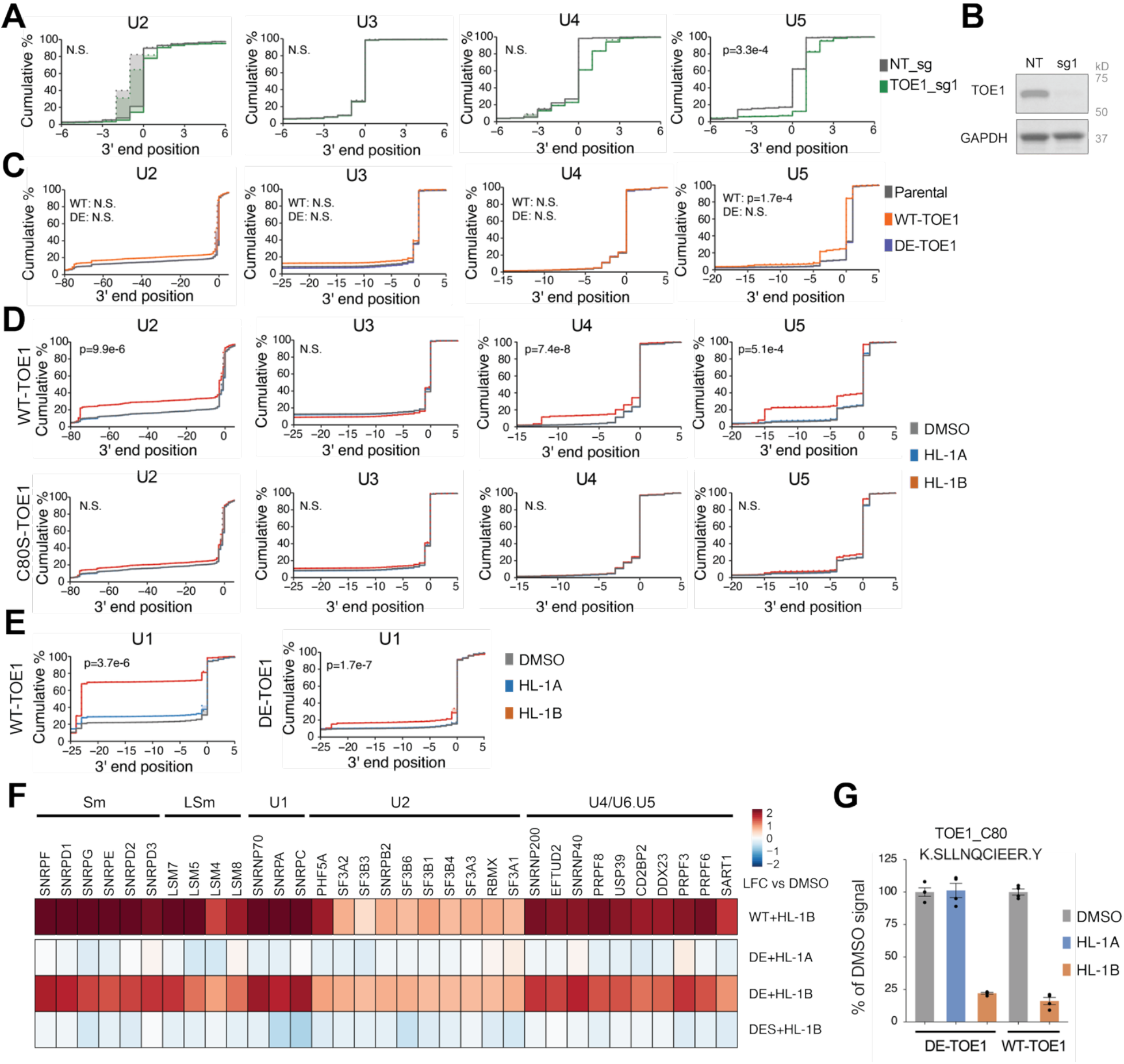
HL-1B promotes TOE1-mediated over-trimming of snRNAs in cells. (A) Cumulative plots comparing 3′ end position distributions of U2, U4, and U5 snRNAs and U3 snoRNA in NCIH520 cells expressing a TOE1-targeting CRISPRi sgRNA (TOE1_sg1) or non-targeting (NT) control sgRNA (NT_sg). (B) Western blot showing TOE1 expression in NCIH520-dCas9-KRAB cells treated with non-targeting (NT) or TOE1-targeting CRISPRi sgRNA (sg1) for 7 days. Results are from a single experiment representative of 2 independent experiments. (C) Cumulative plots comparing 3′ end position distributions of the U2, U4, U5 snRNAs and U3 snoRNA in parental MCC142 cells versus WT-expressing- or DE-TOE1 mutant-expressing MCC142 cells. DE, D64A+E66A, catalytically inactive TOE1 mutant. (D) Cumulative plots comparing 3′ end position distributions of U2, U4, and U5 snRNAs and U3 snoRNA in WT-TOE1- and C80S-TOE1-expressing MCC142 cells treated with HL-1A or HL-1B (2 μM, 3 days) or DMSO. (E) Cumulative plots comparing 3′ end position distributions of the U1 snRNA in WT- or DE-TOE1-expressing MCC142 cells treated with HL-1A or HL-1B (2 μM, 3 days) or DMSO. (F) Heatmap showing proteins that were co-enriched in anti-FLAG IP-MS experiments from FLAG-tagged WT-TOE1-, DE-TOE1-, or DES-TOE1-expressing MCC142 cells treated with HL-1B or HL-1A (5 μM, 8 h) or DMSO. Results are average values of 4 independent experiments. DE, D64A+E66A double TOE1 mutant; DES, D64A+E66A+C80S triple TOE1 mutant. (G) Quantification of TOE1_C80 engagement by the indicated compounds (5 μM, 8 h) as measured by monitoring the C80-containing tryptic peptide in IP-MS experiments performed in (**F**) Data are average values ± SEM normalized to DMSO from 4 independent experiments. For **A**, **C**, **D**, **E**, data are average values from 3 independent experiments of gene-specific 3′ end sequencing. For **A**, the p values indicate comparison between TOE1_sg1 and NT_sg groups. For **C**, the p values indicate comparison between WT-TOE1 or DE-TOE1-expressing vs parental groups. For **D** and **E**, the p values indicate comparisons between HL-1B- vs HL-1A-treated groups. Not significant (N.S.) is defined as p>0.05 or mean difference<1 nucleotide. All p values were calculated using two-sided Student’s t test.

**Figure S11.**
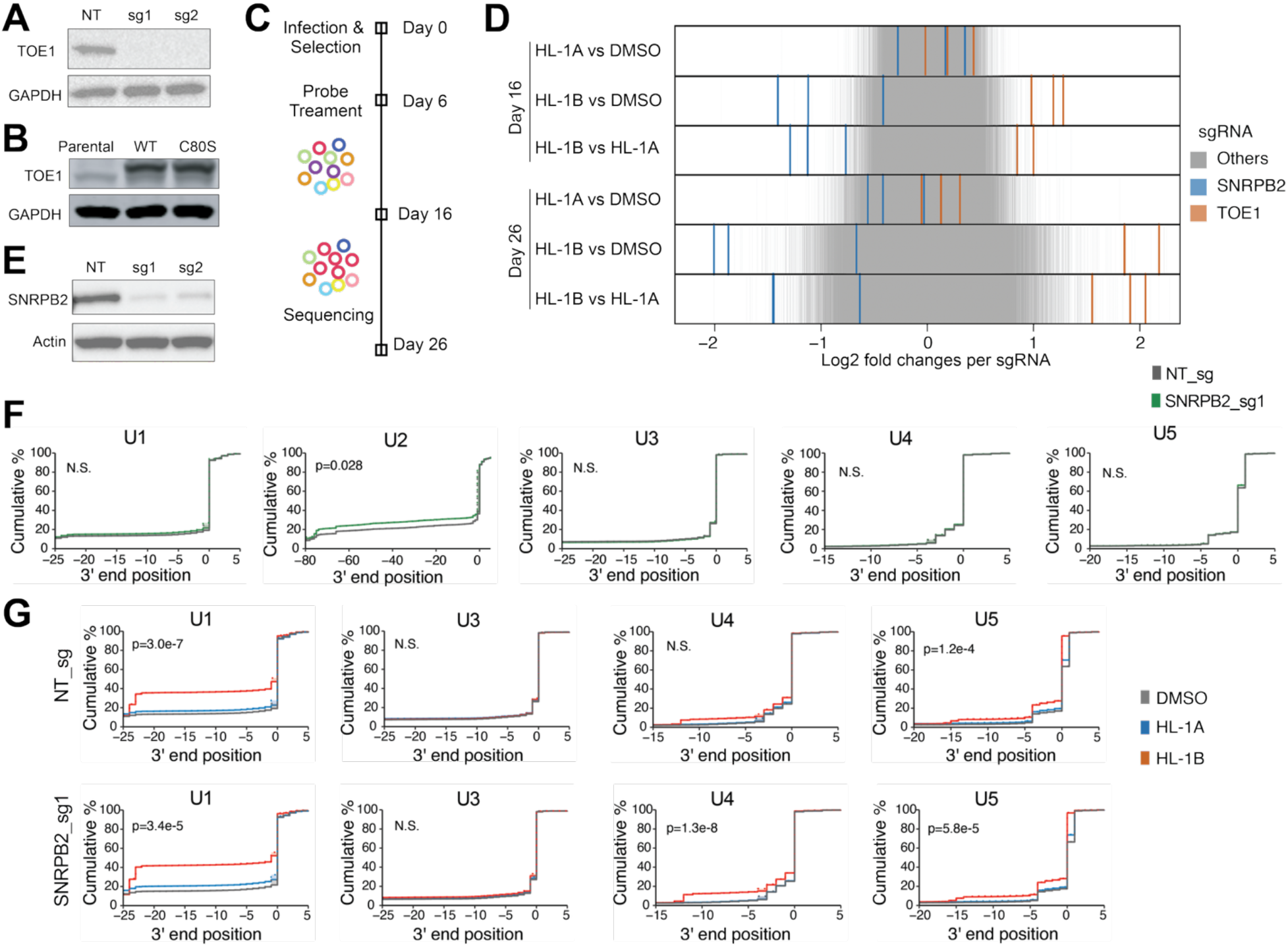
Genetic modifiers of the cancer cell growth effects of HL-1B. (A) Western blot showing TOE1 expression in NCIH520-dCas9-KRAB cells treated with non-targeting (NT) or TOE1-targeting CRISPRi sgRNAs (sg1, sg2) for 7 days. Results are from a single experiment representative of 2 independent experiments. (B) Western blot showing TOE1 expression in parental, WT-TOE1-expressing, and C80S-TOE1-expressing MCC142 cells. Results are from a single experiment representative of 2 independent experiments. (C) Schematic for the workflow of genome-wide CRISPRi screens for identifying genes that confer sensitization or resistance to growth effects of HL-1B. NCIH520 cells expressing dCas9-KRAB were infected with a sgRNA library and then grown in the presence of HL-1A or HL-1B (2 μM) for 10-20 days before sequencing. (D) Enrichment of sgRNAs targeting SNRPB2 (blue) or TOE1 (red) in comparison to the distribution of all other sgRNAs (grey). Each line represents an sgRNA. Data are shown for cells collected on Day 16 (top) and Day 26 (bottom) from the workflow diagram in (**C**). (E) Western blot showing SNRPB2 expression in NCIH520-dCas9-KRAB cells with non-targeting sgRNA (NT) or SNRPB2-targeting CRISPRi sgRNAs (sg1, sg2) for 7 days. Results are from a single experiment representative of 2 independent experiments. (F) Cumulative plots comparing 3′ end position distributions of the U1, U2, U4, U5 snRNAs and U3 snoRNA in NCIH520 cells expressing a SNRPB2-targeting CRISPRi sgRNA (SNRPB2_sg1) or non-targeting (NT) control sgRNA (NT_sg). (G) Cumulative plots comparing 3′ end position distributions of the U1, U4, U5 snRNAs and U3 snoRNA in SNRPB2_sg1 and control NT_sg NCIH520 cells treated with HL-1A or HL-1B (2 μM, 3 days) or DMSO. For **F** and **G**, data are average values from 3 independent experiments of gene-specific 3′ end sequencing. For **F**, the p values indicate comparison between SNRPB2_sg1 and NT_sg groups. For **G**, the p values indicate comparisons between HL-1B- vs HL-1A-treated groups. Not significant (N.S.) is defined as p>0.05 or mean difference < 1 nucleotide. All p values were calculated using two-sided Student’s t test.

**Figure S12.**
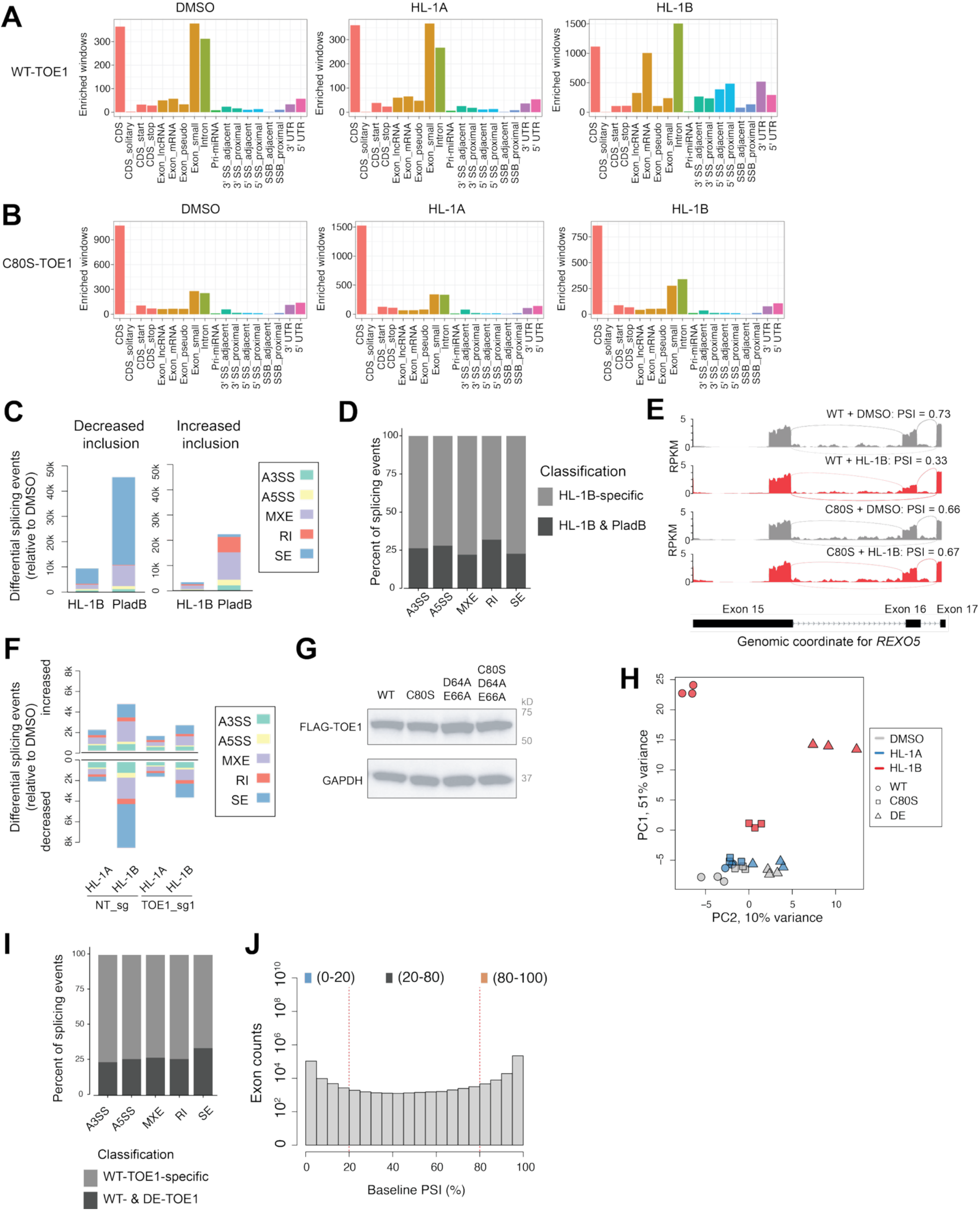
Further characterization of HL-1B effects on alternative splicing. (**A** and **B**) TOE1 eCLIP-seq binding profiles in WT-TOE1-expressing (**A**) or C80S-TOE1-expressing (**B**) MCC142 cells treated with HL-1A or HL-1B (2 μM, 24 h) or DMSO. CDS, coding sequence; UTR, untranslated region; SS, splice site. Proximal denotes within 500 bases and adjacent denotes within 100 bases from the annotated splice sites. For all eCLIP experiments in this figure, data are derived from 2 independent experiments. (C) Summary of alternative splicing events induced by pladB (20 nM, 24 h) or HL-1B (2 μM, 24 h) versus DMSO in WT-TOE1-expressing MCC142 cells, as quantified by RNAseq and rMATS analysis. Significant events defined as having differential (DMSO vs probe) percent spliced in (ΔPSI) values of > 0.1 or < -0.1 and FDR < 0.05. Data are significant events from 3 independent experiments. A3SS, alternative 3′ splice site; A5SS, alternative 5′ splice site; MXE, mutually exclusive exons; RI, retained intron; SE, skipped exons. Inclusion forms are defined as: SE, target exon included; MXE, 5′ exon; A3SS/A5SS, longer exon; RI, intron retained. (D) Subclassification of HL-1B-induced splicing changes in WT-TOE1-expressing MCC142 cells divided into events restricted to HL-1B versus events observed in both HL-1B- and pladB-treated cells. (E) Sashimi plots of REXO5 splicing in WT-TOE1- and C80S-TOE1-expressing MCC142 cells treated with HL-1B (2 μM, 3 days) or DMSO. PSI, percent spliced in. (F) Summary of alternative splicing events induced by HL-1A or HL-1B (2 μM, 24 h) versus DMSO in NT_sg or TOE1_sg1 NCIH520 cells. Data for significantly increased or decreased splicing events are shown in upper and lower portion of graph, respectively. (G) Western blot showing TOE1 expression in WT- or the indicated mutant TOE1-expressing MCC142 cells. Results are from a single experiment representative of 2 independent experiments. (H) PCA plot of RNA-seq data for the indicated groups, showing that HL-1B (2 μM, 3 days) induces much greater stereoselective gene expression effects in WT-TOE1-expressing vs C80S-TOE1- or DE-TOE1-expressing MCC142 cells. Data from cells treated with DMSO or HL-1A (2 μM, 3 days) are also shown. DE, D64A+E66A, catalytically inactive mutant. (I) Subclassification of HL-1B-induced (2 μM, 3 days) splicing changes in WT-TOE1-expressing MCC142 cells divided into events specific to WT-TOE1-expression versus events observed in both WT-TOE1 and DE-TOE1-expressing cells. (J) Histogram showing the distribution of exon counts across bins of baseline PSI levels.

**Figure S13.**
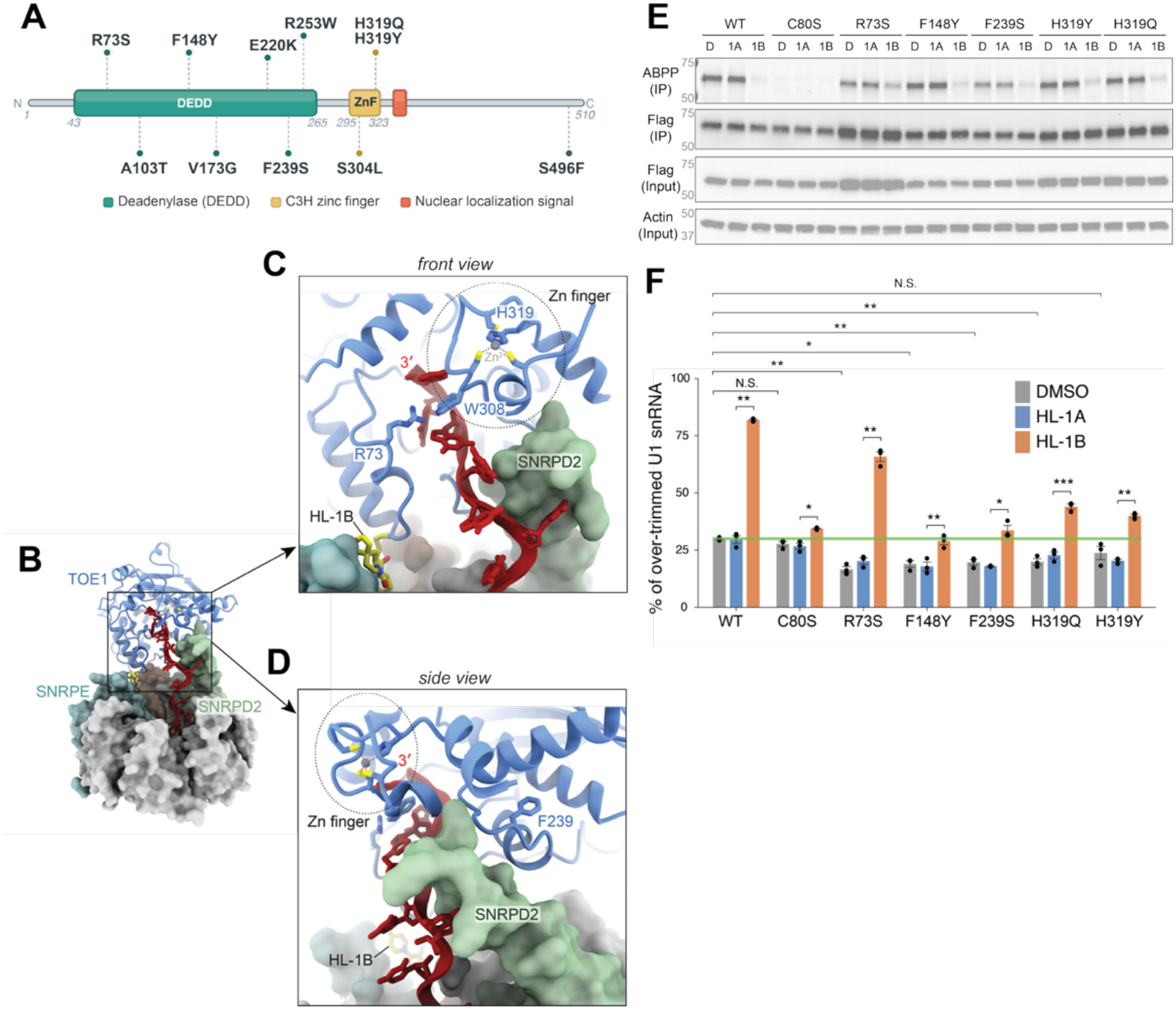
Characterization of pathogenic TOE1 mutants and their sensitivity to HL-1B. (A) The domain architecture of TOE1 marking the location of reported PCH7-associated mutations. The deadenylase domain is colored in green, the C3H zinc finger in yellow, and the nuclear localization signal (NLS) in orange. (B) Overview of the TOE1–HL-1B–U1 snRNP complex, with TOE1 shown as a blue cartoon, U1 snRNA in red, HL-1B in yellow, and the Sm ring as a molecular surface. SNRPD2 and SNRPE are highlighted in green and teal, respectively. Boxes indicate general region enlarged in panels C and D. (C) Front view of the TOE1 active site, highlighting residues R73 and H319 with PCH7-associated mutations. H319 coordinates a Zn^2+^ ion within a zinc finger containing W308, which stacks against the 3′-terminal nucleotides of U1 snRNA. R73 is positioned near W308 and may contribute to substrate positioning. (D) Side view highlighting F239 (another TOE1 residue mutated in PCH7), which makes hydrophobic packing interactions with the TOE1 interior, stabilizing a TOE1 surface loop that directly contacts SNRPD2. The interaction suggests a potential role for F239 in TOE1 engagement with the Sm ring. (E) Gel-ABPP data showing blockade of HL-1B-yne reactivity with WT-TOE1 and PCH7-associated TOE1 mutants by HL-1B. TOE1 variant-expressing MCC142 cells were treated with HL-1A or HL-1B (3 h), followed by HL-1B-yne (1 μM, 1 h) and analysis by gel-ABPP. Endogenous TOE1 levels were reduced in these cell models using dCas9-KRAB paired with TOE1_sg1. Data are from a single experiment representative of 2 independent experiments. (F) Barplots comparing the 3′ tail lengths of the U1 snRNA in MCC142 cells expressing the indicated TOE1 variants and treated with HL-1A or HL-1B (2 μM, 3 days) or DMSO. Endogenous TOE1 levels were reduced in these cell models by using dCas9-KRAB paired with TOE1_sg1. The % over-trimming values were calculated as the fraction of U1 snRNA with 3′ end ≥1 over-trimmed nucleotide from the canonical 3′ end position. Data are average values ± SEM from 3 independent experiments. All p values were calculated using two-sided paired Student’s t test. N.S., not significant, * p<0.05, **p<0.01, ***p<0.001.

## Notes

### Competing Interest Statement

The authors have declared no competing interest.

### Summary of Updates

A supplementary information file has been added that contains information on the synthesis and characterization of novel compounds

