## Supplementary material for "A Molecular Glue that Regulates Processing of Non-Coding RNAs in Human Cells": Table S2

**Cryo-EM data collection, refinement, and validation statistics**

|  | Consensus map of U1-TOE1-TOE1-U1 pseudo-tetramer  (EMD-79946) | Focus refinement of TOE1-TOE1 dimer  (EMD-79947) | Focus refinement of TOE1-U1 pseudo-dimer (EMD-79948) | Focus refinement of U1 snRNP (EMD-79949) | Composite map and model of the U1-TOE1-TOE1-U1 complex  (EMD-79950)  (PDB 40BP) |
| --- | --- | --- | --- | --- | --- |
| **Data collection and processing** |  |  |  |  |  |
| Magnification | 190,000 | 190,000 | 190,000 | 190,000 | 190,000 |
| Voltage (kV) | 200 | 200 | 200 | 200 | 200 |
| Electron exposure (e^–^/Å^2^) | 40 | 40 | 40 | 40 | 40 |
| Defocus range (μm) | -1.5 to -2.5 | -1.5 to -2.5 | -1.5 to -2.5 | -1.5 to -2.5 | -1.5 to -2.5 |
| Pixel size (Å) | 0.74 | 0.74 | 0.74 | 0.74 | 0.74 |
| Symmetry imposed | C1 | C1 | C1 | C1 | C1 |
| Initial particle images (no.) | 121,195 | 121,195 | 121,195 | 121,195 | N/A |
| Final particle images (no.) | 49,197 | 52,697 | 105,137  (symmetry expanded) | 104,763  (symmetry expanded) | N/A |
| Map resolution (Å)  FSC threshold | 3.8  0.143 | 3.86  0.143 | 3.61  0.143 | 3.77  0.143 | 3.77  N/A |
| Map resolution range (Å) | 3.65-8.11 | 3.71-12.21 | 3.46-12.38 | 3.51-11.08 | N/A |
| **Refinement** |  |  |  |  |  |
| Model-to-map-fit, mask CC | N/A | N/A | N/A | N/A | 0.69 |
| Model composition  Non-hydrogen atoms  Protein residues  Nucleotide residues  Ligands |  |  |  |  | 15324  1796  42  4 |
| Mean Isotropic B factor (Å^2^)  Protein  Nucleotide  Ligand |  |  |  |  | 80.39  96.15  74.63 |
| R.m.s. deviations  Bond lengths (Å)  Bond angles (°) |  |  |  |  | 0.006  1.290 |
| Validation  MolProbity score  Clashscore  Poor rotamers (%) |  |  |  |  | 1.95  9.61  1.38 |
| Ramachandran plot  Favored (%)  Allowed (%)  Disallowed (%) |  |  |  |  | 95.17  4.71  0.11 |
