## Supplementary material for "A Molecular Glue that Regulates Processing of Non-Coding RNAs in Human Cells": Synthetic Chemistry Information

### Supplementary Information: Synthesis and Characterization of Novel Compounds

#### General considerations

All NMR spectra were recorded at 298 K unless otherwise noted. $^{1}$H NMR spectra were recorded on Bruker Avance III 400, Avance III HD 400, Avance Neo 400 spectrometers or a ZKNJ QOne Quantum-I Plus 400 spectrometer ($^{1}$H, 400 MHz). $^{1}$H NMR data are reported as follows: chemical shift (δ), multiplicity (s = singlet, d = doublet, t = triplet, m = multiplet; br. = broad), coupling constants, and integration. Chemical shifts are reported in parts per million (ppm) using the appropriate solvent as reference. Analytical supercritical fluid chromatography (SFC) was performed on a Shimadzu LC system (flow rate: 3 mL/min, back pressure: 100 Bar, column temperature: 35 ºC) equipped with a polydiode array detector. Tandem liquid chromatography/mass spectrometry (LC-MS) was performed on an Agilent 1200/1260 series LC/MSD system equipped with an Agilent G6110A mass detector, alternatively a Shimadzu LC-20AD or AB series LC-MS system equipped with Shimadzu SPD-M20A or SPDM40 mass detectors. Mass measurements for high-resolution mass spectrometry (HRMS) were performed on a Waters Xevo G2-XS TOF calibrated against sodium formate clusters and using a LeuEnk lockmass. Expected monoisotopic masses were calculated using MassLynx 4.1 and the m/z values for calibrant and lockmass were MassLynx-default values.

#### Experimental procedures and analytical data

Experimental procedures as well as analytical data for all new stereoprobes are provided.

*Synthesis of WX-02-322*

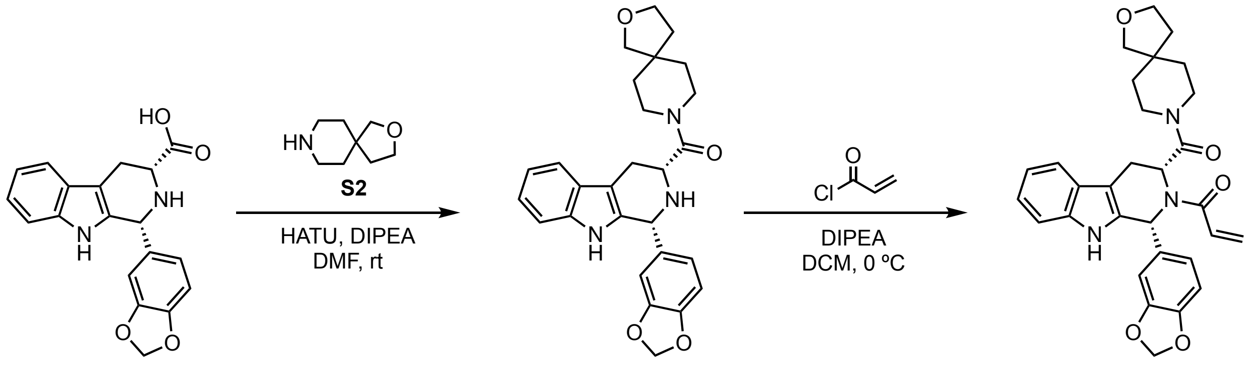


**S1** **S3** **WX-02-322**

1-((1*R*,3*R*)-1-(benzo[*d*][1,3]dioxol-5-yl)-3-(2-oxa-8-azaspiro[4.5]decane-8-carbonyl)-1,3,4,9-tetrahydro-2*H*-pyrido[3,4-*b*]indol-2-yl)prop-2-en-1-one (WX-02-322)
To a solution of S1 (80.0 mg, 238 μmol, 1.0 equiv) and S2 (50.7 mg, 285 μmol, 1.2 equiv) in DMF (2 mL) were added DIPEA (123 mg, 951 μmol, 4.0 equiv) and HATU (181 mg, 476 μmol, 2.0 equiv). The mixture was stirred at rt for 2 h. Upon reaction completion, the mixture was purified by reverse-phase chromatography (mobile phase: A: water (10 mM $NH_3$•$H_2O$), B: acetonitrile; gradient: 0% to 100% B over 20 min) to give S3 (100 mg, 89% yield) as a yellow solid.

To a precooled (0 ºC) solution of S3 (90.0 mg, 196 μmol, 1.0 equiv) and DIPEA (50.6 mg, 392 μmol, 2.0 equiv) in DCM (2 mL) was added acryloyl chloride (26.6 mg, 294 μmol, 1.5 equiv). The mixture was stirred at 0 °C for 1 h. Upon reaction completion, the mixture was concentrated under reduced pressure. The resulting residue was purified by prep-HPLC (mobile phase: A: water (10 mM $NH_4HCO_3$), B: acetonitrile; gradient: 38% to 68% B over 10 min) to give WX-02-322 (25.0 mg, 25% yield) as a white solid.
$^1$H NMR (400 MHz, $CD_3OD$) δ 7.53 (dt, *J* = 7.7, 1.1 Hz, 1H), 7.29 (d, *J* = 8.0 Hz, 1H), 7.24 – 7.06 (m, 2H), 7.03 (ddd, *J* = 8.1, 7.1, 1.1 Hz, 1H), 6.88 (s, 1H), 6.83 (d, *J* = 8.1 Hz, 1H), 6.75 (d, *J* = 8.0 Hz, 1H), 6.61 – 6.27 (m, 1.5H), 6.04 – 5.78 (m, 3.5H), 5.61 – 5.31 (m, 0.5H), 4.58 (br. s, 0.5H), 3.81 (t, *J* = 7.4 Hz, 2H), 3.53 – 3.32 (m, 4H), 3.23 – 3.04 (m, 2H), 2.98 (dd, *J* = 15.4, 6.1 Hz, 1H), 2.88 – 2.42 (m, 1H), 1.82 – 1.52 (m, 2H), 1.49 – 1.24 (m, 3H), 1.21 – 0.92 (m, 1H); mixture of rotamers; 1 exchangeable proton not observed.
HRMS m/z calc. for $C_{30}H_{32}N_3O_5$ $[M+H]^+$ 514.2342 found 514.2357.

1-((1*S*,3*S*)-1-(benzo[*d*][1,3]dioxol-5-yl)-3-(2-oxa-8-azaspiro[4.5]decane-8-carbonyl)-1,3,4,9-tetrahydro-2*H*-pyrido[3,4-*b*]indol-2-yl)prop-2-en-1-one (WX-02-304)
Prepared in similar fashion from *ent*-S1.
$^1$H NMR (400 MHz, $CD_3OD$) δ 7.53 (d, *J* = 7.8 Hz, 1H), 7.29 (d, *J* = 8.1 Hz, 1H), 7.24 – 7.07 (m, 2H), 7.06 – 6.98 (m, 1H), 6.88 (br.s , 1H), 6.83 (d, *J* = 8.2 Hz, 1H), 6.75 (d, *J* = 8.1 Hz, 1H), 6.65 – 6.29 (m, 1.5H), 6.03 – 5.77 (m, 3.5H), 5.65 – 5.30 (m, 0.5H), 4.74 – 4.45 (m, 0.5H), 3.81 (t, *J* = 7.2 Hz, 2H), 3.55 – 3.32 (m, 4H), 3.25 – 3.02 (m, 2H), 2.98 (dd, *J* = 15.5, 6.1 Hz, 1H), 2.89 – 2.37 (m, 1H), 1.80 – 1.51 (m, 2H), 1.49 – 1.22 (m, 3H), 1.19 – 0.85 (m, 1H); mixture of rotamers; 1 exchangeable proton not observed.
HRMS m/z calc. for $C_{30}H_{32}N_3O_5$ $[M+H]^+$ 514.2342 found 514.2338.

*Synthesis of WX-03-79*

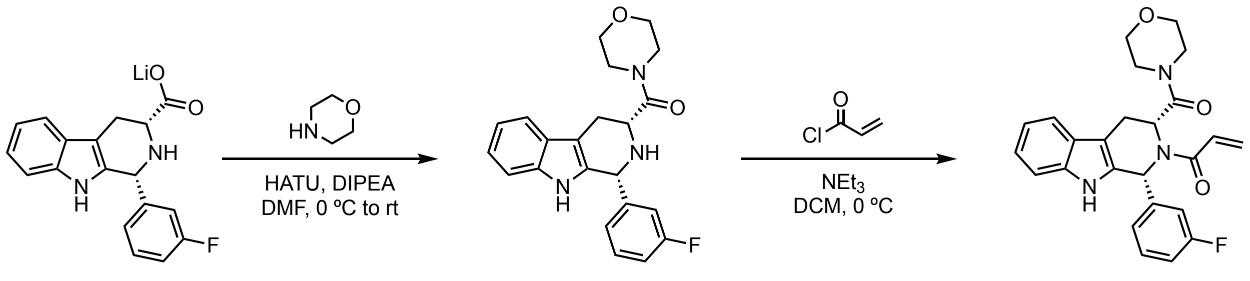


**S4-Li** **S5** **WX-03-79**

1-((1*R*,3*R*)-1-(3-fluorophenyl)-3-(morpholine-4-carbonyl)-1,3,4,9-tetrahydro-2*H*-pyrido[3,4-*b*]indol-2-yl)prop-2-en-1-one (WX-03-79)
To a precooled (0 ºC) solution of S4-Li (600 mg, 1.90 mmol, 1.0 equiv) and morpholine (826 mg, 9.49 mmol, 5.0 equiv) in DMF (7 mL) were added DIPEA (490 mg, 3.79 mmol, 2.0 equiv) and HATU (866 mg, 2.28 mmol, 1.2 equiv). The mixture was allowed to warm to rt and stirred for 1 h. Upon reaction completion, the reaction mixture was purified by reversed-phase HPLC (mobile phase: A: water (10 mM $NH_3$•$H_2O$), B: acetonitrile; gradient: 0% to 100% B over 20 min) to give S5 (512 mg, 71% yield) as a white solid.

To a precooled (0 ºC) solution of S5 (100 mg, 264 µmol, 1.0 equiv) and triethylamine (80 mg, 791 µmol, 3.0 equiv) in DCM (1 mL) was added dropwise a solution of acryloyl chloride (48 mg, 527 µmol, 2.0 equiv) in DCM (1 mL). The mixture was stirred at 0 °C for 1 h. Upon reaction completion, the solvent was removed under reduced pressure. The resulting residue was purified by prep-TLC ($SiO_2$, petroleum ether/EtOAc = 1:1) and prep-HPLC (mobile phase: A: $H_2O$ (10 mM $NH_4HCO_3$), B: acetonitrile; gradient: 28% to 58% B over 15 min) to give WX-03-79 (30 mg, 26% yield) as a white solid.
$^1$H NMR (400 MHz, $CD_3OD$) δ 7.54 (d, *J* = 7.9 Hz, 1H), 7.45 – 7.20 (m, 3H), 7.08 (m, 5.5H), 6.69 – 6.50 (m, 0.5H), 6.41 (d, *J* = 16.5 Hz, 1H), 5.94 (d, *J* = 10.6 Hz, 1H), 5.88 – 5.31 (m, 1H), 3.72 – 3.33 (m, 6H), 3.22 – 2.73 (m, 3H), 2.60 – 1.94 (m, 1H); mixture of rotamers; 1 exchangeable proton not observed.
HRMS m/z calc. for $C_{25}H_{25}FN_3O_3$ $[M+H]^+$ 434.188 found 434.1881.

1-((1*S*,3*S*)-1-(3-fluorophenyl)-3-(morpholine-4-carbonyl)-1,3,4,9-tetrahydro-2*H*-pyrido[3,4-*b*]indol-2-yl)prop-2-en-1-one (WX-03-77)
Prepared in similar fashion from *ent*-S4-Li.
$^1$H NMR (400 MHz, $CD_3OD$) δ 7.54 (dd, *J* = 7.9, 1.0 Hz, 1H), 7.46 – 7.21 (m, 3H), 7.19 – 6.94 (m, 5.5H), 6.69 – 6.49 (m, 0.5H), 6.41 (d, *J* = 16.6 Hz, 1H), 5.94 (d, *J* = 10.6 Hz, 1H), 5.86 – 5.44 (m, 1H), 3.69 – 3.33 (m, 6H), 3.24 – 2.75 (m, 3H), 2.63 – 1.98 (m, 1H); mixture of rotamers; 1 exchangeable proton not observed.
HRMS m/z calc. for $C_{25}H_{25}FN_3O_3$ $[M+H]^+$ 434.188 found 434.1886.

*Synthesis of WX-02-760 (HL-1B)*

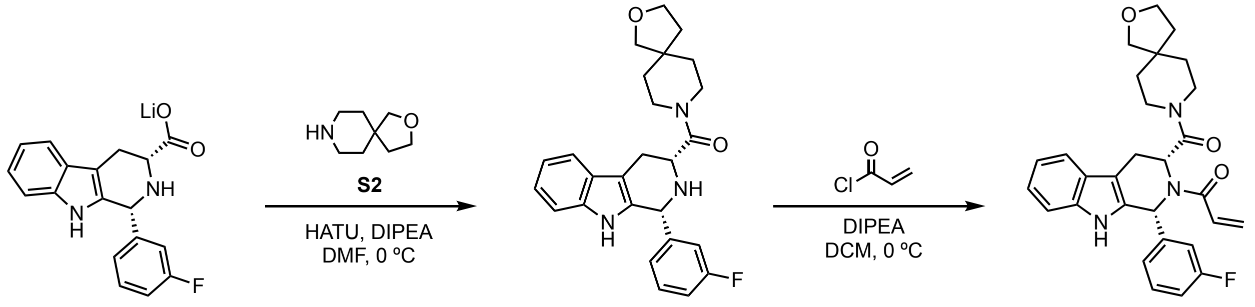


**S4-Li** **S6** **WX-02-760**

1-((1*R*,3*R*)-1-(3-fluorophenyl)-3-(2-oxa-8-azaspiro[4.5]decane-8-carbonyl)-1,3,4,9-tetrahydro-2*H*-pyrido[3,4-*b*]indol-2-yl)prop-2-en-1-one (WX-02-760=HL-1B)
To a precooled (0 ºC) solution of S4-Li (400 mg, 1.26 mmol, 1.0 equiv) and S2 (449 mg, 2.53 mmol, 2.0 equiv) in DMF (3 mL) were added HATU (962 mg, 2.53 mmol, 2.0 equiv) and DIPEA (490 mg, 3.79 mmol, 3.0 equiv). The mixture was stirred at 0 °C for 2 h. Upon reaction completion, the mixture was concentrated under reduced pressure. The resulting residue was purified by reverse-phase column chromatography (A: water (10 mM HCl), B: acetonitrile; gradient: 0% to 100% B over 20 min) to give S6 (500 mg, 90% yield) as a white solid.
To a precooled (0 ºC) solution of S6 (500 mg, 1.15 mmol, 1.0 equiv) and DIPEA (447 mg, 3.46 mmol, 3.0 equiv) in DCM (3 mL) was added acryloyl chloride (157 mg, 1.73 mmol, 1.5 equiv). The mixture was stirred at 0 °C for 0.5 h. Upon reaction completion, the mixture was concentrated

under reduced pressure. The resulting residue was purified by column chromatography ($SiO_2$, petroleum ether/EtOAc = 2:1) and prep-HPLC (mobile phase: A: water (10 mM $NH_4HCO_3$), B: acetonitrile; gradient: 38% to 68% B over 9 min) to give HL-1B (207 mg, 37% yield) as a white solid.

$^1$H NMR (400 MHz, $CD_3OD$) δ 7.54 (d, *J* = 7.8 Hz, 1H), 7.45 – 7.28 (m, 2H), 7.25 (br. s, 1H), 7.20 – 7.08 (m, 3H), 7.08 – 6.96 (m, 2H), 6.69 – 6.49 (m, 1H), 6.41 (br. d, *J* = 16.6 Hz, 1H), 5.94 (d, *J* = 10.6 Hz, 1H), 5.89 – 5.36 (m, 1H), 3.87 – 3.74 (m, 2H), 3.59 – 3.33 (m, 4H), 3.27 – 2.81 (m, 3H), 2.74 – 2.04 (m, 1H), 1.81 – 1.52 (m, 2H), 1.52 – 1.20 (m, 3H), 1.23 – 0.71 (m, 1H); mixture of rotamers; 1 exchangeable proton not observed.

HRMS m/z calc. for $C_{29}H_{31}FN_3O_3$ $[M+H]^+$ 488.2349 found 488.2348.

1-((1*S*,3*S*)-1-(3-fluorophenyl)-3-(2-oxa-8-azaspiro[4.5]decane-8-carbonyl)-1,3,4,9-tetrahydro-2*H*-pyrido[3,4-*b*]indol-2-yl)prop-2-en-1-one (WX-02-759=HL-1A)

Prepared in similar fashion from *ent*-S4-Li.

$^1$H NMR (400 MHz, $CD_3OD$) δ 7.54 (d, *J* = 7.8 Hz, 1H), 7.42 – 7.28 (m, 2H), 7.29 – 7.21 (m, 1H), 7.21 – 7.08 (m, 3H), 7.08 – 6.98 (m, 2H), 6.69 – 6.49 (m, 1H), 6.49 – 6.34 (m, 1H), 5.95 (br. d, *J* = 10.5 Hz, 1H), 5.89 – 5.35 (m, 1H), 3.90 – 3.70 (m, 2H), 3.63 – 3.34 (m, 4H), 3.28 – 2.78 (m, 3H), 2.76 – 2.07 (m, 1H), 1.83 – 1.53 (m, 2H), 1.52 – 1.22 (m, 3H), 1.22 – 0.77 (m, 1H); mixture of rotamers; 1 exchangeable proton not observed.

HRMS m/z calc. for $C_{29}H_{31}FN_3O_3$ $[M+H]^+$ 488.2349 found 488.2341.

*Synthesis of WX-02-964 (HL-1B-yne)*

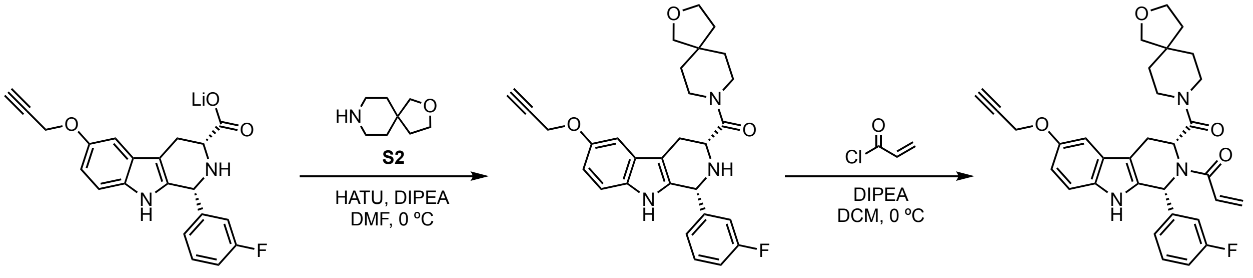


**S7-Li** **S8** **WX-02-964**

1-((1*R*,3*R*)-1-(3-fluorophenyl)-6-(prop-2-yn-1-yloxy)-3-(2-oxa-8-azaspiro[4.5]decane-8-carbonyl)-1,3,4,9-tetrahydro-2*H*-pyrido[3,4-*b*]indol-2-yl)prop-2-en-1-one (WX-02-964=HL-1B-yne)

To a precooled (0 ºC) solution of S7-Li (150 mg, 405 µmol, 1.0 equiv) and S2 (93.6 mg, 527 µmol, 1.3 equiv) in DMF (2 mL) were added DIPEA (209 mg, 1.62 mmol, 4.0 equiv) and HATU (308 mg, 810 µmol, 2.0 equiv). The mixture was stirred at 0 °C for 2 h. Upon reaction completion, the mixture was diluted with water (30 mL) and extracted with EtOAc (20 mL × 3). The combined organic layers were washed with brine (20 mL × 3), dried over Na2SO4, filtered and concentrated under reduced pressure to give a residue. The residue was purified by reverse-phase column chromatography (A: water (10mM HCl), B: acetonitrile; gradient: 0% to 100% B over 20 min) to obtain S8 (100 mg, 95% purity [LC-MS], 48 % yield) as a yellow solid.

To a precooled (0 ºC) solution of S8 (100 mg, 95% purity [LC-MS], 195 µmol, 1.0 equiv) and DIPEA (106 mg, 820 µmol, 4.2 equiv) in DCM (2 mL) was added acryloyl chloride (27.9 mg, 308 µmol, 1.58 equiv). The mixture was stirred at 0 °C for 0.5 h. Upon reaction completion, the mixture was concentrated under reduced pressure. The resulting residue was purified by prep-TLC ($SiO_2$, petroleum ether/EtOAc = 1:2) and prep-HPLC (mobile phase: A: $H_2O$ (10mM $NH_4HCO_3$), B: acetonitrile; gradient: 32% to 62% B over 15 min) to obtain HL-1B-yne (42 mg, 40% yield) as a white solid.
$^1$H NMR (400 MHz, $CDCl_3$) δ 7.78 (br. s, 1H), 7.33 – 7.22 (m, 1H), 7.21 – 7.06 (m, 4.5H), 7.01 – 6.94 (m, 1H), 7.01 – 6.79 (m, 1.5H), 6.91 (dd, *J* = 8.8, 2.5 Hz, 1H), 6.47 (d, *J* = 16.6 Hz, 1H), 6.43 – 6.09 (m, 1H), 5.89 (dd, *J* = 10.6, 1.6 Hz, 1H), 4.73 (d, *J* = 2.4 Hz, 2H), 3.79 (q, *J* = 6.7 Hz, 2H), 3.53 – 3.18 (m, 5H), 2.98 (ddd, *J* = 15.4, 6.2, 1.5 Hz, 1H), 2.98 – 2.75 (m, 1H), 2.51 (t, *J* = 2.4 Hz, 1H), 2.39 – 2.28 (m, 0.5H), 2.27 – 2.14 (m, 0.5H), 1.69 – 1.63 (m, 1H), 1.58 – 1.46 (m, 1H), 1.44 – 1.21 (m, 3H), 1.06 (m, 1H); mixture of rotamers.
HRMS m/z calc. for $C_{32}H_{33}FN_3O_4$ $[M+H]^+$ 542.2455 found 542.2457.

1-((1*S*,3*S*)-1-(3-fluorophenyl)-6-(prop-2-yn-1-yloxy)-3-(2-oxa-8-azaspiro[4.5]decane-8-carbonyl)-1,3,4,9-tetrahydro-2*H*-pyrido[3,4-*b*]indol-2-yl)prop-2-en-1-one (WX-02-963=HL-1A-yne)
Prepared in similar fashion from *ent*-S7-Li.
$^1$H NMR (400 MHz, $CDCl_3$) δ 7.71 (br. s, 1H), 7.32 – 7.26 (m, 1H), 7.23 – 7.06 (m, 4.5H), 6.98 (t, *J* = 8.4 Hz, 1H), 6.94 – 6.90 (m, 1H), 7.01 – 6.79 (m, 1.5H), 6.47 (d, *J* = 16.6 Hz, 1H), 6.44 – 6.19 (m, 1H), 5.90 (d, *J* = 10.6 Hz, 1H), 4.73 (d, *J* = 2.4 Hz, 2H), 3.80 (q, *J* = 6.7 Hz, 2H), 3.58 – 3.23 (m, 5H), 2.98 (dd, *J* = 15.4, 6.1 Hz, 1H), 2.94 – 2.78 (m, 1H), 2.51 (t, *J* = 2.4 Hz, 1H), 2.40 – 2.29 (m, 0.5H), 2.26 – 2.12 (m, 0.5H), 1.72 – 1.59 (m, 1H), 1.58 – 1.46 (m, 1H), 1.44 – 1.18 (m, 3H), 1.16 – 0.92 (m, 1H); mixture of rotamers.
HRMS m/z calc. for $C_{32}H_{33}FN_3O_4$ $[M+H]^+$ 542.2455 found 542.2455.

### Analytical data: NMR spectra

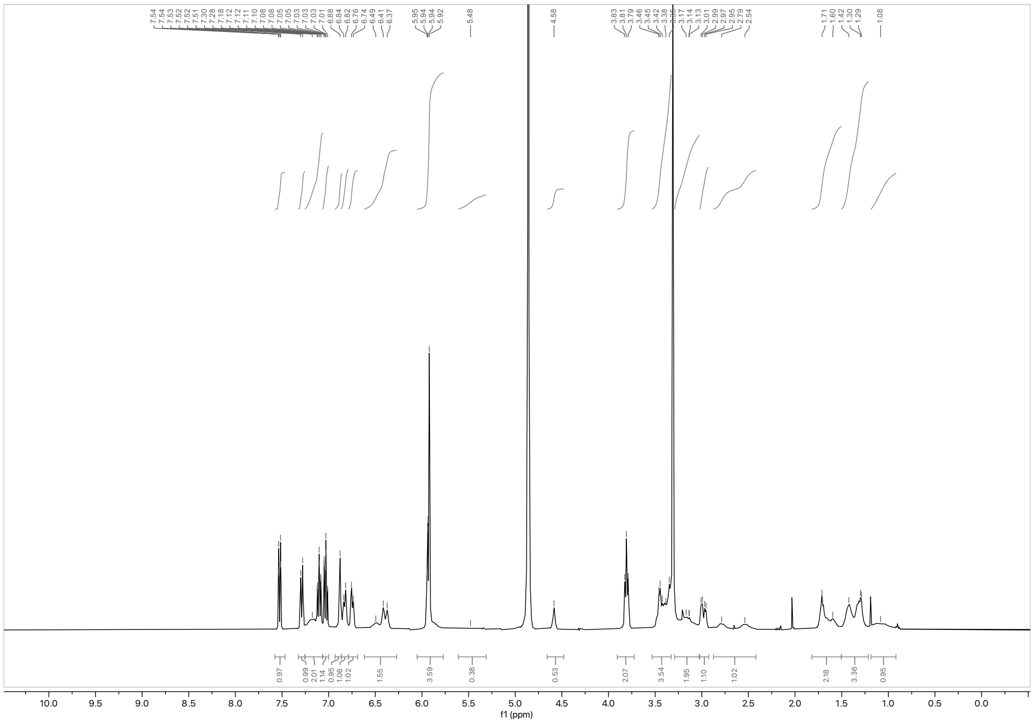

$^1H$ NMR spectrum of WX-02-322 (400 MHz, $CD_3OD$)

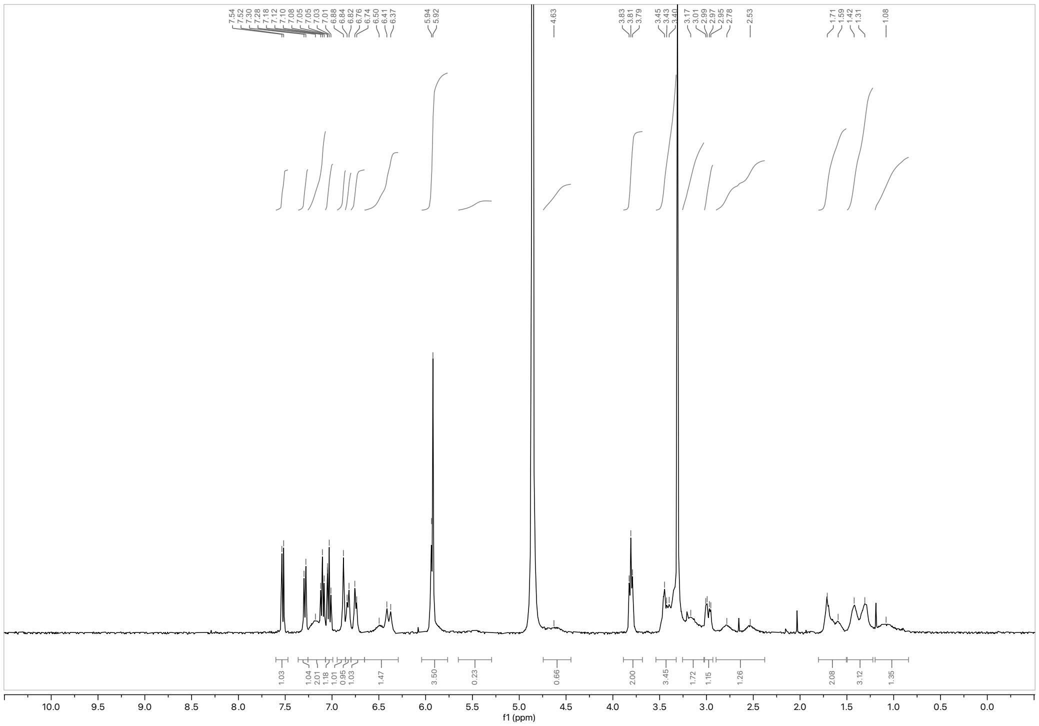

$^1H$ NMR spectrum of WX-02-304 (400 MHz, $CD_3OD$)

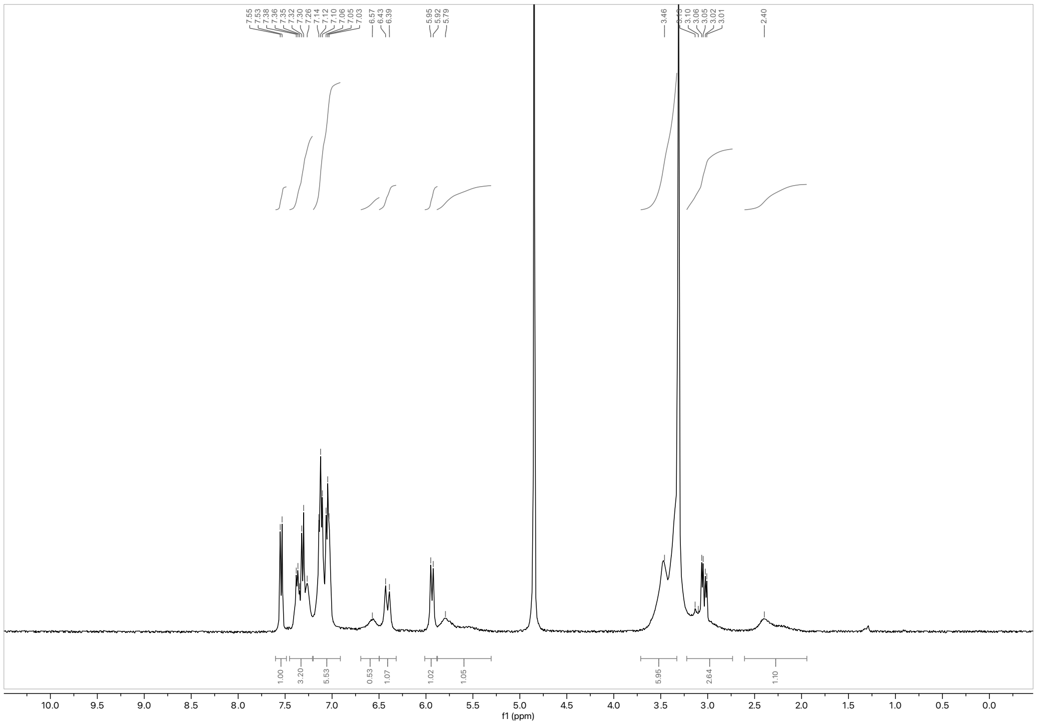

[1]H NMR spectrum of WX-03-79 (400 MHz, $CD_3OD$)

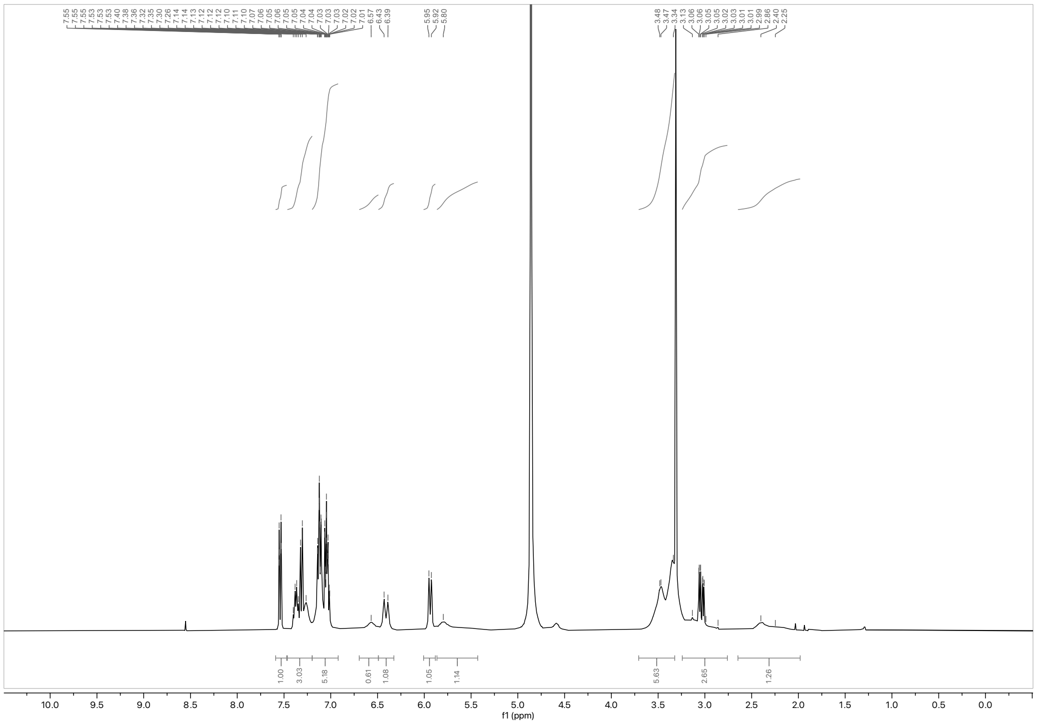

[1]H NMR spectrum of WX-03-77 (400 MHz, $CD_3OD$)

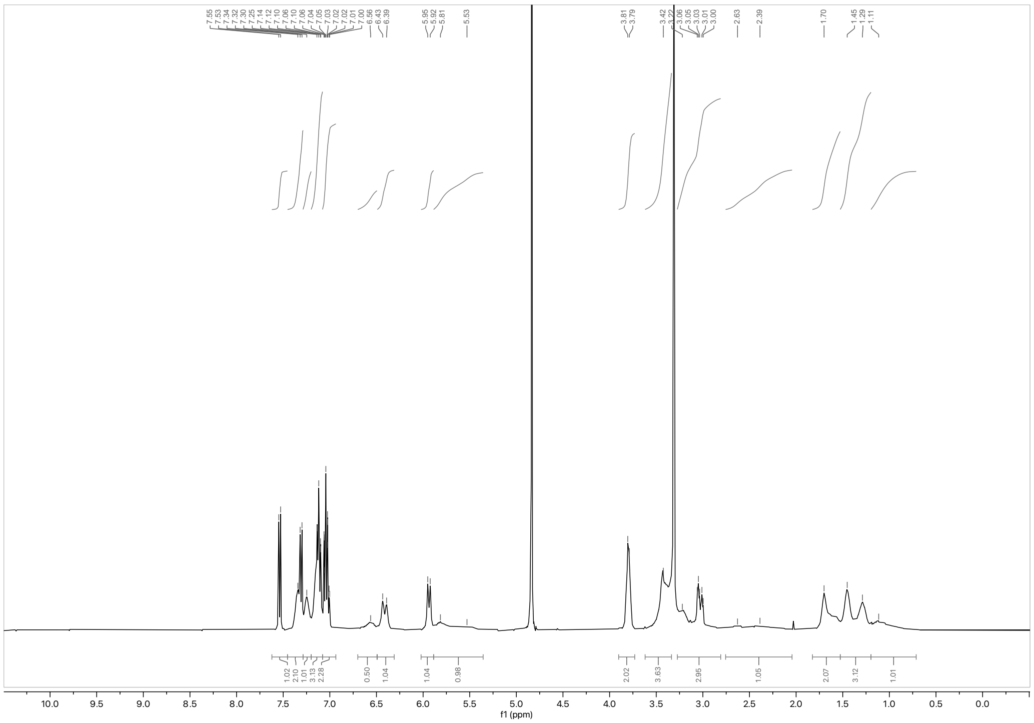

[1]H NMR spectrum of WX-02-760 (HL-1B) (400 MHz, $CD_3OD$)

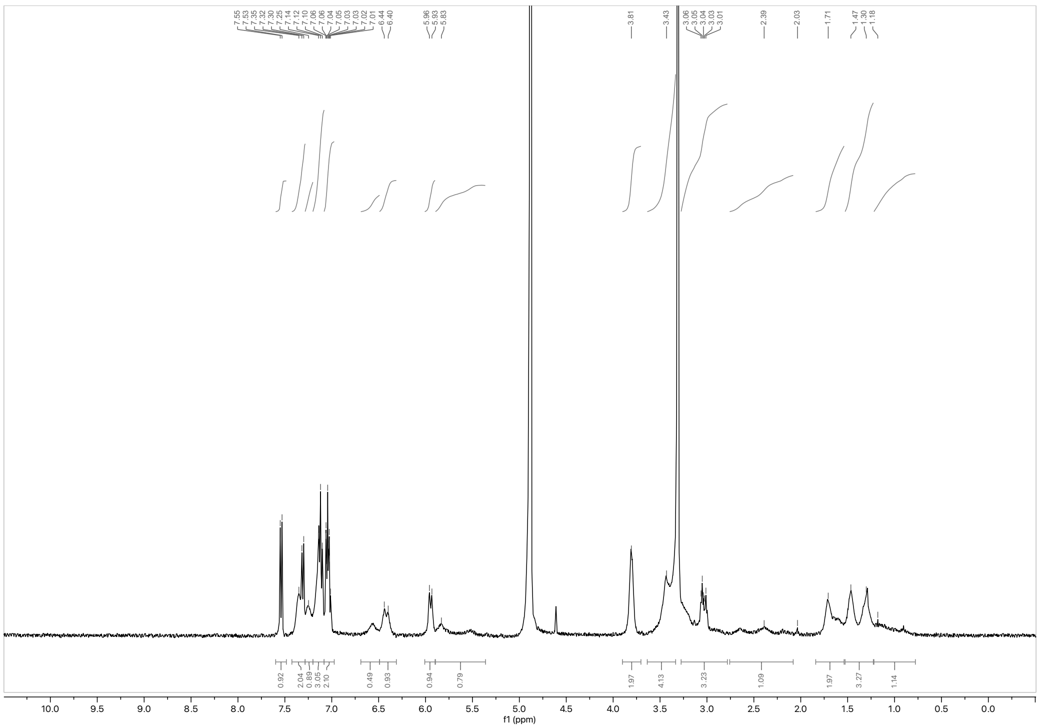

[1]H NMR spectrum of WX-02-759 (HL-1A) (400 MHz, $CD_3OD$)

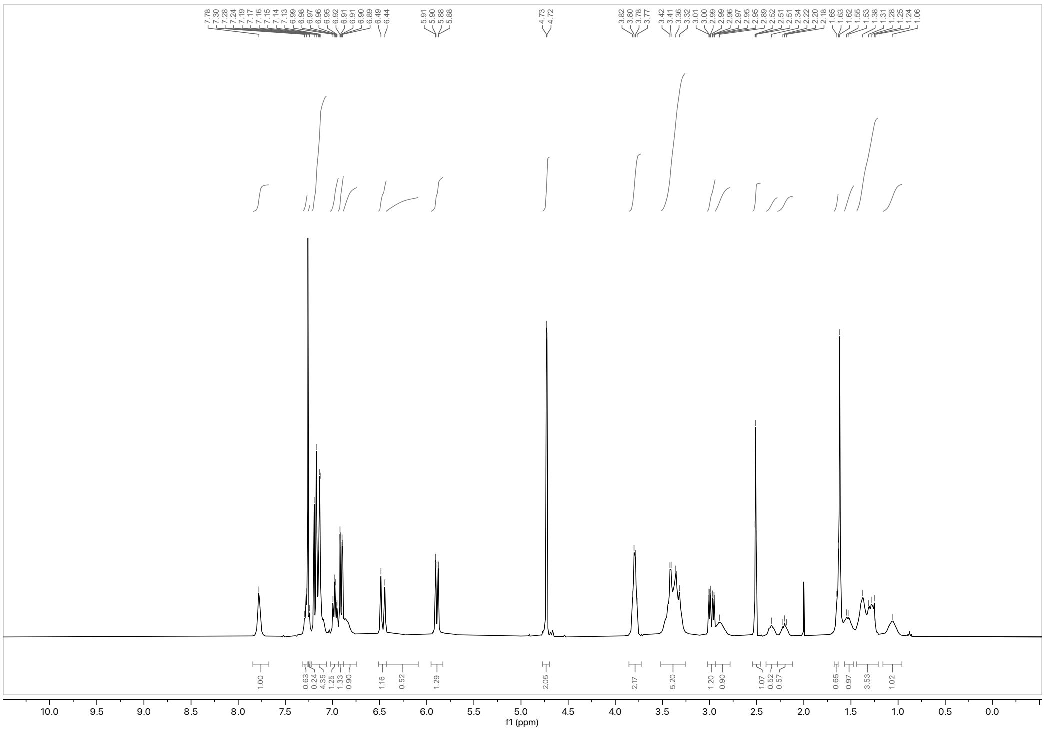


$^{1}H$ NMR spectrum of WX-02-964 (HL-1B-yne) (400 MHz, $CDCl_3$)

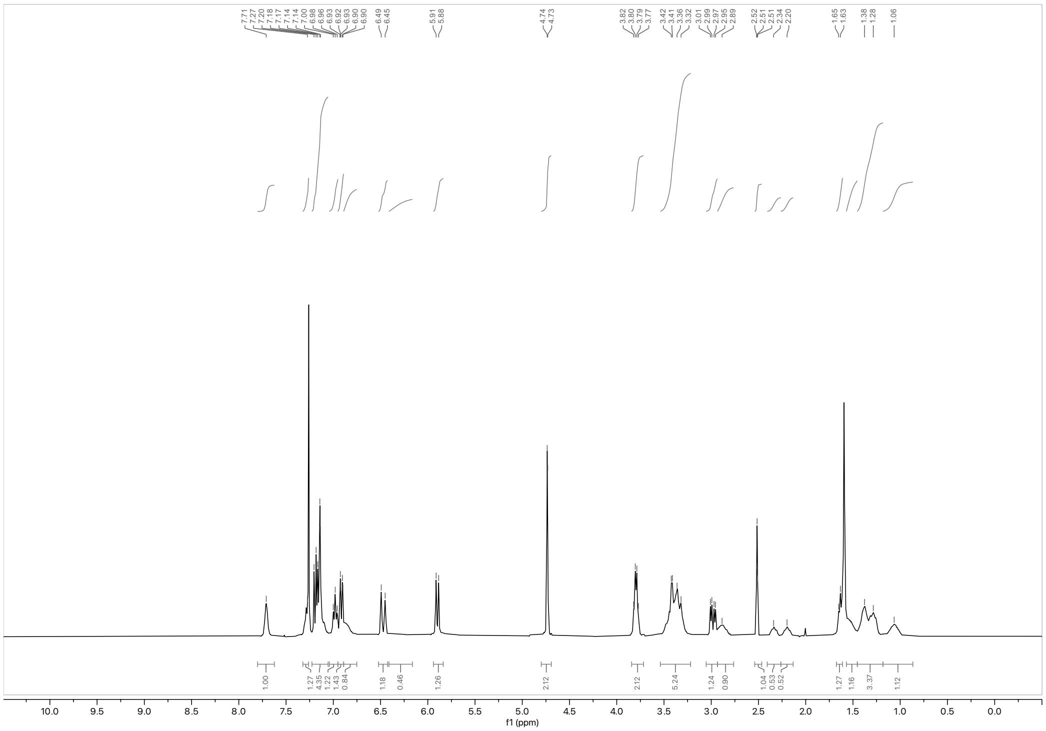


$^{1}H$ NMR spectrum of WX-02-963 (HL-1A-yne) (400 MHz, $CDCl_3$)

**Analytical data: SFC**

WX-02-322

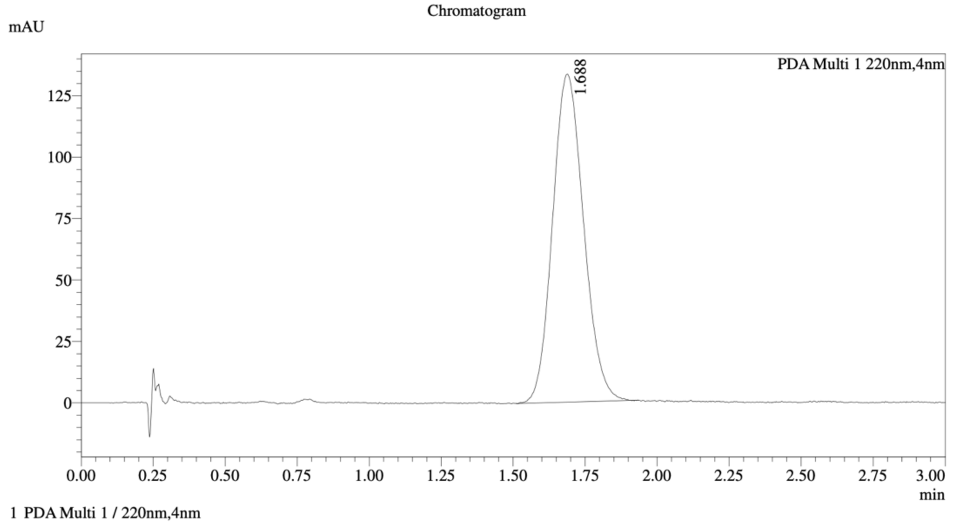


Integration Result

Peak Table

PDA Ch1 220nm

| Peak# | Ret. Time | Height | Height% | Resolution(USP) | Area | Area% |
|---|---|---|---|---|---|---|
| 1 | 1.688 | 133158 | 100.000 | -- | 1000330 | 100.000 |

WX-02-304

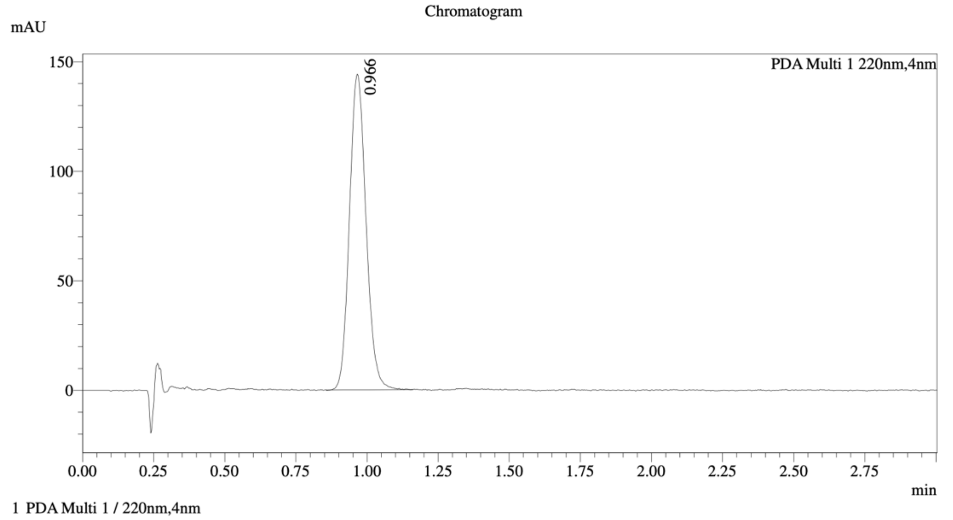


Integration Result

Peak Table

PDA Ch1 220nm

| Peak# | Ret. Time | Height | Height% | Resolution(USP) | Area | Area% |
|---|---|---|---|---|---|---|
| 1 | 0.966 | 143630 | 100.000 | -- | 584307 | 100.000 |

Mixture of WX-02-322 & WX-02-304

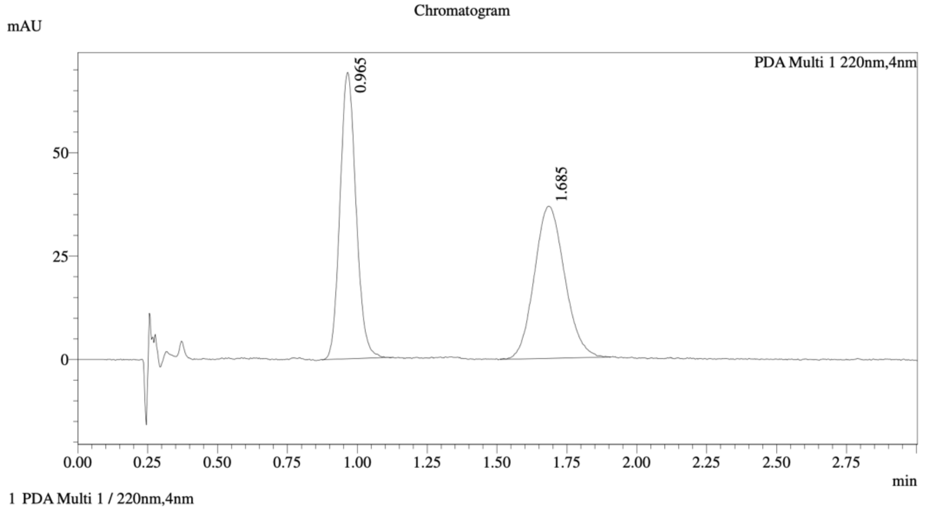


================================================================================

Integration Result

================================================================================

Peak Table

PDA Ch1 220nm

| Peak# | Ret. Time | Height | Height% | Resolution(USP) | Area | Area% |
|---|---|---|---|---|---|---|
| 1 | 0.965 | 69046 | 65.271 | -- | 276039 | 49.840 |
| 2 | 1.685 | 36737 | 34.729 | 4.710 | 277816 | 50.160 |

Method details

Column: Chiralcel OD-3 50×4.6mm I.D., 3 μm
Mobile phase: A: $CO_2$, B: MeOH (0.05% diethylamine)
Elution: 40% B

WX-03-79

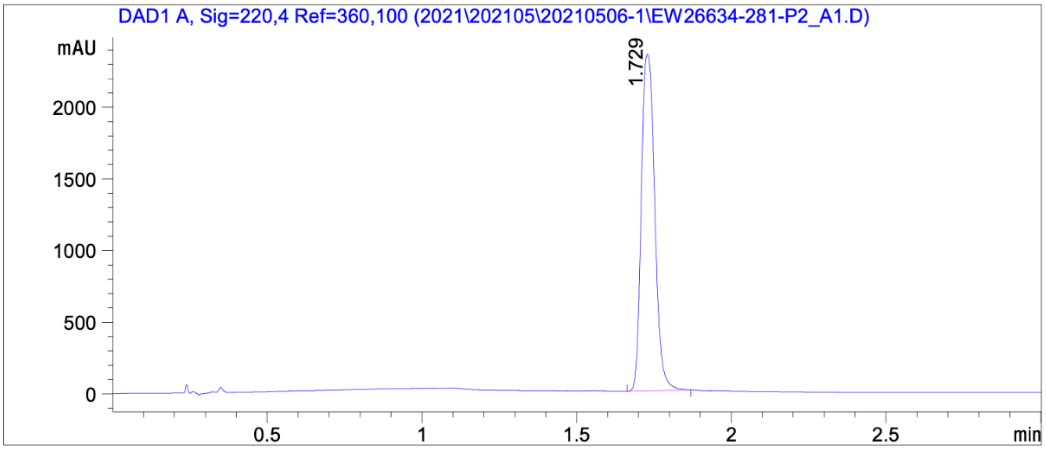


Signal: DAD1 A, Sig=220,4 Ref=360,100

| RetTime [min] | Height | Resolution | Symm. | Area [mAU*s] | Area % |
|---|---|---|---|---|---|
| 1.729 | 2355.278 | | 0.908 | 7092.230 | 100.000 |

WX-03-77

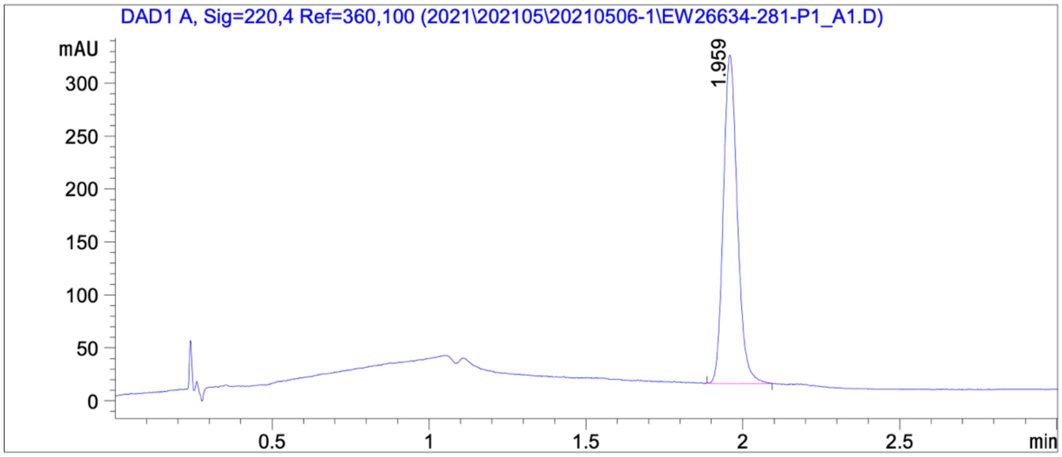


Signal: DAD1 A, Sig=220,4 Ref=360,100

| RetTime [min] | Height | Resolution | Symm. | Area [mAU*s] | Area % |
|---|---|---|---|---|---|
| 1.959 | 311.343 | | 0.901 | 962.277 | 100.000 |

Mixture of WX-03-79 & WX-03-77

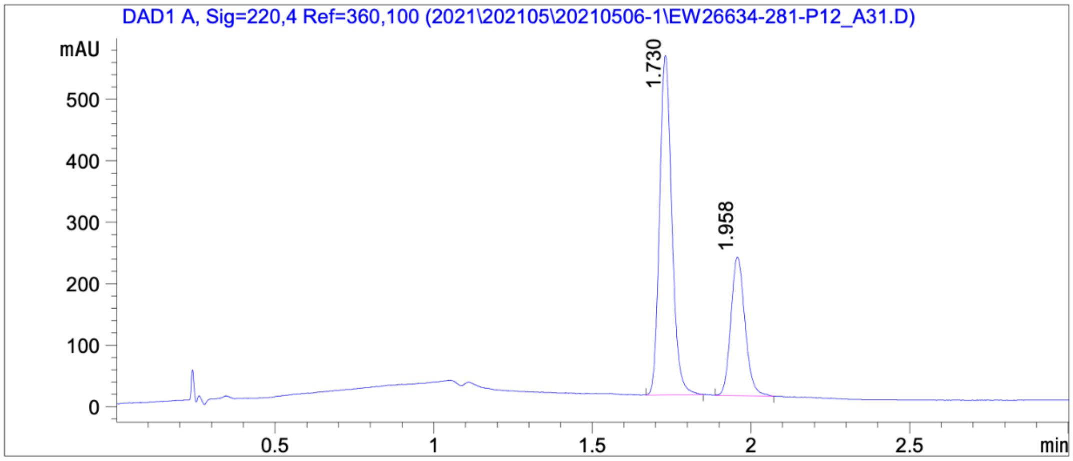


Signal: DAD1 A, Sig=220,4 Ref=360,100

| RetTime [min] | Height | Resolution | Symm. | Area [mAU*s] | Area % |
|---|---|---|---|---|---|
| 1.730 | 555.497 | | 0.749 | 1485.000 | 67.883 |
| 1.958 | 226.676 | 3.010 | 0.689 | 702.581 | 32.117 |

Method details

Column: Chiralpak IC-3 50×4.6mm I.D., 3 μm
Mobile phase: A: $CO_2$, B: MeOH (0.05% diethylamine)
Elution: 5% to 40% B (gradient)

WX-02-760 (HL-1B)

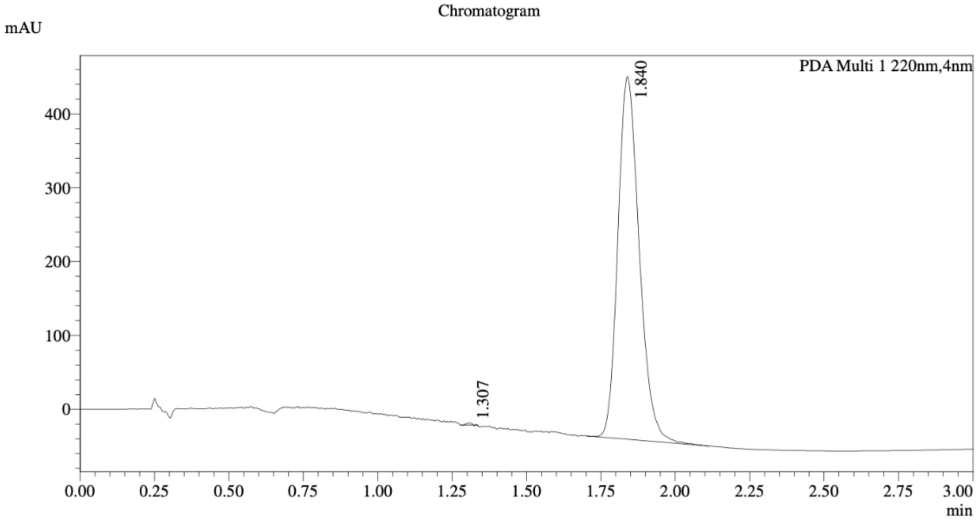


1 PDA Multi 1 / 220nm,4nm

Integration Result

Peak Table

PDA Ch1 220nm

| Peak# | Ret. Time | Height | Height% | Resolution(USP) | Area | Area% |
|---|---|---|---|---|---|---|
| 1 | 1.307 | 2631 | 0.535 | -- | 3284 | 0.134 |
| 2 | 1.840 | 488809 | 99.465 | 5.778 | 2456157 | 99.866 |

WX-02-759 (HL-1A)

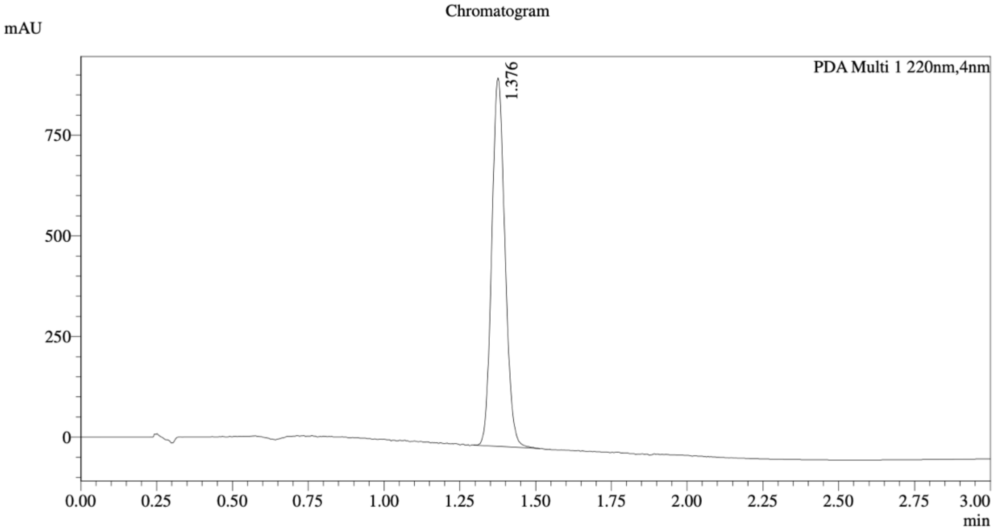


1 PDA Multi 1 / 220nm,4nm

Integration Result

Peak Table

PDA Ch1 220nm

| Peak# | Ret. Time | Height | Height% | Resolution(USP) | Area | Area% |
|---|---|---|---|---|---|---|
| 1 | 1.376 | 902971 | 100.000 | -- | 2772843 | 100.000 |

Mixture of WX-02-760 (HL-1B) & WX-02-759 (HL-1A)

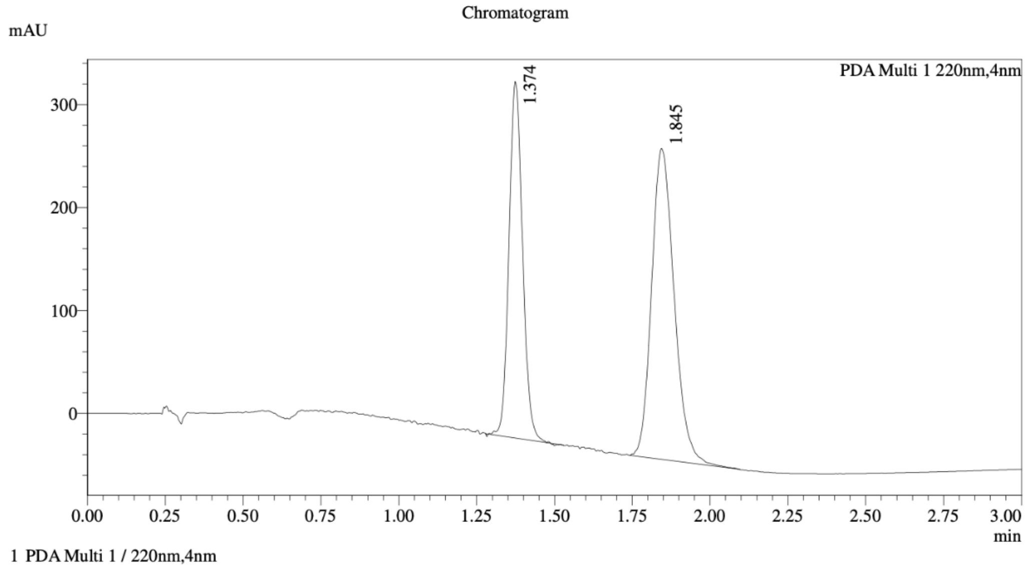


Integration Result

Peak Table

PDA Ch1 220nm

| Peak# | Ret. Time | Height | Height% | Resolution(USP) | Area | Area% |
|---|---|---|---|---|---|---|
| 1 | 1.374 | 343886 | 53.316 | -- | 1049015 | 40.835 |
| 2 | 1.845 | 301108 | 46.684 | 4.362 | 1519886 | 59.165 |

Method details

Column: Chiralpak IC-3 50×4.6mm I.D., 3 μm
Mobile phase: A: $CO_2$, B: EtOH (0.05% diethylamine)
Elution: 5% to 25% B (gradient)

WX-02-964 (HL-1B-yne)

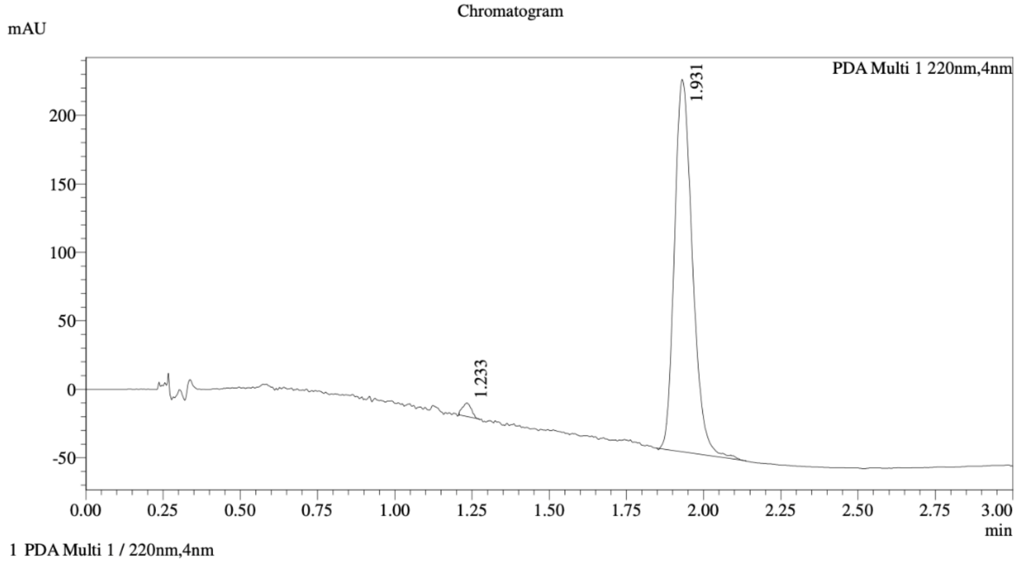


Integration Result

Peak Table

PDA Ch1 220nm

| Peak# | Ret. Time | Height | Height% | Resolution(USP) | Area | Area% |
|---|---|---|---|---|---|---|
| 1 | 1.233 | 9367 | 3.351 | -- | 18926 | 1.691 |
| 2 | 1.931 | 270155 | 96.649 | 8.279 | 1100298 | 98.309 |

WX-02-963 (HL-1A-yne)

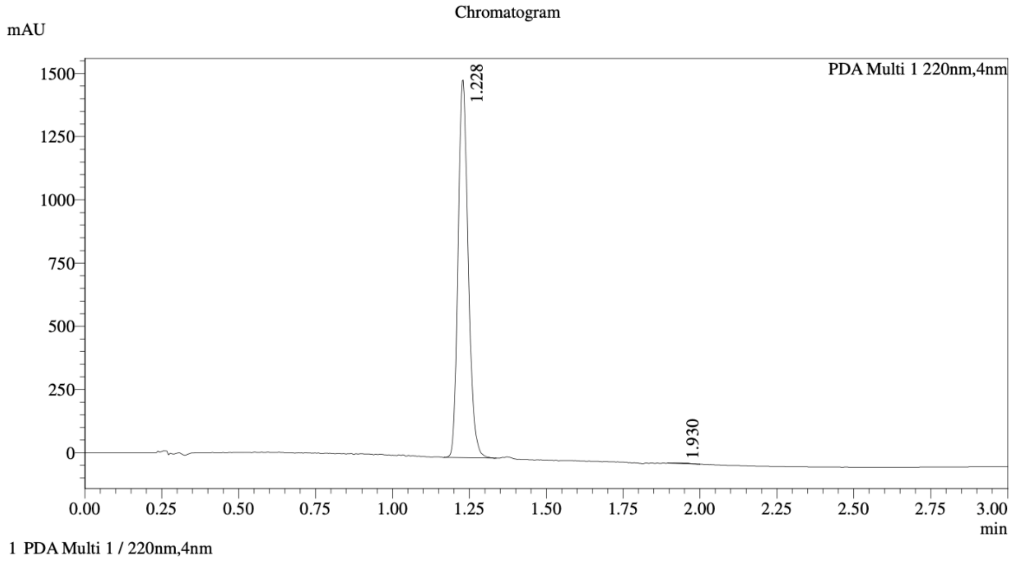


Integration Result

Peak Table

PDA Ch1 220nm

| Peak# | Ret. Time | Height | Height% | Resolution(USP) | Area | Area% |
|---|---|---|---|---|---|---|
| 1 | 1.228 | 1477287 | 99.823 | -- | 3426117 | 99.732 |
| 2 | 1.930 | 2623 | 0.177 | 9.161 | 9190 | 0.268 |

Mixture of WX-02-964 (HL-1B-yne) & WX-02-963 (HL-1A-yne)

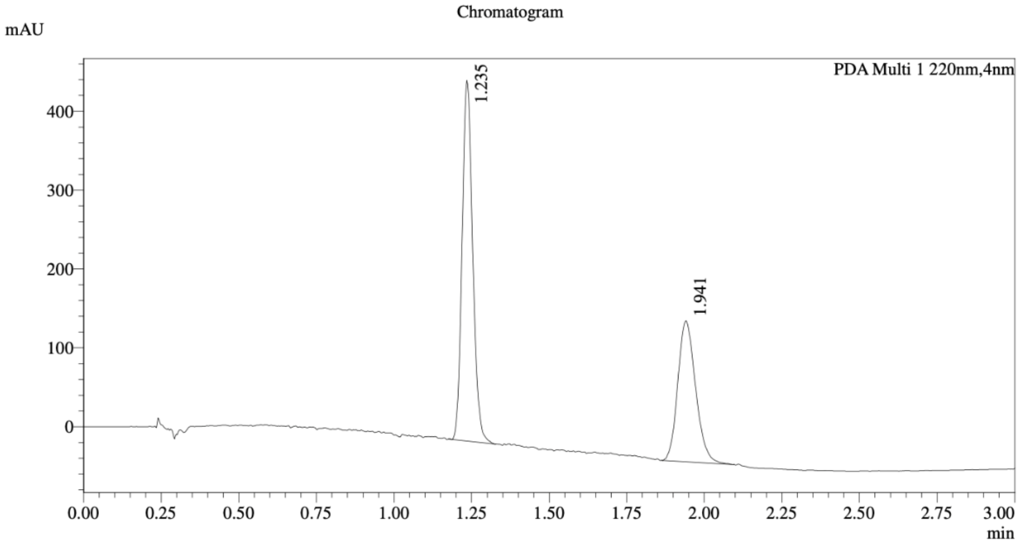


1 PDA Multi 1 / 220nm,4nm

Integration Result

Peak Table

PDA Ch1 220nm

| Peak# | Ret. Time | Height | Height% | Resolution(USP) | Area | Area% |
|---|---|---|---|---|---|---|
| 1 | 1.235 | 448169 | 71.576 | -- | 1083730 | 60.312 |
| 2 | 1.941 | 177975 | 28.424 | 8.173 | 713155 | 39.688 |

Method details

Column: Chiralcel OD-3 50×4.6mm I.D., 3 µm
Mobile phase: A: $CO_2$, B: EtOH (0.05% diethylamine)
Elution: 20% to 60% B (gradient)
